# “Gut microbial communities of velvet worm *Euperipatoides rowelli* (Onychophora) across deadwood microhabitats in southeastern Australia”

**DOI:** 10.1101/2025.10.21.683314

**Authors:** Imelda L. Forteza, Thomas Wong, David Rowell, Teng Li, Francis Tablizo, Ulrike Mathesius, Allen Rodrigo

**Affiliations:** Research School of Biology, Australian National University, 134 Linnaeus Way, Canberra ACT 2601, Australia; The School of Biological Sciences, The University of Auckland, Private Bag 92019, Auckland, New Zealand; Core Facility for Bioinformatics, Philippine Genome Center, University of the Philippines System

**Keywords:** Velvet worm, gut microbiome, saproxylic invertebrates, *Euperipatoides rowelli*, deadwood microhabitats, environmental filtering, 16S rRNA gene, historical refugia, Australia

## Abstract

This study investigated the gut microbial community of the velvet worm *Euperipatoides rowelli*, a saproxylic invertebrate residing in deadwood microhabitats within historical refugia in the Tallaganda forest, New South Wales, Australia. The eucalyptus-dominated temperate forest is marked by unique topography, hydrology, and historical influences from Pleistocene glacial cycles.

We described patterns in gut microbial community structure and composition across eight sites. The velvet worms were unfed for two weeks prior to dissection to lessen the influence of transient prey-associated microorganisms. Amplicon sequencing of the V4-V5 region of the 16S rRNA gene was sequenced using the Illumina MiSeq platform, with positive and negative controls integrated to assess contamination and ensure data quality.

Alpha and beta diversity metrics demonstrated minimal overall variation among sites, suggesting broadly similar community structure. However, a closer look at individual taxa revealed contrasting patterns. *Proteobacteria* dominated the gut microbiome, with genus *A37b* (*Rickettsia*) forming part of the core microbiome in 50% of individuals; its presence varied among sites. In contrast, *Spiroplasma* displayed substantial site-specific variation in relative abundance. These findings suggest that, while overall diversity remained relatively stable, particular taxa displayed heterogenous distributions across historical refugia. Collectively, our findings suggest that gut microbiomes of velvet worms are shaped more by environmental filtering within deadwood microhabitats than by host geographic differences, reflecting a combination of stochastic and niche-driven assembly processes.

**Impact Statement:** We present the first characterization of the gut microbiome in the saproxylic velvet worm *Euperipatoides rowelli* from temperate Australian forests. Despite marked host genetic divergence, microbial communities showed minimal spatial structuring, suggesting substantial environmental filtering within deadwood microhabitats. The limited variation and transient taxa may indicate microbial associations are more likely environmentally acquired rather than host-specific. These findings broaden our understanding of the microbiome assembly in underexplored invertebrates and underscore the influence of ecological context over site-based spatial differentiation in shaping gut microbial communities.

## INTRODUCTION

Microbial communities play critical roles in shaping ecosystem resilience and function [1][2]. A central question in microbial ecology is the extent to which host-microorganism associations reflect host traits, evolutionary history, or environmental context. Resolving this requires evaluating the factors that influence both alpha and beta diversity within and across hosts [3][ 4].

We focus on the gut microbiome of *Euperipatoides rowelli*, a velvet worm (Onychophora) inhabiting deadwood microhabitats across five historical refugia—Harold’s Cross, Eastern Slope Region, Anembo, Pike’s Saddle, and Badja Region—in the Tallaganda region in New South Wales, Australia [5][6]. These eucalyptus-dominated temperate forests, shaped by the Pleistocene glacial-interglacial cycle and complex topography, have served as long-term refugia for saproxylic fauna [7].

*E. rowelli* are soft-bodied, low-dispersal generalist predators with limited gut compartmentalization and carry out enzymatic extra-oral digestion (Manton & Heatley, 1937; [8]. These attributes, coupled with reliance on damp deadwood habitats and irregular feeding, may limit gut microbial diversity and promote environmental acquisition of microorganisms [9]. While social behaviours such as clustering, dominance hierarchies, and cooperative feeding have been documented in *E. rowelli*, their potential influence on microbiome assembly is analogous to trophallaxis and coprophagy in eusocial insects [10][11] and remains undocumented. These ecological and behavioural features raise questions about how host biology and environment shape its gut microbial community. Additionally, we used relative abundance (with appropriate statistical transformations) to capture the inherent compositional structure of microbiomes sampled; neglecting this could obscure genuine biological signals [12].

Despite increased interest in invertebrate microbiomes, the Onychophora remains a poorly investigated phylum. This study addresses a knowledge gap in early-diverging panarthropods by providing the first sequencing-based characterization of the gut microbiome in *E. rowelli*. Our study further asks whether its composition reflects (i) spatial divergence across refugia or (ii) environmental filtering within deadwood microhabitats. Given the long-term geographic isolation of host populations, we hypothesized that gut microbial composition would exhibit spatial structuring corresponding to refugial distribution.

## METHODS

### Sample Collection

From 2018 to 2021, *E. rowelli* were collected from decomposing logs across eight sites (Table 1) within historical refugia distributed along a longitudinal forest gradient in Tallaganda, NSW, under New South Wales Scientific Licence SL102322, issued by the Department of Planning, Industry and Environment under the Biodiversity Conservation Act 2016 (see in Supplementary Permit SL102322). Logs were opened using hand tools to access specimens. Sites 1 and 2 experienced bushfire impact in 2019 but showed signs of post-disturbance recovery by 2020. Specimens were stored with moist wood and frass at 4°C for up to two weeks. This study characterizes the velvet worm gut microbiome for the first time (n = 64). A summary of the processed samples is shown in Supplementary Table S1.

**Table 1.** Description of the Velvet Worm Sampling Sites. Summary of *E. rowelli* sampling sites across the Tallaganda region, including geographic coordinates, associated refugia, and sample counts retained for microbial community analysis.

| Velvet Worms Sampling Sites |  |  |  |  |  |
| --- | --- | --- | --- | --- | --- |
| Localities and Number of Velvet Worms Sampled per Site |  |  |  |  |  |
| Site | Gradient | Locality | Latitude | Longitude | Number of Worms |
| S1 | N | Mulloon Flat camping area | 35°26.262'S | 149°34.212'E | 4 |
| S2 | N | Lowden Park | 35°30.522'S | 149°36.216' E | 10 |
| S3 | S | Badja Road, Badja A | 35°59.262'S | 149°33.246' E | 5 |
| S4 | N | Kindervale | 35°41.49'S | 149°33.246' E | 13 |
| S5 | MW | Cowangerong Firetrail, Kindervale | 35°42.84S | 149°31.296' E | 2 |
| S6 | MW | Anembo | 35°45.492'S | 149°31.062' E | 13 |
| S7a | MW | Hereford Hall | 35°44.818'S | 149°31.818' E | 14 |
| S8a | MW | Badja Road, Badja B | 35°59.4453'S | 149°32.891' E | 3 |
| **Source:** Velvet worms sampled along the semi-isolated refugia in Tallaganda Region were mostly female and sampled across different seasons. |  |  |  |  |  |

### Dissection, DNA extraction, PCR amplification, and Sequencing

To minimize transient prey DNA, velvet worms (Fig. 2) were unfed for two weeks before dissection. Guts were aseptically removed, flash frozen in liquid nitrogen, and stored at -80°C. DNA extractions were done using the Qiagen DNeasy PowerLyzer PowerSoil kit. Tools were ethanol-sterilized between dissections. These extractions were performed in batches of 20 samples, with the order randomised to avoid consecutive processing of samples from the same site or field collection date, thereby minimising potential batch effects. DNA concentrations were analysed using a Qubit 4 Fluorometer (Life Technologies) and amplicon specificity by gel electrophoresis. The genomic DNA samples from the whole gut of velvet worms were normalized to 1 ng/µl concentration. Together with these samples, the positive (ZymoBIOMICS®) and negative controls were amplified alongside the gut samples to assess contamination (see attached Supplementary qzv files). These controls were included throughout the workflow.

**Fig 1.**
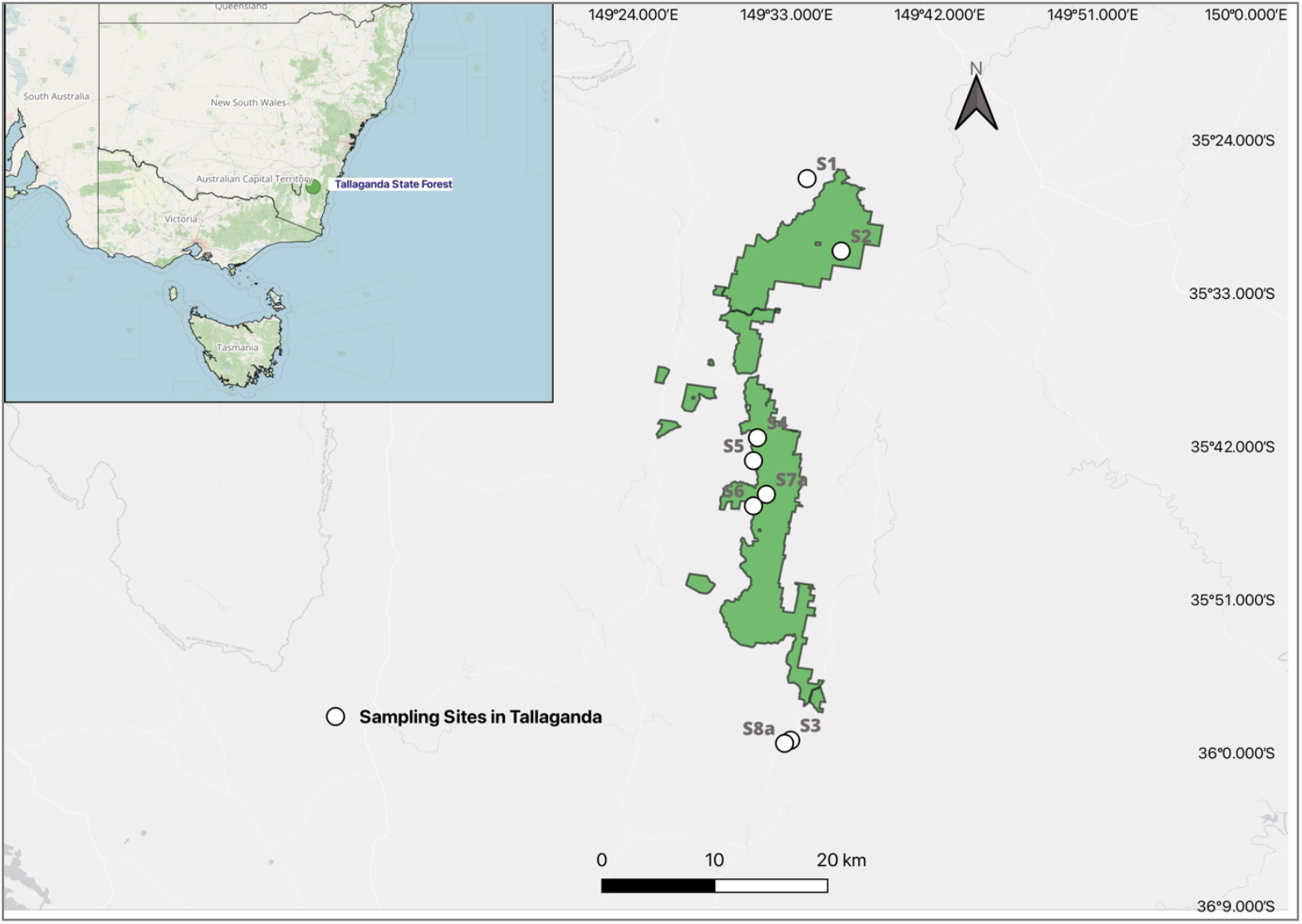
Velvet worm sampling sites. Map of eight velvet worm (*Euperipatoides rowelli)* sampling sites across five semi-isolated refugia in the Tallaganda region, New South Wales, Australia. White circles represent individual locations. Inset: study within Australia (green dot). The map was generated using QGIS (v3.34) using shapefiles from the New South Wales State Forest database and GADM v4.1 databases with base layers from Geoscience Australia.

**Fig 2.**
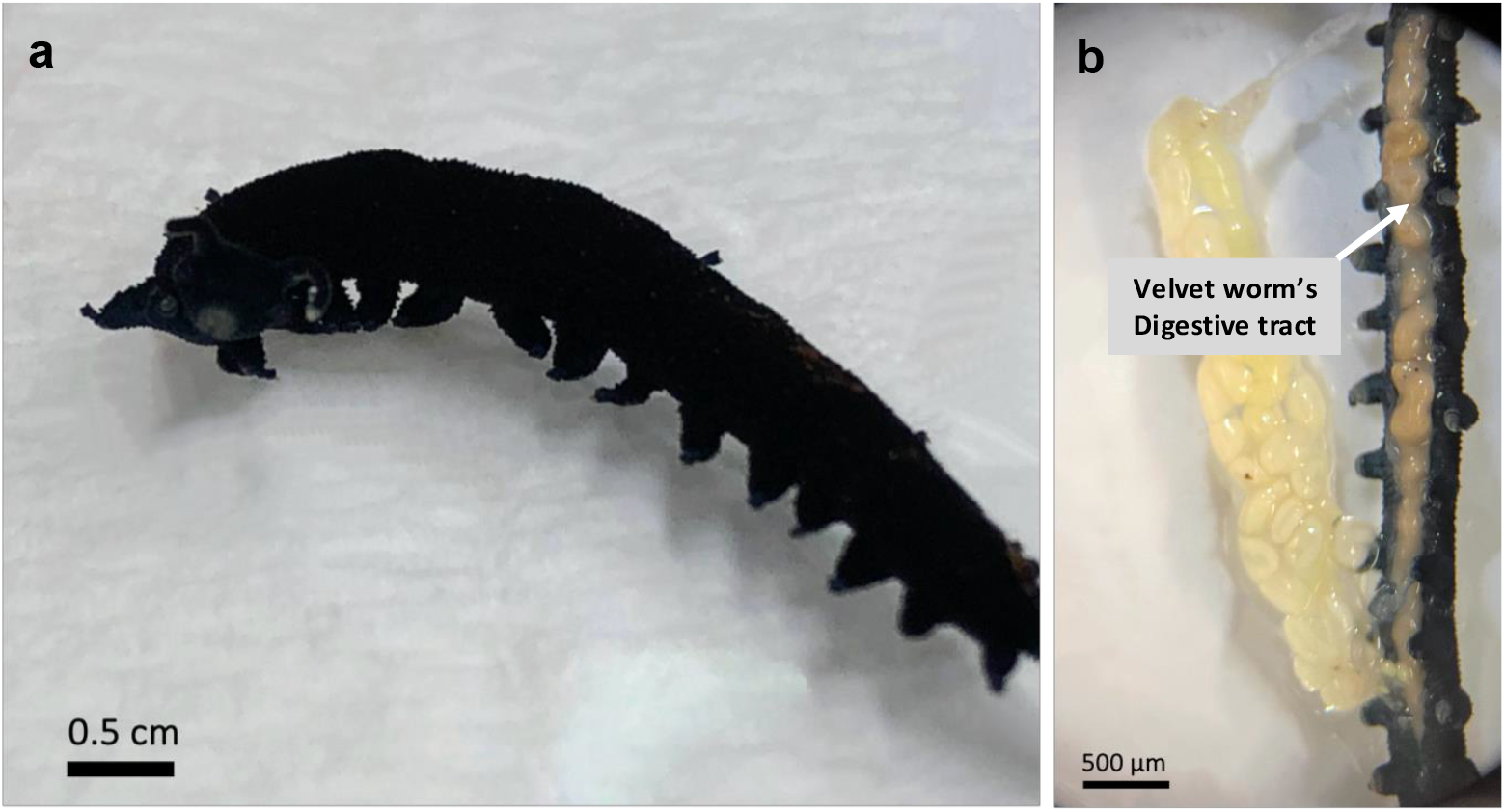
Anatomical context. (a) Adult *E. rowelli*, a saproxylic onychophoran inhabiting decaying logs in temperate forests. (b) Dissected digestive tract showing straight gut morphology. Gut tissues were used for downstream microbial DNA extraction and amplicon sequencing.

The 16S rRNA V4-V5 region was PCR amplified using primer set 515F (5’-GTGYCAGCMGCCGCGGT-3’, Parada et al. 2015) and 926R (5’-CGGYCAATTYMTTTRAGTTT-3’, Quince et al. 2011) according to the protocol described in Needham et al., 2018. Amplicons were pooled and cleaned with AMPure XP beads (Beckman Coulter, CA, USA). The pooled, barcoded amplicon sequencing was carried out at the Australian National University Biomolecular Resource Facility (BRF), ACT, Australia, for sequencing on the Illumina MiSeq platform, producing a 2 x 300 bp mode paired-end read.

### Bioinformatics

Prior to importing the raw reads, metadata were verified using Keemei, an open-source Google Sheets add-on for tabular bioscience file formats [13].

Raw reads were processed using a modified exact amplicon sequence variant pipeline [14] conducted using Quantitative Insights into Microbial Ecology (QIIME2, v2021.11) [15]. Primers from raw reads were removed using *cutadapt* [16]. The raw sequence data were demultiplexed and quality filtered with the q2demux plugin, then quality filtered, denoised, and clustered into ASVs with DADA2 (see Supplementary Table S2 [17]). Taxonomy was assigned to ASVs using the *q2 feature classifier* (Bokulich et al., 2018a) and the SILVA reference database (v132) [18] [19].

### Statistical Analysis

All analyses were conducted in R (v4.1.1, R Core Team 2021) within RStudio (version 2023.09.0). We report nonrarefied data results using relative abundance [20]. Microbial profiles were visualized using stacked bar plots, created using *ggplot2* (version 3.4.3) [21]. Alpha diversity metrics (e.g., Shannon, observed ASVs) were estimated using *phyloseq* (version 1.38.0) [22] and Faith’s phylogenetic diversity (PD) with *picante* (version 1.8.2) [23] packages and tested for group differences using Kruskal-Wallis tests. Beta diversity metrics were estimated using unweighted/weighted UniFrac [24], Bray–Curtis [25], and centered log-ratio (CLR) transformed Euclidean distances. To address the compositional nature of microbiome data, CLR transformation was applied, reducing spurious correlations and ensuring valid multivariate comparisons [12].

To evaluate community differences between sites, permutational analysis of variance (PERMANOVA) was calculated, with analysis of the homogeneity of within-group variation using PERMDISP. Beta diversity was visualized using principal coordinates analysis (PCoA) and nonmetric multidimensional scaling (NMDS) functions on default settings in R *vegan* (version 2.6-4) [26].

The differential abundance analysis using *DESeq2* [27] to evaluate log2-fold changes in CLR-transformed phylum and genus-level abundance across sites. Significantly enriched taxa (adjusted p < 0.05) were visualized in volcano plots.

*UpSetR* (version 1.4.0) was used to generate *UpSet* plots used to explore shared and unique core microbial taxa across sites [28]. These visualisations allow clear representation of taxa consistently present (core microbiota) versus those uniquely associated with specific sites or combinations, offering insight into microbial assembly patterns across fragmented deadwood microhabitats.

Linear mixed-effects models were implemented using *lmerTest* [29] to assess the effects of spatial gradients, with estimated marginal means used for post hoc group comparisons. Full code and reproducible scripts are available in the supplementary material. A compiled version of the R script, generated using *knitr* (v 1.5) [30] is uploaded to a public repository with the link provided in the code availability statement. For detailed methodology, see Supplementary Methods.

## RESULTS

### Sample Collection and Sequencing

Our study characterized the microbial communities inhabiting the onychophoran *E. rowelli* guts across eight sites within the five semi-isolated refugia gradients in the Tallaganda region. Sequencing of V4-V5 16S rRNA gene amplicons obtained from 64 barcoded velvet worm guts yielded a total of 5,903,686 raw reads on the Illumina MiSeq platform. The V4–V5 primer set was selected as it captures both bacterial and archaeal lineages [31][32]. Quality control and filtering steps, including denoising with DADA2 and chimeric sequence removal, produced a final dataset with 624,679 sequence fragments total—an average of 9760 denoised sequences per sample (Supplementary Tables S1-S2 provide full host metadata and sequenced data statistics). Negative controls yielded negligible counts (*Propionibacteria*, ∼10 reads; *Rickettsiella*, ∼2-3 reads). Given their minimal abundance, these taxa were retained in the dataset rather than removed.

Figure 3 shows that *Proteobacteria* (genus: *Ac37b*) and *Firmicutes* (genus: *Spiroplasma*) were the two taxa with high relative abundance found in most velvet worm gut samples in all sites. *Proteobacteria* dominated Site 4 (S4, Kindervale), Site 6 (S6, Anembo), and Site 7a (S7a, Hereford Hall).

**Fig 3.**
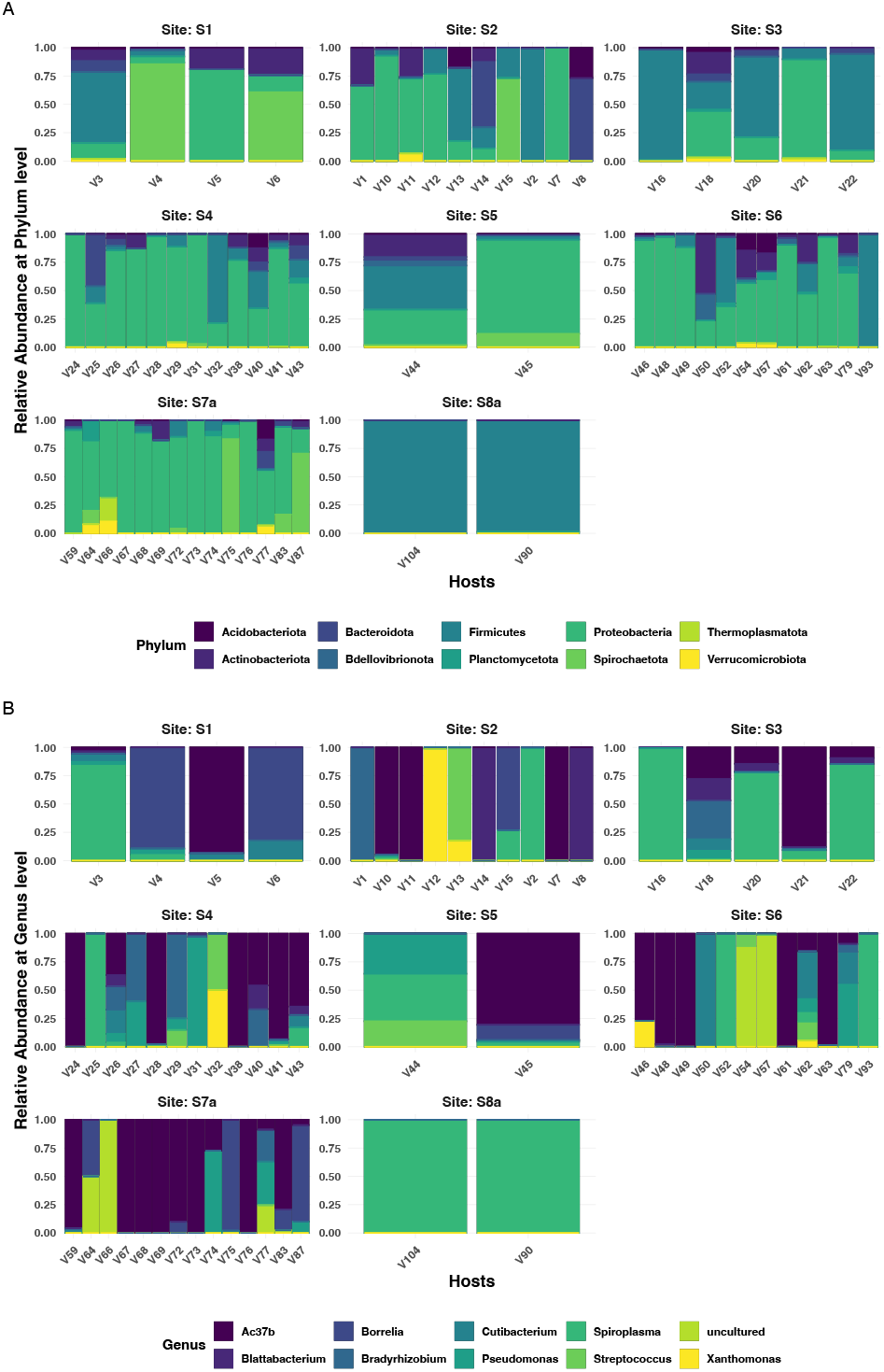
Composition of gut microbial communities in *Euperipatoides rowelli* across eight sites in the Tallaganda region. (a) Stacked bar plots showing the relative abundance of bacterial phyla, highlighting *Proteobacteria* as the prevalent taxa in most samples. Each bar represents an individual velvet worm gut microbiome. **(b)** Stacked bar plots of the same samples at the genus level, with genus *Ac37b* prevalent across most sites except S8a, where its abundance is markedly reduced. Samples V42, V56, and V105 contained ASVs not classified at the genus level and therefore not represented at the genus-level panel.

### Velvet Worm Gut Microbial Diversity

To determine the patterns of gut microbial diversity across the eight sites, the alpha diversity was assessed using Shannon’s index (Shan), Faith’s phylogenetic diversity (PD), and observed species richness (SR) indices for each site (Supplementary Table S3). Among the sites,

Mulloon Flat Camping Area (S1) in the northern region and Badja Road, Badja B (S8a) in the southern region of the Tallaganda Forest had the lowest overall average alpha diversity across all indices. Overall, there were no statistically significant differences in alpha diversity among gut samples taken from eight different sites (Fig. 4; Supplementary Table S3).

**Fig 4.**
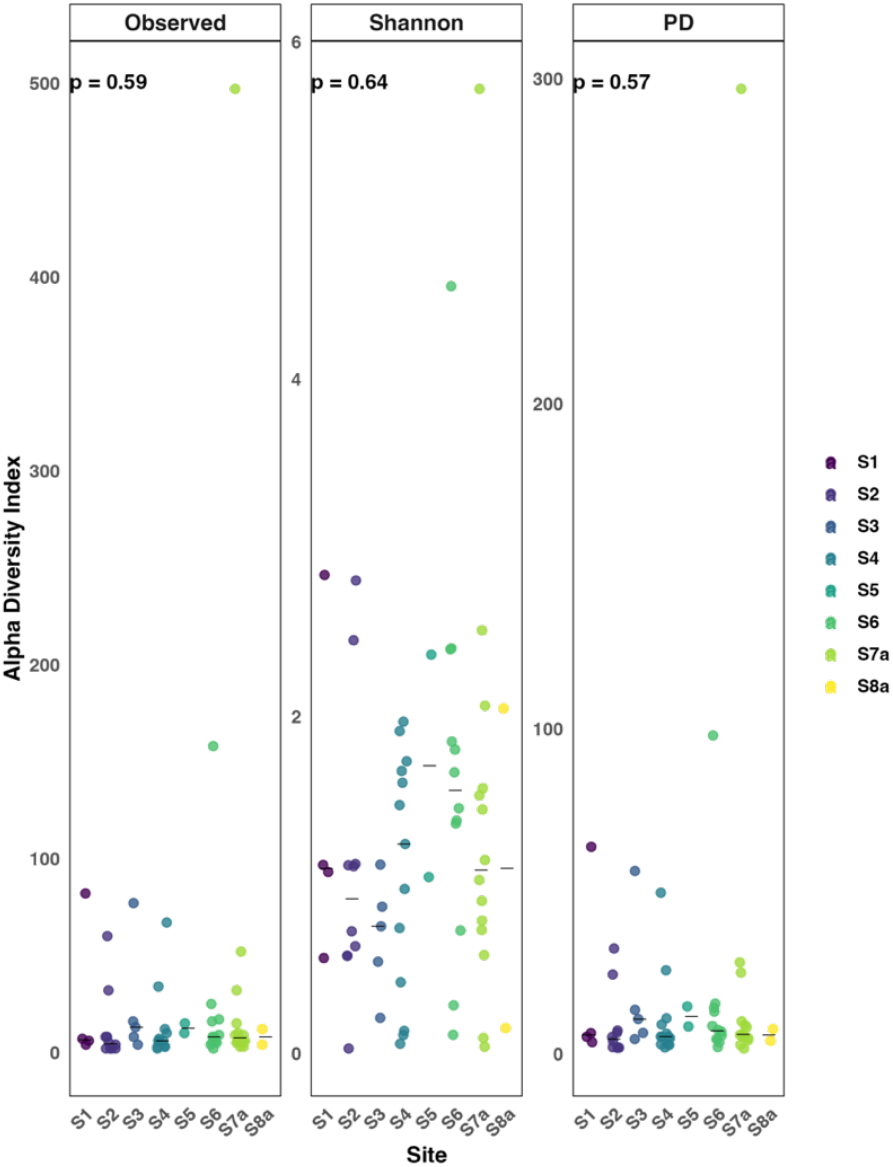
Alpha diversity of gut microbial communities in velvet worms across eight sites in the Tallaganda region. Each dot represents a microbiome from one individual velvet worm. Strip plots showing alpha diversity metrics: Shannon diversity (Shan), Faith’s phylogenetic diversity (PD), and observed species richness (SR) for velvet worm gut microbial communities across eight localities. No significant differences in alpha diversity were observed among sites, indicating broadly similar levels of microbial richness and phylogenetic diversity across the region.

Despite minor differences, statistical testing revealed no significant variation in alpha diversity among sites (Kruskal-Wallis, all p > 0.05; Fig. 4). This finding held when applying the linear mixed-effects model (LMM) [29] analysis to account for unbalanced sample sizes and within-site variability across the sites. Results indicated no significant differences in alpha diversity among sites (all p > 0.05), with and without adjusting for multiple comparisons (see Supplementary Tables S9-S11, Fig. S1). While our analyses primarily emphasize bacterial community composition, the identification of archaeal lineages provides supplementary context for understanding saproxylic gut microbiomes.

### Site Effect on Gut Microbial Beta Diversity

Beta diversity was assessed to determine spatial structuring in gut microbial communities across sampling sites, using four distance metrics: Euclidean (CLR-transformed), Bray-Curtis, unweighted UniFrac (UniFrac), and weighted UniFrac (WUniFrac). Permutational multivariate analysis of variance (PERMANOVA) revealed that variation explained by site was low (R^2^ = 0.07-0.12) across all metrics, with F values ranging from 0.70 to 1.22 (all p > 0.05), indicating no significant differences in community composition among localities (see Supplementary Tables S4-S5).

Ordination analyses (PCoA plots; Fig. 5, see Supplementary Fig. S3A) visualized these relationships, revealing considerable overlap among the sites, suggesting weak spatial structuring. Moreover, the intra-site variability often exceeded inter-site variability, particularly at UniFrac and CLR Euclidean distances, suggesting that velvet worm gut microbiomes were not strongly shaped by geographic differences.

**Fig 5.**
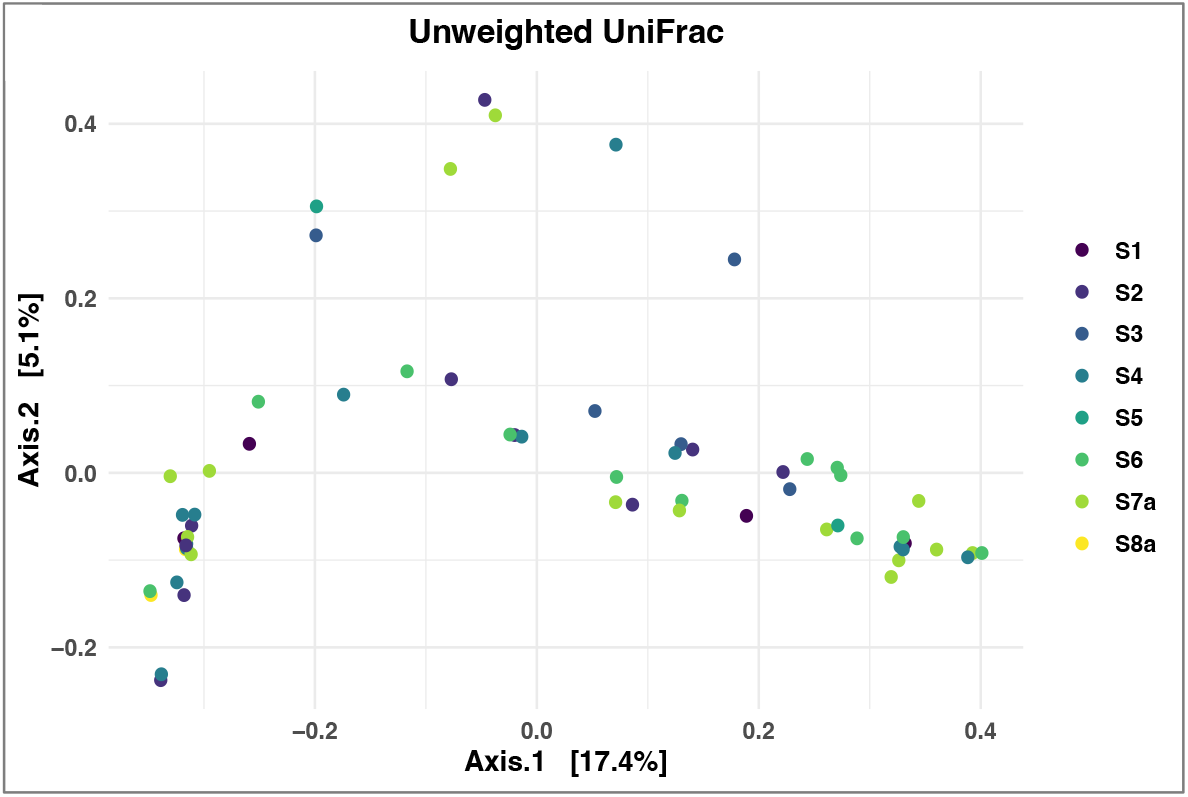
Beta diversity of gut microbial communities in velvet worms across eight sites in the Tallaganda region. Principal Coordinate Analysis (PCoA) plots based on UniFrac distances at the ASV level. The percentage of variance explained by each axis is shown in parentheses. Each point represents an individual velvet worm gut sample, colour-coded by site.

The lack of notable site-level differentiation may indicate several issues. First, velvet worms were held unfed for two weeks prior to dissection to minimize transient prey-associated microorganisms. However, recent studies show that prey-derived bacteria can persist in host guts in dormant states despite hosts being unfed [33] [34]. Second, localized variation in microbial exposure may have been caused by deadwood microhabitats, which differ in decomposition stage and microbial load [37]. Deadwood may function as a buffered microhabitat that shields *E. rowelli* from external environmental fluctuations, particularly moisture and temperature variations, potentially contributing to the observed stability in gut microbial communities across geographically isolated communities. Finally, the residual microbial DNA may indicate the existence of dormant highlighting the challenges in differentiating between transient and resident gut microbial communities [36][37] (see Supplementary Material, Tables S3-S4). Together, these factors may be the primary drivers of the gut microbial community composition in *E. rowelli* rather than geographic distance.

### Shared and unique microbial taxa across sites

Understanding the spatial structure of the microbiome across natural populations remains a central challenge in microbial ecology. To explore gut microbial dynamics in velvet worms (*E. rowelli*), we evaluated shared and unique gut microbial taxa across eight sampling sites within the historical refugia in the Tallaganda region.

A core microbiome threshold was defined as those occurring in ≥ 50% of individual hosts and exhibiting a relative abundance of at least 0.001. These thresholds were selected by comparing core membership across a range of prevalence cut-offs from 30% to 100% [38] [39] [40], ensuring that at least one consistently present taxon was retained in the final core. Taxonomic profiling showed *Proteobacteria* as the predominant phylum among samples. According to the cut-off level used, only one taxon, *Proteobacteria*, mostly genus *Ac37b*, was found. This suggests that velvet worms possess a very limited core microbiome. The consistent dominance at the phylum level contrasts with the absence of *Ac37b* at certain sites, indicating spatial heterogeneity at the genus level, even though it constituted part of the core microbiome in 50% of individuals. This pattern indicates that *Ac37b* may function as a generalist or environmentally acquired symbiont, aligning with the velvet worm’s generalist feeding behaviour and saproxylic environment.

Figure 6 shows a summary of core and unique ASVs using an UpSet plot. The number of taxa shred by each site combination is shown in the top bars, while each column represents a different site combination. The presence of *Proteobacteria* (mostly *Ac37b*) across all sites shows its ubiquity in the velvet worm guts under varying microhabitat conditions. Conversely, the phylum *Bacteroidota* (formerly *Bacteroidetes*) was exclusively identified at site S3 (Badja Road, Badja A), indicating localized microbial signatures. Full core membership matrices are presented in Supplementary Tables S6-S8.

**Fig 6.**
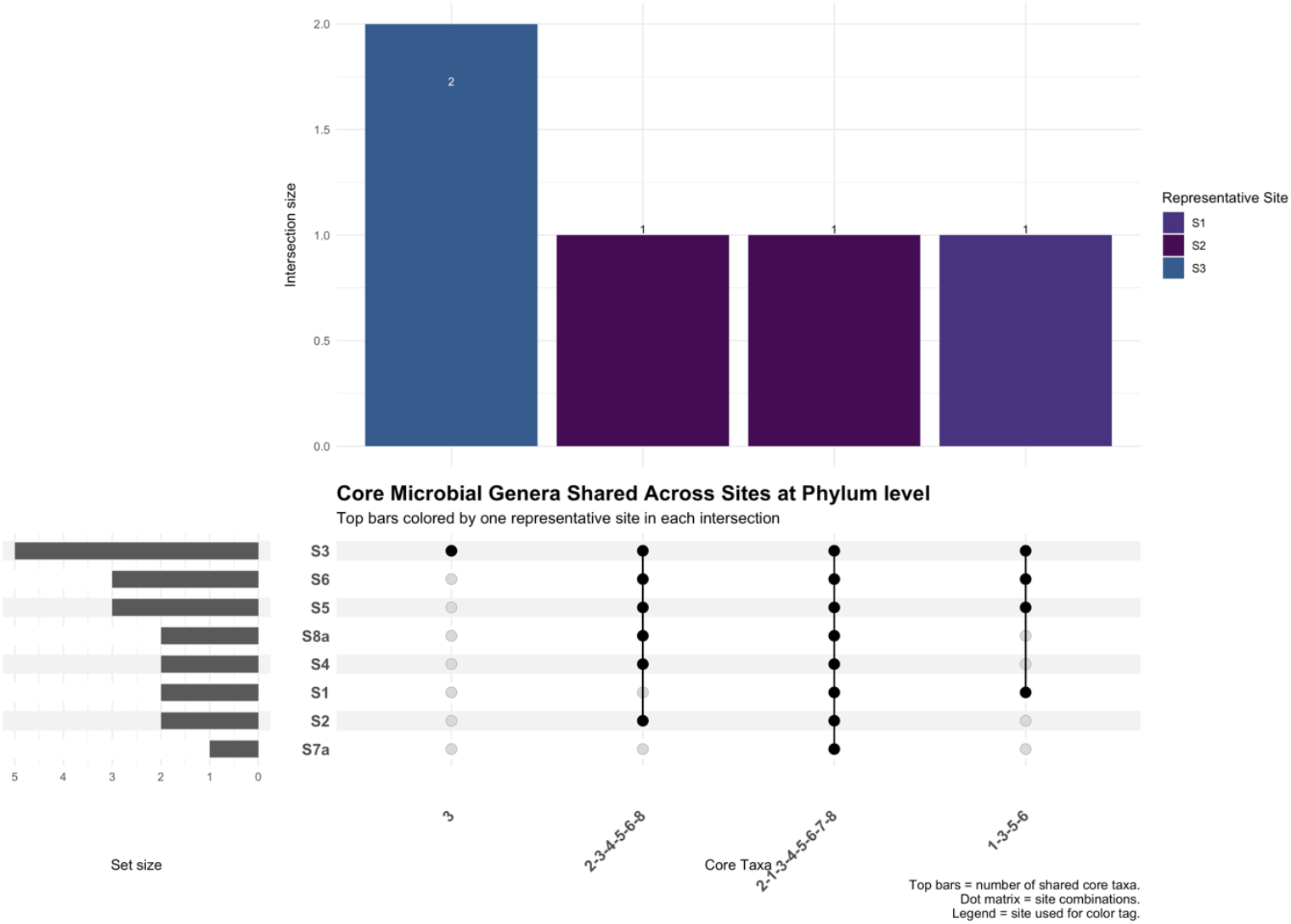
Upset plot of shared and unique ASVs (amplicon sequence variants) in velvet worm gut microbial communities across eight sites in the Tallaganda region. Columns represent site combinations, with colored dots below each column indicating the sites included in each intersection (representative site colors correspond to sampling sites). Top bars indicate the number of taxa shared per intersection. Only one taxon, phylum *Proteobacteria (Ac37b)*, was found in ≥ 50% of individuals across all sites, suggesting a highly limited core microbiome.

The findings suggest that *E. rowelli* possesses a sparse and environmentally influenced gut microbiome. Rather than maintaining a stable gut microbiome, velvet worms may have acquired their microbiome stochastically through local environmental exposure, possibly driven by deadwood decay stages or microhabitat heterogeneity [41].

As observed in this study, a substantial number of velvet worms exhibited a low diversity of gut microbes. This suggests that velvet worms, as a group, do not heavily depend on their gut microbial communities for support. Moreover, this is consistent with the hypothesis that saproxylic velvet worms may acquire much of their microbial community stochastically or environmentally, rather than maintaining a stable, host-specific microbial community.

### Differential Abundance Analysis

Differential abundance analysis using DESeq2 identified microbial genera with significantly different relative abundance across eight velvet worm sampling sites: S1 (Mulloon Flat camping area); S2 (Lowden Park); S3 (Badja Road, Badja A); S4 (Kindervale); S5 (Cowangerong Firetrail, Kindervale); S6 (Anembo); S7a (Hereford Hall); and S8a (Badja Road, Badja B).

Figure 7 summarizes the pairwise comparisons, presenting volcano plots. That illustrates log2 fold changes and statistical significance (-log10 adjusted p-value) for each taxon. The genus *Spiroplasma* (*Firmicutes*, formerly *Tenericutes*) was the most frequently observed differentially abundant taxon, particularly in comparisons involving Sites 5, 6, 7a, and 8a— suggesting its ecological role varies across local habitats. In contrast, an observed differentially reduced relative abundance of *Ac37b* (*Proteobacteria*) in multiple site comparisons. Differences in microbial abundance rather than absolute presence or absence, indicating the influence of microhabitat-level factors. At broader taxonomic levels, *Firmicutes* and *Proteobacteria* were the most differentially abundant phyla across comparisons (see Supplementary Table S12).

**Fig 7.**
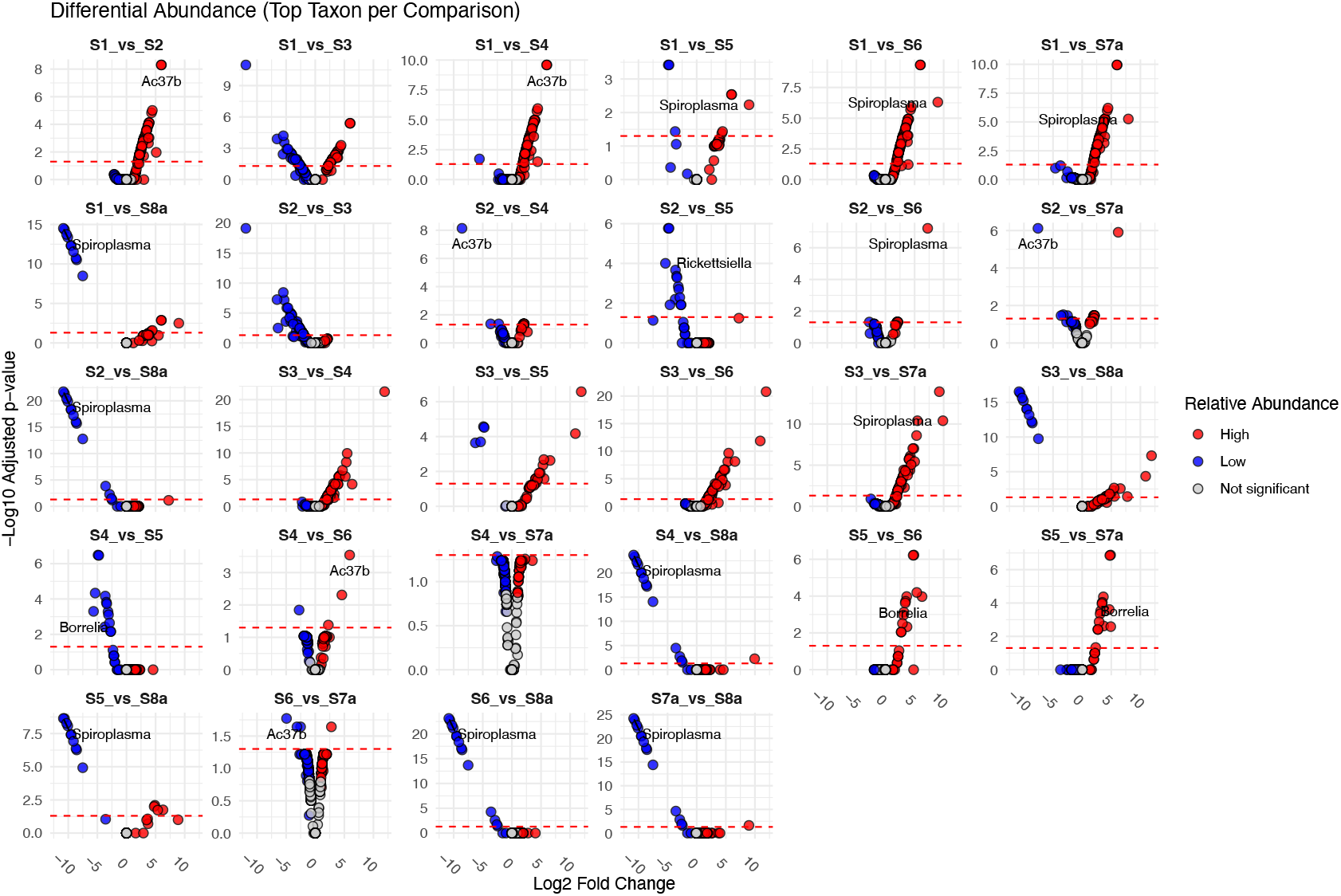
Volcano plots of differentially abundant microbial genera across site pairwise comparisons. Each panel displays log2 fold change (x-axis) vs. -log10 adjusted p-value (y-axis). Taxa significantly enriched in the first site are shown in red (“High”); significantly less abundant taxa are in blue (“Low”); non-significant taxa (adjusted ≥ 0.05) are in gray. A dashed red line marks the significance threshold (adjusted p = 0.05). The top genus per comparison is labelled. *Spiroplasma* frequently observed as the top differentially abundant genus across multiple site comparisons, while *Ac37b* (*Rickettsia)* and, less frequently, *Borrelia* and *Rickettsiella* were also identified as top taxa in select cases.

Collectively, these results imply that while site or microhabitat differences (e.g., deadwood decomposition) cause compositional changes in the velvet worm gut microbiome, these do not result in changes in overall diversity. This is consistent with the idea that the composition of the microbial community in *E. rowelli* is shaped by ambient environmental filtering rather than strong host-specific structuring (see Supplementary Tables S3-S4).

## DISCUSSION

This study is the first to characterize the gut microbiome of *Euperipatoides rowelli*, a saproxylic invertebrate found only in historical refugia in the Tallaganda region (NSW, Australia). The gut microbiome of *E. rowelli* did not exhibit any corresponding site-based spatial differentiation, despite the species’ well-documented genetic divergence throughout the Tallaganda region [42]. Alpha and beta diversity metrics showed little variation across sites, suggesting that host divergence does not significantly influence the composition of microbial communities.

This pattern is comparable with observations in other invertebrates, particularly predatory species, where gut microbial diversity is low and less dependent on host-specific structure [43] [44]. However, factors such as gut morphology [45] [46], diet and feeding mode [10] [47], gut transit time [48], and social behaviour [49] may have more influence than host phylogeny alone.

Moreover, none of the hosts were fed for two weeks prior to dissection in order to reduce the quantity of prey-derived and other transient microbes [50][51]. However, prolonged fasting may have an impact on the gut microbial community. Fasting increases the alpha diversity in certain invertebrates, most likely through commensal organisms breaking down substrates derived from the host.

The relative abundance of *Spiroplasma* (*Firmicutes*, formerly *Tenericutes*) and *Ac37b (Proteobacteria)* was observed to be generally low across sites. This suggests the presence of underlying ecological processes such as competitive exclusion, niche partitioning, or differential environmental filtering [52].

Some species of *Spiroplasma* engage in symbioses, which are suggested as a form of defense against parasitoids and pathogens and may also act as a form of modulation of arthropod reproduction [53][54]. In contrast, *Ac37b* is an obligate intracellular bacterium that may interact with the host via a fundamentally different mechanism [55][56]. Some extracellular taxa which inhabit the gut lumen and interact with host cells indirectly through metabolites or competitive exclusion, intracellular symbionts like *Ac37b* reside within host cells and may directly modulate host function. Distinct life histories may result in antagonistic interactions or priority effects, wherein early colonists inhibit the establishment of subsequent taxa [57][55]. Future research (e.g., meta-transcriptomic or metagenomic profiling) is needed to disentangle patterns of microbial succession and potential host interactions.

While taxonomic shifts are evident, the functional roles of *Spiroplasma* and *Ac37b* in *E. rowelli* remain unknown. The feeding habits and gut architecture of velvet worms make it uncertain to determine whether these microorganisms are transient, environmentally acquired, or contribute to digestion, immunity, or host metabolism.

Functional inference pipelines, or shotgun metagenomics, could assist in identifying whether these taxa encode metabolic traits relevant to nutrient acquisition or other ecological functions [58][59]. These observed compositional patterns emphasize the need for future research linking taxonomic presence with functional potential, particularly in microbiomes in invertebrate lineages.

Interpreting such shifts is complicated by the compositional nature of microbiome data. While centered log-ratio transformation helps mitigate these distortions [12], relative increases may result from a given taxon, reductions in others, or both [60][61]. Nonetheless, compositional shifts, not changes in overall diversity, emerged as the primary feature among sites.

Core microbiome analysis revealed a single genus, Ac37b, present in ≥ 50% of individuals across all sites [62][38][63][64]. This supports the view that *E. rowelli* hosts an environmentally structured microbiome rather than a stable, host-specific microbiome [65][40]. The exclusive occurrence of *Spiroplasma* in Site 3 further highlights the influence of local conditions. Although the concept of a core microbiome has been widely used, [66] recent studies emphasize the functional role of rare taxa and challenge the rigid definitions of “core” [67][68].

Low microbial diversity or the absence of stable gut communities has also been observed in several invertebrates [69][43][70]. Their biology likely contributes: a simple gut [8], short retention times, and extra-oral digestion likely impede stable gut microbial colonization [71][72]. Furthermore, *E. rowelli* might be more dependent on prey-derived taxa or transient ambient microorganisms [73] [74]. Velvet worms were unfed for two weeks prior to dissection to reduce the influence of transient, prey-associated bacteria. However, it remains unclear whether this unfed period also affected resident gut microbial communities. In invertebrates, distinctions between resident and transient microorganisms can be fluid, as some taxa may persist from prior diets or enter dormant states during unfed periods. This corresponds with broader patterns reported in carnivorous invertebrates [43][75]. Understanding the reasons some animals fall at the lower end of the microbial dependence spectrum, characteristics linked to this pattern include carnivory, extra-oral digestion, and rudimentary gut anatomy, continues to pose a challenge [76][77][78][79][43].

Temporal and environmental factors may influence microbiome composition. The velvet worm samples were collected from the period prior to and following the 2019 Australian bushfires (2018-2021). These disturbances may have impacted prey availability or host physiology; but such variation may not be fully captured here. Additionally, reliance on microhabitats such as the dead wood may affect microbial diversity.

## CONCLUSION

In summary, *E. rowelli* hosts a gut microbiome that is sparse and shows limited geographic structuring across historical refugia. Instead, patterns point to local ecological filtering within deadwood microhabitats and to host biology (e.g., predatory lifestyle, extra-oral digestion) that reduces the need for stable microbial associations. These findings suggest that velvet worms harbour transient, environmentally structured gut microbial communities, underscoring the value of studying underrepresented taxa to broaden our understanding of animal-microbe associations.

## Abbreviations

DESeq2: Differential Expression Sequencing 2;
EOD: Extra-Oral Digestion;
LMM: Linear Mixed-Effects Model;
rRNA: Ribosomal Ribonucleic Acid;
VW: Velvet Worm;
CLR: Center-Log Ratio;
PERMANOVA: Permutational Multivariate Analysis;
PERMADISP: Permutational Multivariate Analysis of Dispersion;
NMDS: Non-Metric Multidimensional Scaling;
PCoA: Principal Coordinate Analysis

## Acknowledgements

The authors gratefully acknowledge Dr Oliver Stuart and Dr Rosie Harris for their assistance with fieldwork. Sincere appreciation is extended to Mr Wesley Keys for his continued dedication and field support following his retirement. The authors also thank Associate Professor Paul Cooper for invaluable assistance with laboratory reagents and Dr Jack Eggerton and Dr Monica Ruibal for their support with logistics and equipment.

## Author Contributions

Study conception and design: IF, AR, DR; Sample collection and laboratory work: IF, DR, TW; Bioinformatics and statistical analysis: IF, TW, TL, FT; Data interpretation: IF, UM, TW, TL, FT; Manuscript and editing: ILF, AR, UM, DR, TW, TL, FT; Supervision/Advisory Contributions: AR, UM, DR

## Ethical Statement

Field collections and experimental procedures involving *Euperipatoides rowelli* were conducted in accordance with Australian National University (ANU) guidelines for the ethical treatment of invertebrates in research and under Scientific Licence SL102322 issued to Dr David Rowell by the NSW National Parks and Wildlife Service under the Biodiversity Conservation Act 2016, authorizing invertebrate collection and research within Tallaganda National Park, New South Wales.

## References

1. Shade, A., & Handelsman, J. Beyond the Venn diagram: the hunt for a core microbiome. Environ. Microbiol. 2012; 141 4–12.

2. Philippot, L., Griffiths, B. S., & Langenheder, S. Microbial community resilience across ecosystems and multiple disturbances. Microbiol. Mol. Biol. Rev. 2021; 852 10–1128.

3. Cappellato, M., Baruzzo, G., Patuzzi, I., et al. Modeling microbial community networks: methods and tools. Curr. Genomics 2021; 224 267.

4. Berry, D., & Widder, S. Deciphering microbial interactions and detecting keystone species with co-occurrence networks. Front. Microbiol. 2014; 5 219

5. Barclay, S. D., Rowell, D. M., & Ash, J. E. Pheromonally mediated colonization patterns in the velvet worm Euperipatoides rowelli (Onychophora). J. Zool. 2000; 2504 437–446.

6. Garrick, R. C., Sands, C. J., Rowell, D. M., et al. Phylogeography recapitulates topography: very fine-scale local endemism of a saproxylic ‘giant’springtail at Tallaganda in the Great Dividing Range of southeast Australia. Mol. Ecol. 2004; 1311 3329–3344.

7. Hope, G. Hope, G.S. (1994). 15 Quaternary vegetation. History of the Australian vegetation: Cretaceous to recent, 368. 15 Quat. Veg. Hist. Aust. Veg. Cretac. Recent 1994 1994; Jul 28:368:368.

8. Vannier, J., Liu, J., Lerosey-Aubril, R., et al. Sophisticated digestive systems in early arthropods. Nat. Commun. 2014; 51 1–9.

9. Pastorelli, R., Paletto, A., Agnelli, A.E., Lagomarsino, A., De Meo, I. Microbial Diversity and Ecosystem Functioning in Deadwood of Black Pine of a Temperate Forest. Forests 2021; 10 18;12;(10):1418.

10. Ley, R. E., Peterson, D. A., & Gordon, J. I. Ecological and evolutionary forces shaping microbial diversity in the human intestine. Cell 2006; 1244 837–848

11. Lange, C., Boyer, S., Bezemer, T.M., Lefort, M.-C., Dhami, M.K., Biggs, E., et al. Impact of intraspecific variation in insect microbiomes on host phenotype and evolution. ISME J. 2023; 11 1;17;(11):1798–1807.

12. Gloor, G. B., & Reid, G. Compositional analysis: a valid approach to analyze microbiome high-throughput sequencing data. Can. J. Microbiol. 2016; 628 692–703.

13. Rideout, J. R., Chase, J. H., Bolyen, E., et al. Keemei: cloud-based validation of tabular bioinformatics file formats in Google Sheets. Gigascience 2016; 51 S13742–016.

14. Needham, D. M., Fichot, E. B., Wang, E., et al. Dynamics and interactions of highly resolved marine plankton via automated high-frequency sampling. ISME J. 2018; 1210 2417–2432.

15. Bolyen, E., Rideout, J. R., Dillon, M. R., et al. Reproducible, interactive, scalable and extensible microbiome data science using QIIME 2. Nat. Biotechnol. 2019; 378 852–857.

16. Martin, M. Cutadapt removes adapter sequences from high-throughput sequencing reads. EMBnet J. 2011; 171 10–12.

17. Callahan, B. J., McMurdie, P. J., Rosen, M. J., et al. DADA2: High-resolution sample inference from Illumina amplicon data. Nat. Methods 2016; 137 581–583.

18. Quast, C., Pruesse, E., Yilmaz, P., Gerken, J., Schweer, T., Yarza, P., et al. The SILVA ribosomal RNA gene database project: improved data processing and web-based tools. Nucleic Acids Res. 2012 11 27;41;(D1): D590–D596.

19. Yilmaz, P., Parfrey, L. W., Yarza, P., et al. The SILVA and “all-species living tree project (LTP)” taxonomic frameworks. Nucleic Acids Res. 2014; 42D1 D643–D648.

20. McMurdie, P. J., & Holmes, S. Waste not, want not: why rarefying microbiome data is inadmissible. PLoS Comput. Biol. 2014; 104 E1003531.

21. Wickham, H. ggplot2. Wiley Interdiscip. Rev. Comput. Stat. 2011; 32 180–185.

22. McMurdie, P. J., & Holmes, S. phyloseq: An R Package for Reproducible Interactive Analysis and Graphics of Microbiome Census Data. Tech Rep Stanf. Univ Stanf. CA USA 2019.

23. Kembel, S. W., Cowan, P. D., Helmus, M. R., et al. Picante: R tools for integrating phylogenies and ecology. Bioinforma. 2010; 2611 1463–1464.

24. Lozupone, C., Hamady, M., & Knight, R. UniFrac–an online tool for comparing microbial community diversity in a phylogenetic context. BMC Bioinforma. 2006; 71 1–14.

25. Bray, J. R., & Curtis, J. T. An ordination of the upland forest communities of southern Wisconsin. Ecol. Monogr. 1957; 274 326–349.

26. Oksanen, J., Blanchet, F. G., Kindt, R., et al. Community ecology package. R Package Version 2018; 2 5–2.

27. Love, M. I., Huber, W., & Anders, S. Moderated estimation of fold change and dispersion for RNA-seq data with DESeq2. Genome Biol. 2014; 1512 1–21.

28. Conway, J.R., Lex, A., Gehlenborg, N. UpSetR: an R package for the visualization of intersecting sets and their properties. Bioinformatics 2017; 09 15;33;(18):2938–2940.

29. Kuznetsova, A., Brockhoff, P. B., & Christensen, R. H. B. lmerTest package: tests in linear mixed effects models. J. Stat. Softw. 2017; 8213.

30. Xie, Y. Xie Y (2025). knitr: A General-Purpose Package for Dynamic Report Generation in R. R package version 1.50, https://yihui.org/knitr/. Knitr Gen.-Purp. Package Dyn. Rep. Gener. R R Package Version x2025; 150.

31. Abellan-Schneyder, I., Matchado, M.S., Reitmeier, S., Sommer, A., Sewald, Z., Baumbach, J., et al. Primer, Pipelines, Parameters: Issues in 16S rRNA Gene Sequencing. mSphere 2021; 02 24;6;(1): e01202–20.

32. Varliero, G., Lebre, P.H., Adams, B., Chown, S.L., Convey, P., Dennis, P.G., et al. Biogeographic survey of soil bacterial communities across Antarctica. Microbiome 2024; 01 12;12;(1):9.

33. Kennedy, S. R., Tsau, S., Gillespie, R., et al. Are you what you eat? A highly transient and prey-influenced gut microbiome in the grey house spider Badumna longinqua. Mol. Ecol. 2020; 1001–1015 295.

34. Chalifour, B., & Li, J. Characterization of the gut microbiome in wild rocky mountainsnails (Oreohelix strigosa). Anim. Microbiome 2021; 3 1–13.

35. Baniel, A., Amato, K.R., Beehner, J.C., Bergman, T.J., Mercer, A., Perlman, R.F., et al. Seasonal shifts in the gut microbiome indicate plastic responses to diet in wild geladas. Microbiome 2021; 01 23;9;(1):26.

36. Zhang, C., Derrien, M., Levenez, F., et al. Ecological robustness of the gut microbiota in response to ingestion of transient food-borne microbes. ISME J. 2016; 109 2235–2245.

37. McDonald, M. D., Owusu-Ansah, C., Ellenbogen, J. B., et al. What is microbial dormancy? Trends Microbiol. 2024; 322 142–150.

38. Ainsworth, T. D., Fordyce, A. J., & Camp, E. F. The other microeukaryotes of the coral reef microbiome. Trends Microbiol. 2017; 2512 980–991.

39. Aira, M., Pérez-Losada, M., Crandall, K.A., Domínguez, J. Host taxonomy determines the composition, structure, and diversity of the earthworm cast microbiome under homogenous feeding conditions. FEMS Microbiol. Ecol. 2022; 08 23;98;(9): fiac093.

40. Neu, A. T., Allen, E. E., & Roy, K. Defining and quantifying the core microbiome: Challenges and prospects. Proc. Natl. Acad. Sci. 2021; 11851 E2104429118.

41. Shade, A., & Stopnisek, N. Abundance-occupancy distributions to prioritize plant core microbiome membership. Curr. Opin. Microbiol. 2019; 49 50–58.

42. Bull, J.K., Sunnucks, P. Strong genetic structuring without assortative mating or reduced hybrid survival in an onychophoran in the Tallaganda State Forest region, Australia: Properties of an Onychophoran Hybrid Zone. Biol. J. Linn. Soc. 2014 03;111;(3):589–602.

43. Hammer, T. J., Janzen, D. H., Hallwachs, W., et al. Caterpillars lack a resident gut microbiome. Proc. Natl. Acad. Sci. 2017; 11436 9641–9646.

44. Hammer, T. J., Sanders, J. G., & Fierer, N. Not all animals need a microbiome. FEMS Microbiol. Lett. 2019; 36610 Fnz117.

45. Engel, P., & Moran, N. A. The gut microbiota of insects–diversity in structure and function. FEMS Microbiol. Rev. 2013; 375 699–735.

46. Rodríguez, J. M., Murphy, K., Stanton, C., et al. The composition of the gut microbiota throughout life, with an emphasis on early life. Microb. Ecol. Health Dis. 2015; 261 26050.

47. Spor, A., Koren, O., & Ley, R. Unravelling the effects of the environment and host genotype on the gut microbiome. Nat. Rev. Microbiol. 2011; 94 279–290.

48. McConnell, E. L., Fadda, H. M., & Basit, A. W. Gut instincts: explorations in intestinal physiology and drug delivery. Int. J. Pharm. 2008; 3642 213–226.

49. Reinhard, J., & Rowell, D. M. Social behaviour in an Australian velvet worm, Euperipatoides rowelli (Onychophora: Peripatopsidae). 2005; J. Zool. 2671 p1-7.

50. Hu, G., Zhang, L., Yun, Y., Peng, Y. Taking insight into the gut microbiota of three spider species: No characteristic symbiont was found corresponding to the special feeding style of spiders. Ecol. Evol. 2019; 07;9;(14):8146–8156.

51. Liu, Y., Liu, J., Zhang, X., Yun, Y. Diversity of Bacteria Associated with Guts and Gonads in Three Spider Species and Potential Transmission Pathways of Microbes within the Same Spider Host. Insects 2023; 09 29;14;(10):792.

52. Debray, R., Herbert, R.A., Jaffe, A.L., Crits-Christoph, A., Power, M.E., Koskella, B. Priority effects in microbiome assembly. Nat. Rev. Microbiol. 2022; 02;20;(2):109–121.

53. Duron, O., Bouchon, D., Boutin, S., Bellamy, L., Zhou, L., Engelstädter, J., et al. The diversity of reproductive parasites among arthropods: Wolbachiado not walk alone. BMC Biol. 2008; 12;6;(1):27.

54. Ballinger, M.J., Perlman, S.J. The defensive Spiroplasma. Curr. Opin. Insect Sci. 2019; 04; 32:36–41.

55. Bastounis, E.E., Radhakrishnan, P., Prinz, C.K., Theriot, J.A. Mechanical Forces Govern Interactions of Host Cells with Intracellular Bacterial Pathogens. Microbiol. Mol. Biol. Rev. 2022; 06 15;86;(2): e00094-20.

56. Su, S., Hong, M., Cui, M. Y., et al. Microbial diversity of ticks and a novel typhus group Rickettsia species (Rickettsiales bacterium Ac37b) in Inner Mongolia, China. Parasite 2023; 3058.

57. García-Bayona, L., Comstock, L.E. Bacterial antagonism in host-associated microbial communities. Science 2018; 09 21;361;(6408): eaat2456.

58. Ellegaard, K.M., Engel, P. Beyond 16S rRNA Community Profiling: Intra-Species Diversity in the Gut Microbiota. Front. Microbiol. 2016; 09 21 7.

59. Yang, S.-Y., Han, S.M., Lee, J.-Y., Kim, K.S., Lee, J.-E., Lee, D.-W. Advancing Gut Microbiome Research: The Shift from Metagenomics to Multi-Omics and Future Perspectives. J. Microbiol. Biotechnol. 2025; 03 26;35: e2412001.

60. Tsilimigras, M.C.B., Fodor, A.A. Compositional data analysis of the microbiome: fundamentals, tools, and challenges. Ann. Epidemiol. 2016; 05;26;(5):330–335.

61. Lloréns-Rico, V., Vieira-Silva, S., Gonçalves, P.J., Falony, G., Raes, J. Benchmarking microbiome transformations favors experimental quantitative approaches to address compositionality and sampling depth biases. Nat. Commun. 2021; 06 11;12;(1):3562.

62. Sabree, Z. L., Hansen, A. K., & Moran, N. A. Independent studies using deep sequencing resolve the same set of core bacterial species dominating gut communities of honey bees. PloS One 2012; 77 E41250.

63. Sweet, M. J., & Bulling, M. T. On the importance of the microbiome and pathobiome in coral health and disease. Front. Mar. Sci. 2017; 4 9.

64. Risely, A. Applying the core microbiome to understand host–microbe systems. J. Anim. Ecol. 2020; 897 1549–1558.

65. Louca, S., Polz, M. F., Mazel, F., et al. Function and functional redundancy in microbial systems. Nat. Ecol. Evol. 2018; 26 936–943.

66. Turnbaugh, P. J., Ley, R. E., Hamady, M., et al. The human microbiome project. Nat. 2007; 4497164 804–810.

67. Jousset, A., Bienhold, C., Chatzinotas, A., Gallien, L., Gobet, A., Kurm, V., et al. Where less may be more: how the rare biosphere pulls ecosystems strings. ISME J. 2017; 04 1;11;(4):853–862.

68. Antwis, R. E., Edwards, K. L., Unwin, B., et al. Rare gut microbiota associated with breeding success, hormone metabolites and ovarian cycle phase in the critically endangered eastern black rhino. Microbiome 2019; 71 1–12.

69. Betcher, M.A., Fung, J.M., Han, A.W., O’Connor, R., Seronay, R., Concepcion, G.P., et al. Microbial Distribution and Abundance in the Digestive System of Five Shipworm Species (Bivalvia: Teredinidae). PLoS ONE 2012; 09 20;7;(9): e45309.

70. Kwong, W. K., Medina, L. A., Koch, H., et al. Dynamic microbiome evolution in social bees. Sci. Adv. 2017; 33 E1600513.

71. Appel, H. M., & Schultz, J. C. Oak tannins reduce effectiveness of Thuricide (Bacillus thuringiensis) in the gypsy moth (Lepidoptera: Lymantriidae). J. Econ. Entomol. 1994; 876 1736–1742.

72. Dillon, R. J., & Dillon, V. M. The gut bacteria of insects: nonpathogenic interactions. Annu. Rev. Entomol. 2004; 491 71–92.

73. Moran, N. A., Ochman, H., & Hammer, T. J. Evolutionary and ecological consequences of gut microbial communities. Annu. Rev. Ecol. Evol. Syst. 2019; 50 451–475.

74. Weingarten, E. A., Atkinson, C. L., & Jackson, C. R. The gut microbiome of freshwater Unionidae mussels is determined by host species and is selectively retained from filtered seston. PLoS One 2019; 1411 E0224796.

75. Kroetsch, S.A., Kidd, K.A., Monk, W.A., Culp, J.M., Compson, Z.G., Pavey, S.A. The effects of taxonomy, diet, and ecology on the microbiota of riverine macroinvertebrates. Ecol. Evol. 2020;12;10;(24):14000–14019.

76. Shelomi, M., Lo, W. S., Kimsey, L. S., et al. Analysis of the gut microbiota of walking sticks (Phasmatodea). BMC Res. Notes 2013; 61 1–10.

77. Coon, K. L., Vogel, K. J., Brown, M. R., et al. Mosquitoes rely on their gut microbiota for development. Mol. Ecol. 2014; 2311 2727–2739.

78. Yun, J.-H., Roh, S.W., Whon, T.W., Jung, M.-J., Kim, M.-S., Park, D.-S., et al. Insect Gut Bacterial Diversity Determined by Environmental Habitat, Diet, Developmental Stage, and Phylogeny of Host. Appl. Environ. Microbiol. 2014; 09;80;(17):5254–5264.

79. Sanders, J. G., Beichman, A. C., Roman, J., et al. Baleen whales host a unique gut microbiome with similarities to both carnivores and herbivores. Nat. Commun. 2015; 61 1–8.

