## Supplementary_Figures for "“Gut microbial communities of velvet worm *Euperipatoides rowelli* (Onychophora) across deadwood microhabitats in southeastern Australia”"

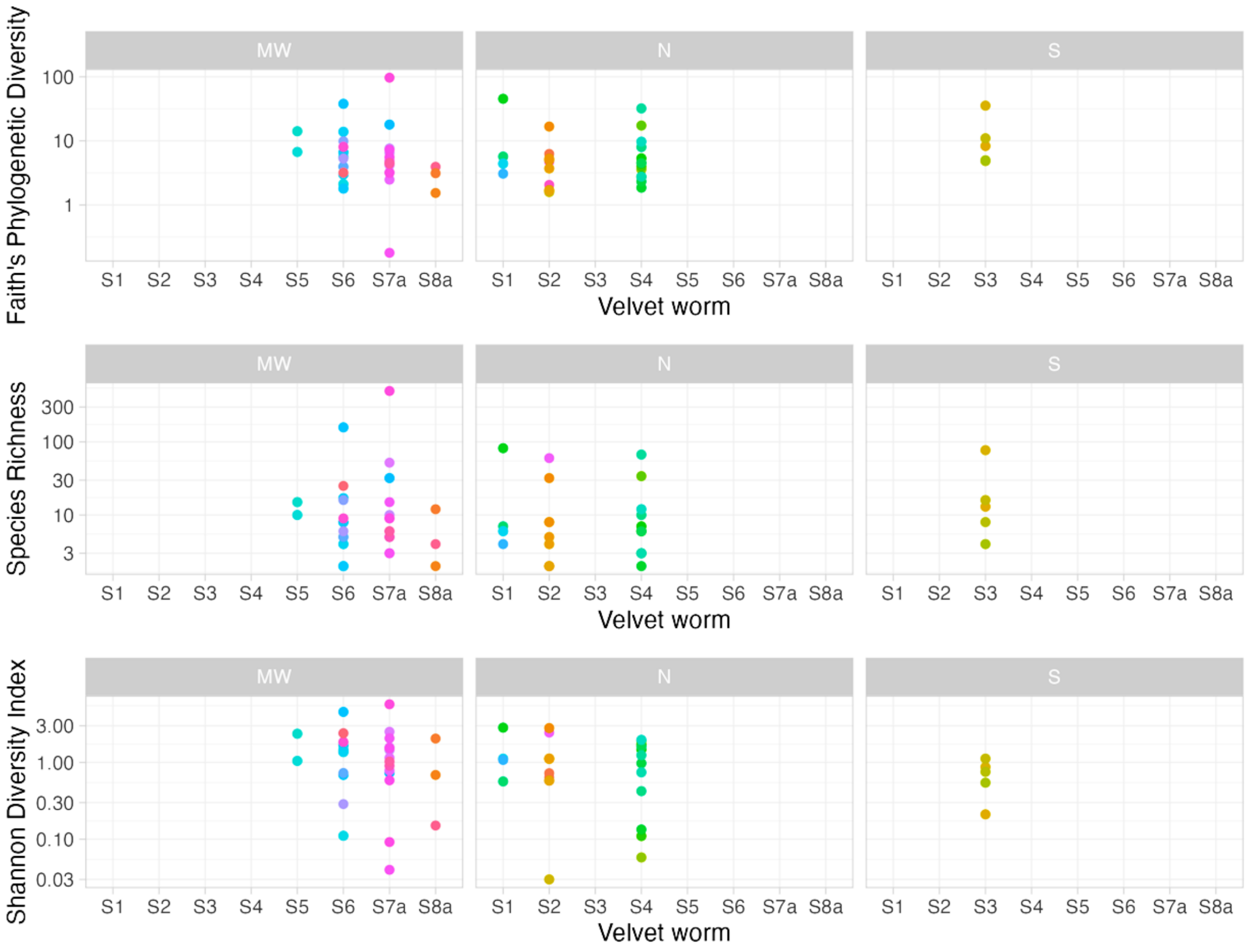


**Fig S1 | Log-Transformed Alpha Diversity Indices of *Euperipatoides rowelli* Gut Microbiota Across Sites, Grouped by Gradient (N, MW, S)**

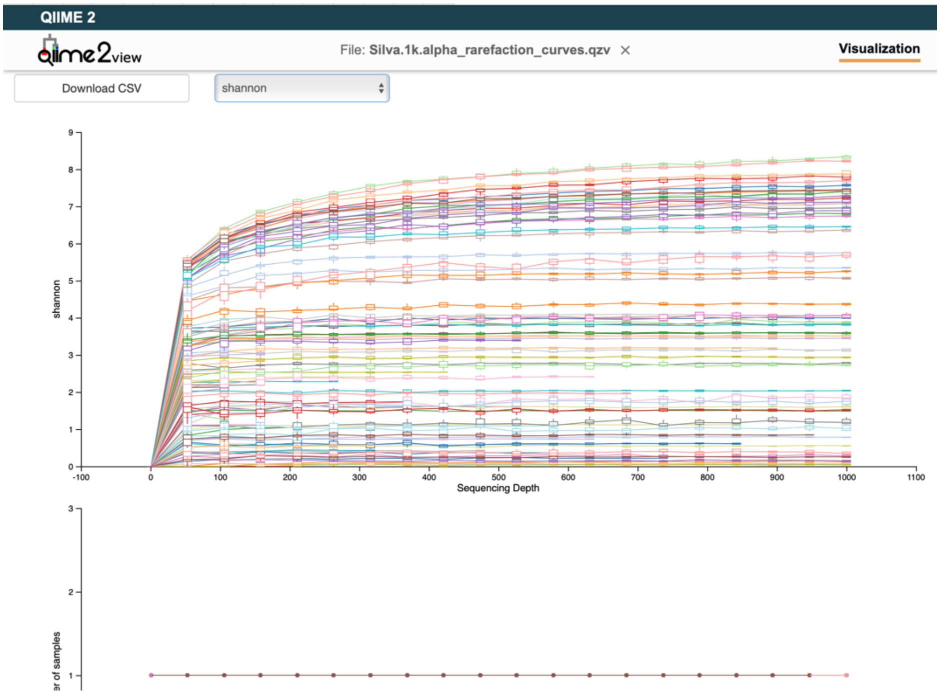


Fig. S2A | **Alpha-rarefaction** of velvet worm at 1000 reads, Shannon

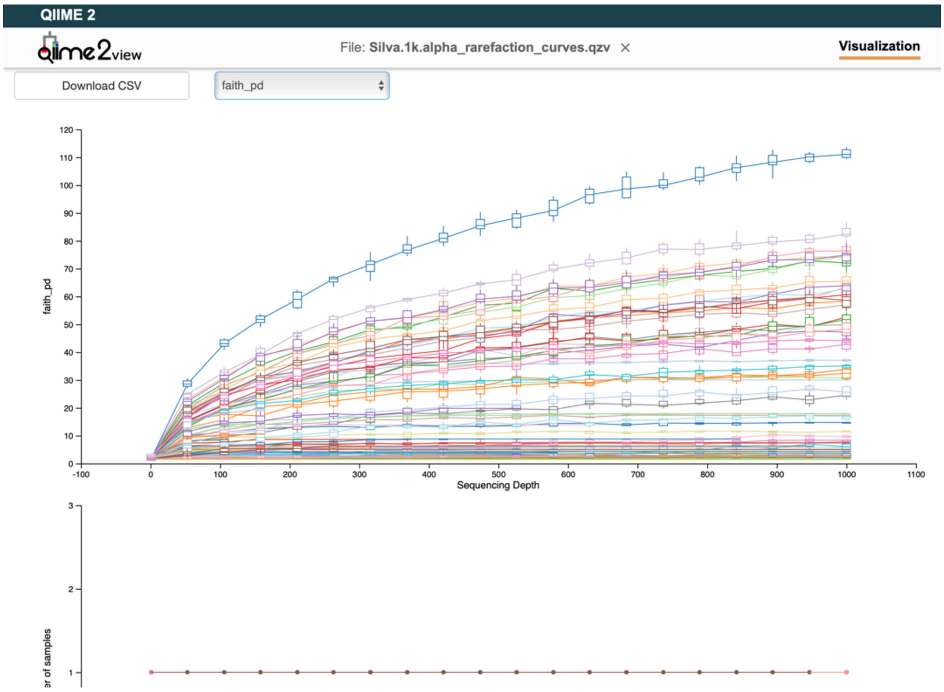


Fig. S2B | **Alpha-rarefaction** of velvet worm at 1000 reads, Faith's PD

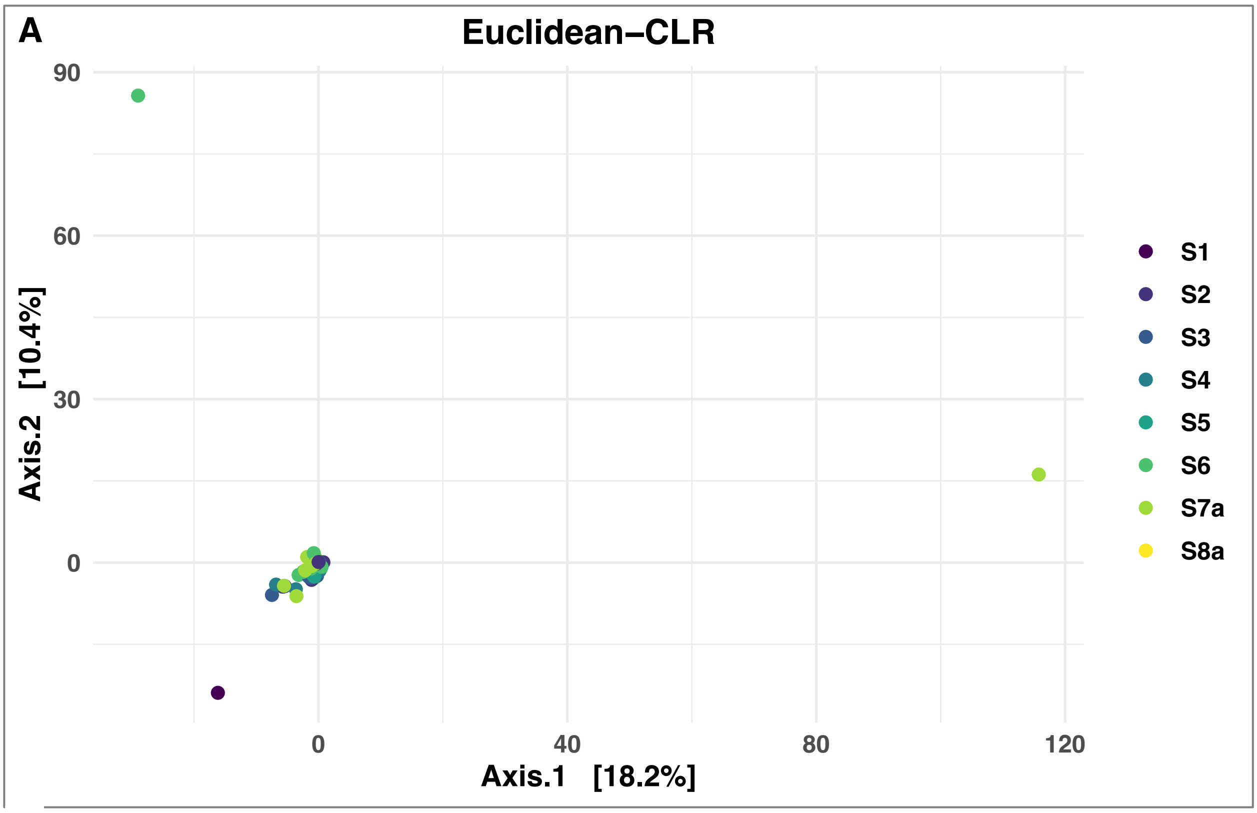


**Fig S3A| Principal Coordinate Analysis** (PCoA) plots based on centered log-ratio (CLR)-transformed Euclidean distances at the ASV level. The percentage of variance explained by each axis is shown in parentheses. Each point represents an individual velvet worm gut sample, colour-coded by site
