## Supplementary_Tables for "“Gut microbial communities of velvet worm *Euperipatoides rowelli* (Onychophora) across deadwood microhabitats in southeastern Australia”"

**Table S1. Velvet Worms MIxS Metadata**

| | | | | Environment | | | | |
|---|---|---|---|---|---|---|---|---|
| ID | Date | Geo location | Lat, Long | Env biome | Env material | Env feature | Host | Target gene |
| V01 | 07/30/2018 | Australia: New South Wales, Tallaganda forest | 35°30.522'S,149°36.216'E. | temperate forest biome | animal whole gut | deadwood microhabitat | Euperipatoides rowelli (velvet worm) | 16S rRNA V4-V5 region |
| V02 | 07/30/2018 | Australia: New South Wales, Tallaganda forest | 35°30.522'S,149°36.216'E. | temperate forest biome | animal whole gut | deadwood microhabitat | Euperipatoides rowelli (velvet worm) | 16S rRNA V4-V5 region |
| V03 | 07/30/2018 | Australia: New South Wales, Tallaganda forest | 35°26.262'S,149°34.212'E. | temperate forest biome | animal whole gut | deadwood microhabitat | Euperipatoides rowelli (velvet worm) | 16S rRNA V4-V5 region |
| V04 | 07/30/2018 | Australia: New South Wales, Tallaganda forest | 35°26.262'S,149°34.212'E. | temperate forest biome | animal whole gut | deadwood microhabitat | Euperipatoides rowelli (velvet worm) | 16S rRNA V4-V5 region |
| V05 | 07/30/2018 | Australia: New South Wales, Tallaganda forest | 35°26.262'S,149°34.212'E. | temperate forest biome | animal whole gut | deadwood microhabitat | Euperipatoides rowelli (velvet worm) | 16S rRNA V4-V5 region |
| V06 | 07/30/2018 | Australia: New South Wales, Tallaganda forest | 35°26.262'S,149°34.212'E. | temperate forest biome | animal whole gut | deadwood microhabitat | Euperipatoides rowelli (velvet worm) | 16S rRNA V4-V5 region |
| V07 | 07/30/2018 | Australia: New South Wales, Tallaganda forest | 35°30.522'S,149°36.216'E. | temperate forest biome | animal whole gut | deadwood microhabitat | Euperipatoides rowelli (velvet worm) | 16S rRNA V4-V5 region |
| V08 | 07/30/2018 | Australia: New South Wales, Tallaganda forest | 35°30.522'S,149°36.216'E. | temperate forest biome | animal whole gut | deadwood microhabitat | Euperipatoides rowelli (velvet worm) | 16S rRNA V4-V5 region |
| V10 | 07/30/2018 | Australia: New South Wales, Tallaganda forest | 35°30.522'S,149°36.216'E. | temperate forest biome | animal whole gut | deadwood microhabitat | Euperipatoides rowelli (velvet worm) | 16S rRNA V4-V5 region |
| V11 | 07/30/2018 | Australia: New South Wales, Tallaganda forest | 35°30.522'S,149°36.216'E. | temperate forest biome | animal whole gut | deadwood microhabitat | Euperipatoides rowelli (velvet worm) | 16S rRNA V4-V5 region |
| V12 | 07/30/2018 | Australia: New South Wales, Tallaganda forest | 35°30.522'S,149°36.216'E. | temperate forest biome | animal whole gut | deadwood microhabitat | Euperipatoides rowelli (velvet worm) | 16S rRNA V4-V5 region |
| V13 | 07/30/2018 | Australia: New South Wales, Tallaganda forest | 35°30.522'S,149°36.216'E. | temperate forest biome | animal whole gut | deadwood microhabitat | Euperipatoides rowelli (velvet worm) | 16S rRNA V4-V5 region |
| V14 | 07/30/2018 | Australia: New South Wales, Tallaganda forest | 35°30.522'S,149°36.216'E. | temperate forest biome | animal whole gut | deadwood microhabitat | Euperipatoides rowelli (velvet worm) | 16S rRNA V4-V5 region |
| V15 | 07/30/2018 | Australia: New South Wales, Tallaganda forest | 35°30.522'S,149°36.216'E. | temperate forest biome | animal whole gut | deadwood microhabitat | Euperipatoides rowelli (velvet worm) | 16S rRNA V4-V5 region |
| V16 | 07/30/2018 | Australia: New South Wales, Tallaganda forest | 35°59.262'S,149°33.246'E | temperate forest biome | animal whole gut | deadwood microhabitat | Euperipatoides rowelli (velvet worm) | 16S rRNA V4-V5 region |
| V18 | 07/30/2018 | Australia: New South Wales, Tallaganda forest | 35°59.262'S,149°33.246'E | temperate forest biome | animal whole gut | deadwood microhabitat | Euperipatoides rowelli (velvet worm) | 16S rRNA V4-V5 region |
| V20 | 07/30/2018 | Australia: New South Wales, Tallaganda forest | 35°59.262'S,149°33.246'E | temperate forest biome | animal whole gut | deadwood microhabitat | Euperipatoides rowelli (velvet worm) | 16S rRNA V4-V5 region |
| V21 | 07/30/2018 | Australia: New South Wales, Tallaganda forest | 35°59.262'S,149°33.246'E | temperate forest biome | animal whole gut | deadwood microhabitat | Euperipatoides rowelli (velvet worm) | 16S rRNA V4-V5 region |
| V22 | 07/30/2018 | Australia: New South Wales, Tallaganda forest | 35°59.262'S,149°33.246'E | temperate forest biome | animal whole gut | deadwood microhabitat | Euperipatoides rowelli (velvet worm) | 16S rRNA V4-V5 region |
| V23 | 11/07/2018 | Australia: New South Wales, Tallaganda forest | 35°41.49'S,149°33.246'E | temperate forest biome | animal whole gut | deadwood microhabitat | Euperipatoides rowelli (velvet worm) | 16S rRNA V4-V5 region |
| V24 | 11/07/2018 | Australia: New South Wales, Tallaganda forest | 35°41.49'S,149°33.246'E | temperate forest biome | animal whole gut | deadwood microhabitat | Euperipatoides rowelli (velvet worm) | 16S rRNA V4-V5 region |
| V25 | 11/07/2018 | Australia: New South Wales, Tallaganda forest | 35°41.49'S,149°33.246'E | temperate forest biome | animal whole gut | deadwood microhabitat | Euperipatoides rowelli (velvet worm) | 16S rRNA V4-V5 region |
| V26 | 11/07/2018 | Australia: New South Wales, Tallaganda forest | 35°41.49'S,149°33.246'E | temperate forest biome | animal whole gut | deadwood microhabitat | Euperipatoides rowelli (velvet worm) | 16S rRNA V4-V5 region |
| V27 | 11/07/2018 | Australia: New South Wales, Tallaganda forest | 35°41.49'S,149°33.246'E | temperate forest biome | animal whole gut | deadwood microhabitat | Euperipatoides rowelli (velvet worm) | 16S rRNA V4-V5 region |
| V28 | 11/07/2018 | Australia: New South Wales, Tallaganda forest | 35°41.49'S,149°33.246'E | temperate forest biome | animal whole gut | deadwood microhabitat | Euperipatoides rowelli (velvet worm) | 16S rRNA V4-V5 region |
| V29 | 11/07/2018 | Australia: New South Wales, Tallaganda forest | 35°41.49'S,149°33.246'E | temperate forest biome | animal whole gut | deadwood microhabitat | Euperipatoides rowelli (velvet worm) | 16S rRNA V4-V5 region |
| V31 | 11/07/2018 | Australia: New South Wales, Tallaganda forest | 35°41.49'S,149°33.246'E | temperate forest biome | animal whole gut | deadwood microhabitat | Euperipatoides rowelli (velvet worm) | 16S rRNA V4-V5 region |
| V32 | 11/07/2018 | Australia: New South Wales, Tallaganda forest | 35°41.49'S,149°33.246'E | temperate forest biome | animal whole gut | deadwood microhabitat | Euperipatoides rowelli (velvet worm) | 16S rRNA V4-V5 region |
| V38 | 11/07/2018 | Australia: New South Wales, Tallaganda forest | 35°41.49'S,149°33.246'E | temperate forest biome | animal whole gut | deadwood microhabitat | Euperipatoides rowelli (velvet worm) | 16S rRNA V4-V5 region |
| V40 | 11/07/2018 | Australia: New South Wales, Tallaganda forest | 35°41.49'S,149°33.246'E | temperate forest biome | animal whole gut | deadwood microhabitat | Euperipatoides rowelli (velvet worm) | 16S rRNA V4-V5 region |
| V41 | 11/07/2018 | Australia: New South Wales, Tallaganda forest | 35°41.49'S,149°33.246'E | temperate forest biome | animal whole gut | deadwood microhabitat | Euperipatoides rowelli (velvet worm) | 16S rRNA V4-V5 region |
| V42 | 11/07/2018 | Australia: New South Wales, Tallaganda forest | 35°41.49'S,149°33.246'E | temperate forest biome | animal whole gut | deadwood microhabitat | Euperipatoides rowelli (velvet worm) | 16S rRNA V4-V5 region |
| V43 | 11/07/2018 | Australia: New South Wales, Tallaganda forest | 35°41.49'S,149°33.246'E | temperate forest biome | animal whole gut | deadwood microhabitat | Euperipatoides rowelli (velvet worm) | 16S rRNA V4-V5 region |
| V44 | 11/07/2018 | Australia: New South Wales, Tallaganda forest | 35°42.84S,149°33.246'E | temperate forest biome | animal whole gut | deadwood microhabitat | Euperipatoides rowelli (velvet worm) | 16S rRNA V4-V5 region |

**Table S1. Velvet Worms MIxS Metadata**

| ID | Date | Geo location | Lat, Long | Environment<br>Env biome | <br>Env material | <br>Env feature | Host | Target gene |
|---|---|---|---|---|---|---|---|---|
| V45 | 11/07/2018 | Australia: New South Wales, Tallaganda forest | 35º42.84S,149º33.246'E | temperate forest biome | animal whole gut | deadwood microhabitat | Euperipatoides rowelli (velvet worm) | 16S rRNA V4-V5 region |
| V46 | 11/07/2018 | Australia: New South Wales, Tallaganda forest | 35º45.492'S,149º31.062'E | temperate forest biome | animal whole gut | deadwood microhabitat | Euperipatoides rowelli (velvet worm) | 16S rRNA V4-V5 region |
| V48 | 11/07/2018 | Australia: New South Wales, Tallaganda forest | 35º45.492'S,149º31.062'E | temperate forest biome | animal whole gut | deadwood microhabitat | Euperipatoides rowelli (velvet worm) | 16S rRNA V4-V5 region |
| V49 | 11/07/2018 | Australia: New South Wales, Tallaganda forest | 35º45.492'S,149º31.062'E | temperate forest biome | animal whole gut | deadwood microhabitat | Euperipatoides rowelli (velvet worm) | 16S rRNA V4-V5 region |
| V50 | 11/07/2018 | Australia: New South Wales, Tallaganda forest | 35º45.492'S,149º31.062'E | temperate forest biome | animal whole gut | deadwood microhabitat | Euperipatoides rowelli (velvet worm) | 16S rRNA V4-V5 region |
| V52 | 11/07/2018 | Australia: New South Wales, Tallaganda forest | 35º45.492'S,149º31.062'E | temperate forest biome | animal whole gut | deadwood microhabitat | Euperipatoides rowelli (velvet worm) | 16S rRNA V4-V5 region |
| V54 | 11/07/2018 | Australia: New South Wales, Tallaganda forest | 35º45.492'S,149º31.062'E | temperate forest biome | animal whole gut | deadwood microhabitat | Euperipatoides rowelli (velvet worm) | 16S rRNA V4-V5 region |
| V56 | 11/07/2018 | Australia: New South Wales, Tallaganda forest | 35º45.492'S,149º31.062'E | temperate forest biome | animal whole gut | deadwood microhabitat | Euperipatoides rowelli (velvet worm) | 16S rRNA V4-V5 region |
| V57 | 11/07/2018 | Australia: New South Wales, Tallaganda forest | 35º45.492'S,149º31.062'E | temperate forest biome | animal whole gut | deadwood microhabitat | Euperipatoides rowelli (velvet worm) | 16S rRNA V4-V5 region |
| V59 | 11/07/2018 | Australia: New South Wales, Tallaganda forest | 35º44.818'S,149º31.818'E | temperate forest biome | animal whole gut | deadwood microhabitat | Euperipatoides rowelli (velvet worm) | 16S rRNA V4-V5 region |
| V61 | 07/31/2019 | Australia: New South Wales, Tallaganda forest | 35º45.492'S,149º31.062'E | temperate forest biome | animal whole gut | deadwood microhabitat | Euperipatoides rowelli (velvet worm) | 16S rRNA V4-V5 region |
| V62 | 07/31/2019 | Australia: New South Wales, Tallaganda forest | 35º45.492'S,149º31.062'E | temperate forest biome | animal whole gut | deadwood microhabitat | Euperipatoides rowelli (velvet worm) | 16S rRNA V4-V5 region |
| V63 | 07/31/2019 | Australia: New South Wales, Tallaganda forest | 35º45.492'S,149º31.062'E | temperate forest biome | animal whole gut | deadwood microhabitat | Euperipatoides rowelli (velvet worm) | 16S rRNA V4-V5 region |
| V64 | 07/31/2019 | Australia: New South Wales, Tallaganda forest | 35º44.818'S,149º31.818'E | temperate forest biome | animal whole gut | deadwood microhabitat | Euperipatoides rowelli (velvet worm) | 16S rRNA V4-V5 region |
| V66 | 07/31/2019 | Australia: New South Wales, Tallaganda forest | 35º44.818'S,149º31.818'E | temperate forest biome | animal whole gut | deadwood microhabitat | Euperipatoides rowelli (velvet worm) | 16S rRNA V4-V5 region |
| V67 | 07/31/2019 | Australia: New South Wales, Tallaganda forest | 35º44.818'S,149º31.818'E | temperate forest biome | animal whole gut | deadwood microhabitat | Euperipatoides rowelli (velvet worm) | 16S rRNA V4-V5 region |
| V68 | 07/31/2019 | Australia: New South Wales, Tallaganda forest | 35º44.818'S,149º31.818'E | temperate forest biome | animal whole gut | deadwood microhabitat | Euperipatoides rowelli (velvet worm) | 16S rRNA V4-V5 region |
| V69 | 07/31/2019 | Australia: New South Wales, Tallaganda forest | 35º44.818'S,149º31.818'E | temperate forest biome | animal whole gut | deadwood microhabitat | Euperipatoides rowelli (velvet worm) | 16S rRNA V4-V5 region |
| V72 | 07/31/2019 | Australia: New South Wales, Tallaganda forest | 35º44.818'S,149º31.818'E | temperate forest biome | animal whole gut | deadwood microhabitat | Euperipatoides rowelli (velvet worm) | 16S rRNA V4-V5 region |
| V73 | 07/31/2019 | Australia: New South Wales, Tallaganda forest | 35º44.818'S,149º31.818'E | temperate forest biome | animal whole gut | deadwood microhabitat | Euperipatoides rowelli (velvet worm) | 16S rRNA V4-V5 region |
| V74 | 07/31/2019 | Australia: New South Wales, Tallaganda forest | 35º44.818'S,149º31.818'E | temperate forest biome | animal whole gut | deadwood microhabitat | Euperipatoides rowelli (velvet worm) | 16S rRNA V4-V5 region |
| V75 | 07/31/2019 | Australia: New South Wales, Tallaganda forest | 35º44.818'S,149º31.818'E | temperate forest biome | animal whole gut | deadwood microhabitat | Euperipatoides rowelli (velvet worm) | 16S rRNA V4-V5 region |
| V76 | 07/31/2019 | Australia: New South Wales, Tallaganda forest | 35º44.818'S,149º31.818'E | temperate forest biome | animal whole gut | deadwood microhabitat | Euperipatoides rowelli (velvet worm) | 16S rRNA V4-V5 region |
| V77 | 07/31/2019 | Australia: New South Wales, Tallaganda forest | 35º44.818'S,149º31.818'E | temperate forest biome | animal whole gut | deadwood microhabitat | Euperipatoides rowelli (velvet worm) | 16S rRNA V4-V5 region |
| V79 | 12/18/2020 | Australia: New South Wales, Tallaganda forest | 35º45.492'S,149º31.062'E | temperate forest biome | animal whole gut | deadwood microhabitat | Euperipatoides rowelli (velvet worm) | 16S rRNA V4-V5 region |
| V83 | 12/18/2020 | Australia: New South Wales, Tallaganda forest | 35º44.818'S,149º31.818'E | temperate forest biome | animal whole gut | deadwood microhabitat | Euperipatoides rowelli (velvet worm) | 16S rRNA V4-V5 region |
| V87 | 12/18/2020 | Australia: New South Wales, Tallaganda forest | 35º44.818'S,149º31.818'E | temperate forest biome | animal whole gut | deadwood microhabitat | Euperipatoides rowelli (velvet worm) | 16S rRNA V4-V5 region |
| V90 | 01/19/2021 | Australia: New South Wales, Tallaganda forest | 35º59.4453'S,149º32.891'E | temperate forest biome | animal whole gut | deadwood microhabitat | Euperipatoides rowelli (velvet worm) | 16S rRNA V4-V5 region |
| V93 | 01/19/2021 | Australia: New South Wales, Tallaganda forest | 35º45.492'S,149º31.062'E | temperate forest biome | animal whole gut | deadwood microhabitat | Euperipatoides rowelli (velvet worm) | 16S rRNA V4-V5 region |
| V104 | 01/19/2021 | Australia: New South Wales, Tallaganda forest | 35º59.4453'S,149º32.891'E | temperate forest biome | animal whole gut | deadwood microhabitat | Euperipatoides rowelli (velvet worm) | 16S rRNA V4-V5 region |
| V105 | 01/19/2021 | Australia: New South Wales, Tallaganda forest | 35º59.4453'S,149º32.891'E | temperate forest biome | animal whole gut | deadwood microhabitat | Euperipatoides rowelli (velvet worm) | 16S rRNA V4-V5 region |

### Table S2. QIIME2 Processing Summary

Counts and percentages across filtering and denoising steps

| Sample ID | Input reads | Filtered | % passed filter | Denoised | Merged | % merged | Non-chimeric | % non-chimeric |
|---|---|---|---|---|---|---|---|---|
| V1 | 20 | 17 | 85% | 15 | 6 | 30% | 6 | 30% |
| V2 | 1,821 | 1,717 | 94.29% | 1,709 | 1,496 | 82.15% | 1,496 | 82.15% |
| V3 | 4,983 | 4,699 | 94.3% | 4,664 | 4,297 | 86.23% | 4,283 | 85.95% |
| V4 | 4,424 | 1,353 | 30.58% | 1,339 | 1,150 | 25.99% | 1,150 | 25.99% |
| V5 | 17 | 13 | 76.47% | 5 | 5 | 29.41% | 5 | 29.41% |
| V6 | 45 | 41 | 91.11% | 37 | 37 | 82.22% | 37 | 82.22% |
| V7 | 87,121 | 82,881 | 95.13% | 82,264 | 75,458 | 86.61% | 75,429 | 86.58% |
| V8 | 778 | 71 | 9.13% | 46 | 15 | 1.93% | 15 | 1.93% |
| V10 | 1,486 | 65 | 4.37% | 60 | 60 | 4.04% | 60 | 4.04% |
| V11 | 102 | 95 | 93.14% | 69 | 56 | 54.9% | 56 | 54.9% |
| V12 | 3,217 | 597 | 18.56% | 592 | 564 | 17.53% | 564 | 17.53% |
| V13 | 797 | 167 | 20.95% | 167 | 115 | 14.43% | 115 | 14.43% |
| V14 | 95 | 88 | 92.63% | 51 | 27 | 28.42% | 27 | 28.42% |
| V15 | 42 | 40 | 95.24% | 30 | 30 | 71.43% | 30 | 71.43% |
| V16 | 621,594 | 203,618 | 32.76% | 203,058 | 148,207 | 23.84% | 147,210 | 23.68% |
| V18 | 43,509 | 41,030 | 94.3% | 40,855 | 40,080 | 92.12% | 39,847 | 91.58% |
| V20 | 3,090 | 2,895 | 93.69% | 2,883 | 2,619 | 84.76% | 2,607 | 84.37% |
| V21 | 9,291 | 663 | 7.14% | 647 | 603 | 6.49% | 603 | 6.49% |
| V22 | 7,491 | 1,195 | 15.95% | 1,183 | 1,063 | 14.19% | 1,063 | 14.19% |
| V24 | 25,255 | 23,952 | 94.84% | 23,942 | 23,052 | 91.28% | 23,052 | 91.28% |
| V25 | 102 | 88 | 86.27% | 73 | 57 | 55.88% | 57 | 55.88% |
| V26 | 2,236 | 2,076 | 92.84% | 2,063 | 1,986 | 88.82% | 1,986 | 88.82% |
| V27 | 143 | 128 | 89.51% | 79 | 46 | 32.17% | 46 | 32.17% |
| V28 | 21,268 | 1,212 | 5.7% | 1,210 | 645 | 3.03% | 645 | 3.03% |
| V29 | 487 | 92 | 18.89% | 84 | 82 | 16.84% | 82 | 16.84% |
| V31 | 5,542 | 583 | 10.52% | 577 | 244 | 4.4% | 234 | 4.22% |
| V32 | 570 | 66 | 11.58% | 60 | 33 | 5.79% | 33 | 5.79% |
| V38 | 237 | 218 | 91.98% | 216 | 213 | 89.87% | 213 | 89.87% |
| V40 | 1,844 | 1,760 | 95.44% | 1,679 | 1,664 | 90.24% | 1,664 | 90.24% |
| V41 | 8,719 | 8,179 | 93.81% | 7,962 | 7,802 | 89.48% | 7,790 | 89.35% |
| V42 | 74 | 70 | 94.59% | 38 | 20 | 27.03% | 20 | 27.03% |
| V43 | 276 | 262 | 94.93% | 198 | 181 | 65.58% | 181 | 65.58% |
| V44 | 460 | 384 | 83.48% | 273 | 213 | 46.3% | 202 | 43.91% |
| V45 | 1,393 | 1,301 | 93.4% | 1,294 | 1,292 | 92.75% | 1,292 | 92.75% |
| V46 | 795 | 730 | 91.82% | 727 | 723 | 90.94% | 723 | 90.94% |
| V48 | 100 | 98 | 98% | 86 | 85 | 85% | 85 | 85% |
| V49 | 78 | 72 | 92.31% | 62 | 62 | 79.49% | 62 | 79.49% |
| V50 | 34 | 29 | 85.29% | 17 | 17 | 50% | 17 | 50% |
| V52 | 824 | 769 | 93.33% | 668 | 666 | 80.83% | 666 | 80.83% |
| V54 | 538 | 48 | 8.92% | 27 | 17 | 3.16% | 17 | 3.16% |
| V56 | 21 | 19 | 90.48% | 9 | 9 | 42.86% | 9 | 42.86% |
| V57 | 9,750 | 9,041 | 92.73% | 8,560 | 7,764 | 79.63% | 7,346 | 75.34% |
| V59 | 11,922 | 11,288 | 94.68% | 11,205 | 11,087 | 93% | 11,067 | 92.83% |

**Source:** QIIME2 pipeline outputs for velvet worm dataset.

**Table S2. QIIME2 Processing Summary**

Counts and percentages across filtering and denoising steps

| Sample ID | Input reads | Filtered | % passed filter | Denoised | Merged | % merged | Non-chimeric | % non-chimeric |
|---|---|---|---|---|---|---|---|---|
| V61 | 132 | 121 | 91.67% | 94 | 85 | 64.39% | 85 | 64.39% |
| V62 | 341 | 315 | 92.38% | 236 | 187 | 54.84% | 187 | 54.84% |
| V63 | 616 | 599 | 97.24% | 550 | 517 | 83.93% | 517 | 83.93% |
| V64 | 242 | 222 | 91.74% | 149 | 137 | 56.61% | 137 | 56.61% |
| V66 | 146 | 92 | 63.01% | 37 | 24 | 16.44% | 24 | 16.44% |
| V67 | 2,223,558 | 1,095,911 | 49.29% | 1,094,217 | 1,073,861 | 48.29% | 188,501 | 8.48% |
| V68 | 92 | 87 | 94.57% | 62 | 48 | 52.17% | 48 | 52.17% |
| V69 | 78 | 44 | 56.41% | 33 | 28 | 35.9% | 28 | 35.9% |
| V72 | 1,142 | 978 | 85.64% | 723 | 564 | 49.39% | 552 | 48.34% |
| V73 | 20,694 | 20,091 | 97.09% | 20,089 | 19,983 | 96.56% | 19,983 | 96.56% |
| V74 | 670 | 559 | 83.43% | 325 | 193 | 28.81% | 193 | 28.81% |
| V75 | 1,114 | 1,005 | 90.22% | 983 | 979 | 87.88% | 979 | 87.88% |
| V76 | 16,369 | 15,731 | 96.1% | 15,314 | 15,223 | 93% | 15,223 | 93% |
| V77 | 27,614 | 26,627 | 96.43% | 25,372 | 21,995 | 79.65% | 19,909 | 72.1% |
| V79 | 171 | 154 | 90.06% | 145 | 137 | 80.12% | 137 | 80.12% |
| V83 | 783 | 714 | 91.19% | 146 | 138 | 17.62% | 138 | 17.62% |
| V87 | 216 | 197 | 91.2% | 183 | 165 | 76.39% | 153 | 70.83% |
| V90 | 1,425 | 786 | 55.16% | 771 | 771 | 54.11% | 771 | 54.11% |
| V93 | 94,626 | 53,246 | 56.27% | 53,009 | 52,702 | 55.7% | 28,258 | 29.86% |
| V104 | 34,219 | 19,405 | 56.71% | 19,284 | 19,273 | 56.32% | 16,755 | 48.96% |
| V105 | 78 | 56 | 71.79% | 14 | 9 | 11.54% | 9 | 11.54% |

**Source:** QIIME2 pipeline outputs for velvet worm dataset.

**Table S3.Velvet Worm Gut Microbial Alpha Diversity Measures**

Includes gradient, site description and crude whole velvet worm gut DNA concentration

| ID | Site | Gradient | Locality | PD | SR | Shan | DNA Concentration |
|---|---|---|---|---|---|---|---|
| V1 | S2 | N | Lowden Park | 1.68 | 2.00 | 0.64 | 63.30 |
| V2 | S2 | N | Lowden Park | 1.59 | 2.00 | 0.03 | 14.00 |
| V3 | S1 | N | Mulloon Flat Camping area | 45.35 | 82.00 | 2.84 | 15.50 |
| V4 | S1 | N | Mulloon Flat Camping area | 5.68 | 7.00 | 0.57 | 14.20 |
| V5 | S1 | N | Mulloon Flat Camping area | 4.41 | 6.00 | 1.12 | 15.30 |
| V6 | S1 | N | Mulloon Flat Camping area | 3.07 | 4.00 | 1.08 | 15.80 |
| V7 | S2 | N | Lowden Park | 4.71 | 60.00 | 2.45 | 11.30 |
| V8 | S2 | N | Lowden Park | 2.05 | 2.00 | 0.58 | 3.60 |
| V10 | S2 | N | Lowden Park | 6.26 | 8.00 | 0.72 | 3.64 |
| V11 | S2 | N | Lowden Park | 16.73 | 32.00 | 2.81 | 24.00 |
| V12 | S2 | N | Lowden Park | 5.00 | 8.00 | 1.12 | 5.50 |
| V13 | S2 | N | Lowden Park | 5.25 | 5.00 | 1.12 | 0.25 |
| V14 | S2 | N | Lowden Park | 3.71 | 4.00 | 1.11 | 0.10 |
| V15 | S2 | N | Lowden Park | 1.70 | 2.00 | 0.58 | 3.32 |
| V16 | S3 | S | Badja Road, Badja A | 8.31 | 13.00 | 0.21 | 1.32 |
| V18 | S3 | S | Badja Road, Badja A | 35.45 | 77.00 | 0.87 | 15.10 |
| V20 | S3 | S | Badja Road, Badja A | 10.98 | 16.00 | 1.12 | 4.28 |
| V21 | S3 | S | Badja Road, Badja A | 4.84 | 8.00 | 0.76 | 3.36 |
| V22 | S3 | S | Badja Road, Badja A | 4.98 | 4.00 | 0.55 | 1.57 |
| V24 | S4 | N | Kindervale | 3.59 | 6.00 | 0.06 | 3.14 |
| V25 | S4 | N | Kindervale | 4.55 | 6.00 | 1.68 | 3.78 |
| V26 | S4 | N | Kindervale | 17.33 | 34.00 | 1.91 | 2.40 |
| V27 | S4 | N | Kindervale | 3.97 | 6.00 | 1.73 | 0.76 |
| V28 | S4 | N | Kindervale | 2.74 | 3.00 | 0.11 | 0.80 |
| V29 | S4 | N | Kindervale | 5.33 | 7.00 | 1.47 | 0.77 |
| V31 | S4 | N | Kindervale | 1.86 | 2.00 | 0.13 | 0.84 |
| V32 | S4 | N | Kindervale | 2.32 | 3.00 | 0.98 | 0.80 |
| V38 | S4 | N | Kindervale | 4.50 | 6.00 | 1.61 | 0.88 |
| V40 | S4 | N | Kindervale | 7.99 | 10.00 | 0.42 | 23.20 |
| V41 | S4 | N | Kindervale | 32.06 | 67.00 | 1.24 | 0.84 |
| V42 | S4 | N | Kindervale | 2.77 | 3.00 | 0.74 | 5.66 |
| V43 | S4 | N | Kindervale | 9.71 | 12.00 | 1.97 | 1.23 |
| V44 | S5 | MW | Cowangerong Firetrail | 14.12 | 15.00 | 2.37 | 2.42 |

**Source:Alpha Diversity Metrics: PD, FAith's Phylogenetic Diversity; SR, Species Richness; Shan, Shannon's Index**

### Table S3.Velvet Worm Gut Microbial Alpha Diversity Measures

Includes gradient, site description and crude whole velvet worm gut DNA concentration

| ID | Site | Gradient | Locality | PD | SR | Shan | DNA Concentration |
|---|---|---|---|---|---|---|---|
| V45 | S5 | MW | Cowangerong Firetrail | 6.71 | 10.00 | 1.05 | 2.86 |
| V46 | S6 | MW | Anembo | 5.56 | 8.00 | 1.36 | 0.87 |
| V48 | S6 | MW | Anembo | 2.12 | 2.00 | 0.11 | 36.20 |
| V49 | S6 | MW | Anembo | 3.94 | 5.00 | 1.46 | 37.40 |
| V50 | S6 | MW | Anembo | 2.98 | 4.00 | 1.38 | 0.80 |
| V52 | S6 | MW | Anembo | 13.85 | 17.00 | 1.67 | 4.16 |
| V54 | S6 | MW | Anembo | 6.63 | 8.00 | 1.80 | 9.90 |
| V56 | S6 | MW | Anembo | 1.80 | 2.00 | 0.69 | 8.26 |
| V57 | S6 | MW | Anembo | 37.85 | 158.00 | 4.55 | 6.96 |
| V59 | S7a | MW | Hereford Hall | 17.91 | 32.00 | 0.73 | 5.56 |
| V61 | S6 | MW | Anembo | 3.97 | 5.00 | 0.73 | 1.87 |
| V62 | S6 | MW | Anembo | 9.86 | 16.00 | 2.40 | 36.20 |
| V63 | S6 | MW | Anembo | 5.31 | 6.00 | 0.29 | 13.80 |
| V64 | S7a | MW | Hereford Hall | 7.62 | 10.00 | 1.45 | 29.20 |
| V66 | S7a | MW | Hereford Hall | 6.49 | 5.00 | 1.57 | 31.20 |
| V67 | S7a | MW | Hereford Hall | 4.19 | 52.00 | 2.51 | 25.20 |
| V68 | S7a | MW | Hereford Hall | 3.22 | 5.00 | 1.15 | 25.40 |
| V69 | S7a | MW | Hereford Hall | 2.48 | 3.00 | 0.79 | 33.40 |
| V72 | S7a | MW | Hereford Hall | 4.89 | 15.00 | 2.06 | 2.00 |
| V73 | S7a | MW | Hereford Hall | 0.18 | 3.00 | 0.04 | 1.77 |
| V74 | S7a | MW | Hereford Hall | 7.23 | 9.00 | 1.53 | 2.09 |
| V75 | S7a | MW | Hereford Hall | 5.54 | 6.00 | 0.58 | 2.80 |
| V76 | S7a | MW | Hereford Hall | 3.17 | 9.00 | 0.09 | 4.48 |
| V77 | S7a | MW | Hereford Hall | 96.78 | 497.00 | 5.72 | 3.60 |
| V79 | S6 | MW | Anembo | 8.04 | 9.00 | 1.85 | 29.80 |
| V83 | S7a | MW | Hereford Hall | 4.55 | 5.00 | 0.91 | 35.35 |
| V87 | S7a | MW | Hereford Hall | 4.41 | 6.00 | 1.03 | 6.69 |
| V90 | S8a | MW | Badja Road, Badja B | 3.94 | 4.00 | 0.15 | 60.20 |
| V93 | S6 | MW | Anembo | 3.19 | 25.00 | 2.40 | 9.56 |
| V104 | S8a | MW | Badja Road, Badja B | 3.12 | 12.00 | 2.05 | 29.40 |
| V105 | S8a | MW | Badja Road, Badja B | 1.54 | 2.00 | 0.69 | 12.10 |

**Source:Alpha Diversity Metrics: PD, FAith's Phylogenetic Diversity; SR, Species Richness; Shan, Shannon's Index**

### Table S4.Velvet Worm Gut Microbial Beta Diversity Statistical Analysis

PERMANOVA Results by Site

| Site | Degrees of Freedom | Sum of Squares | Mean Squares | F Value | R-squared |
|---|---|---|---|---|---|
| **Euclidean (CLR)** | 7.00 | 9140.801 | 1305.829. | 1.11 | 0.11 |
| **WUNIFrac** | 7.00 | 0.3160279 | 0.04514684 | 0.70 | 0.07 |

This table shows PERMANOVA results for velvet worm gut microbiomes using Euclidean (CLR) and Unweig distances. Neither metric detected significant site-level differences ($p > 0.05$), suggesting weak phylogen compositional structure across sites.

**Table S5.Permutation test for homogeneity of multivariate dispersions Results by Site**

For each distance, the test evaluates whether microbial community variation (dispersion) significantly differs across fo

| Site | Degrees of Freedom | Sum of Squares | Mean Squares | F Value | Permutations |
|---|---|---|---|---|---|
| **Groups :Euclidean** | 7.00 | 836.40 | 119.48 | 0.36 | 999.00 |
| **Groups:WUNIFrac** | 7.00 | 0.13 | 0.02 | 0.87 | 999.00 |

Both Euclidean ($p = 0.905$) and Weighted UniFrac ($p = 0.538$) results are non-significant, indicating no evi unequal dispersion across sites, and supporting the assumption of homogeneity for PERMANOVA.

| Table 6A.Core gut microbiome of velvet worms based on 30 % prevalence | | | | | | | | |
|---|---|---|---|---|---|---|---|---|
| **Phylum** | **S1** | **S2** | **S3** | **S4** | **S5** | **S6** | **S7a** | **S8a** |
| Actinobacteriota | 1 | 1 | 1 | 1 | 1 | 1 | 1 | 0 |
| Firmicutes | 1 | 1 | 1 | 1 | 1 | 1 | 1 | 1 |
| Bacteroidota | 1 | 1 | 1 | 1 | 1 | 0 | 0 | 1 |
| Proteobacteria | 1 | 1 | 1 | 1 | 1 | 1 | 1 | 1 |
| Spirochaetota | 1 | 0 | 0 | 0 | 1 | 0 | 1 | 0 |
| Acidobacteriota | 0 | 0 | 1 | 0 | 0 | 0 | 0 | 1 |
| Verrucomicrobiota | 0 | 0 | 1 | 0 | 0 | 0 | 0 | 0 |
| Bdellovibrionota | 0 | 0 | 0 | 0 | 1 | 0 | 0 | 0 |
| Crenarchaeota | 0 | 0 | 0 | 0 | 1 | 0 | 0 | 0 |
| Myxococcota | 0 | 0 | 0 | 0 | 1 | 0 | 0 | 0 |
| Thermoplasmatota | 0 | 0 | 0 | 0 | 1 | 0 | 0 | 0 |
| Planctomycetota | 0 | 0 | 0 | 0 | 0 | 1 | 0 | 0 |

This table lists bacterial taxa identified as part of the core microbiome across velvet worm gut samples. These core taxa represent consistently associated microbial members across sampling sites.Detection threshold = 0.001

### Table 6B.Core gut microbiome of velvet worms based on 50 % prevalence

| Genus | S1 | S2 | S3 | S4 | S5 | S6 | S7a | S8a |
|---|---|---|---|---|---|---|---|---|
| Cutibacterium | 1 | 0 | 0 | 0 | 0 | 0 | 0 | 0 |
| Sphingomonas | 0 | 0 | 1 | 0 | 0 | 0 | 0 | 0 |
| Spiroplasma | 0 | 0 | 1 | 0 | 1 | 0 | 0 | 1 |
| Blattabacterium | 0 | 0 | 1 | 0 | 0 | 0 | 0 | 0 |
| Bradyrhizobium | 0 | 0 | 1 | 0 | 0 | 0 | 0 | 0 |
| Enhydrobacter | 0 | 0 | 1 | 0 | 0 | 0 | 0 | 0 |
| Ac37b | 0 | 0 | 1 | 1 | 0 | 1 | 1 | 0 |
| Mycobacterium | 0 | 0 | 1 | 0 | 0 | 0 | 0 | 0 |
| Diplorickettsia | 0 | 0 | 0 | 0 | 0 | 0 | 0 | 1 |

This table lists bacterial taxa identified as part of the core microbiome across velvet worm gut samples. These core taxa represent consistently associated microbial members across sampling sites.Detection threshold = 0.001

### Table 7.Core gut microbiome of velvet worms based on 40 % prevalence

| Phylum | S1 | S2 | S3 | S4 | S5 | S6 | S7a | S8a |
|---|---|---|---|---|---|---|---|---|
| Actinobacteriota | 1 | 1 | 1 | 1 | 1 | 1 | 1 | 0 |
| Firmicutes | 1 | 1 | 1 | 1 | 1 | 1 | 0 | 1 |
| Proteobacteria | 1 | 1 | 1 | 1 | 1 | 1 | 1 | 1 |
| Bacteroidota | 1 | 0 | 1 | 0 | 1 | 0 | 0 | 0 |
| Spirochaetota | 1 | 0 | 0 | 0 | 1 | 0 | 0 | 0 |
| Acidobacteriota | 0 | 0 | 1 | 0 | 0 | 0 | 0 | 0 |
| Bdellovibrionota | 0 | 0 | 0 | 0 | 1 | 0 | 0 | 0 |
| Crenarchaeota | 0 | 0 | 0 | 0 | 1 | 0 | 0 | 0 |
| Myxococcota | 0 | 0 | 0 | 0 | 1 | 0 | 0 | 0 |
| Thermoplasmatota | 0 | 0 | 0 | 0 | 1 | 0 | 0 | 0 |

This table lists bacterial taxa identified as part of the core microbiome across velvet worm gut samples. These core taxa represent consistently associated microbial members across sampling sites.Detection threshold = 0.001

### Table 7B.Core gut microbiome of velvet worms based on 40 % prevalence

| Genus | S1 | S2 | S3 | S4 | S5 | S6 | S7a | S8a |
|---|---|---|---|---|---|---|---|---|
| Cutibacterium | 1 | 0 | 0 | 0 | 1 | 0 | 0 | 0 |
| Spiroplasma | 1 | 0 | 1 | 0 | 1 | 0 | 0 | 1 |
| Borrelia | 1 | 0 | 0 | 0 | 1 | 0 | 0 | 0 |
| Ac37b | 1 | 0 | 1 | 1 | 1 | 1 | 1 | 0 |
| Mycobacterium | 1 | 0 | 1 | 0 | 0 | 0 | 0 | 0 |
| Pseudomonas | 1 | 0 | 0 | 0 | 1 | 0 | 0 | 0 |
| Nocardioides | 1 | 0 | 0 | 0 | 0 | 0 | 0 | 0 |
| Sphingomonas | 0 | 0 | 1 | 0 | 1 | 0 | 0 | 0 |
| Blattabacterium | 0 | 0 | 1 | 0 | 0 | 0 | 0 | 0 |
| Bradyrhizobium | 0 | 0 | 1 | 0 | 1 | 0 | 0 | 0 |
| Enhydrobacter | 0 | 0 | 1 | 0 | 0 | 0 | 0 | 0 |
| Rickettsiella | 0 | 0 | 0 | 0 | 1 | 0 | 0 | 0 |
| Haliangium | 0 | 0 | 0 | 0 | 1 | 0 | 0 | 0 |
| Enterococcus | 0 | 0 | 0 | 0 | 1 | 0 | 0 | 0 |
| Kitasatospora | 0 | 0 | 0 | 0 | 1 | 0 | 0 | 0 |
| Streptococcus | 0 | 0 | 0 | 0 | 1 | 0 | 0 | 0 |
| Candidatus | 0 | 0 | 0 | 0 | 1 | 0 | 0 | 0 |
| Brevundimonas | 0 | 0 | 0 | 0 | 1 | 0 | 0 | 0 |
| Escherichia-Shigella | 0 | 0 | 0 | 0 | 1 | 0 | 0 | 0 |
| Ralstonia | 0 | 0 | 0 | 0 | 1 | 0 | 0 | 0 |
| Staphylococcus | 0 | 0 | 0 | 0 | 1 | 0 | 0 | 0 |
| Methylobacterium-Methylorubrum | 0 | 0 | 0 | 0 | 1 | 0 | 0 | 0 |
| Chryseobacterium | 0 | 0 | 0 | 0 | 1 | 0 | 0 | 0 |

This table lists bacterial taxa identified as part of the core microbiome across velvet worm gut samples. These core taxa represent consistently associated microbial members across sampling sites.Detection threshold = 0.001

### Table 7B.Core gut microbiome of velvet worms based on 40 % prevalence

| Genus | S1 | S2 | S3 | S4 | S5 | S6 | S7a | S8a |
|---|---|---|---|---|---|---|---|---|
| Phenylobacterium | 0 | 0 | 0 | 0 | 1 | 0 | 0 | 0 |
| Diplorickettsia | 0 | 0 | 0 | 0 | 0 | 0 | 0 | 1 |

This table lists bacterial taxa identified as part of the core microbiome across velvet worm gut samples. These core taxa represent consistently associated microbial members across sampling sites.Detection threshold = 0.001

Table 8.Core gut microbiome of velvet worms based on 50 % prevalence

| Phylum | S1 | S2 | S3 | S4 | S5 | S6 | S7a | S8a |
|---|---|---|---|---|---|---|---|---|
| Firmicutes | 0 | 1 | 1 | 1 | 1 | 1 | 0 | 1 |
| Proteobacteria | 1 | 1 | 1 | 1 | 1 | 1 | 1 | 1 |
| Actinobacteriota | 1 | 0 | 1 | 0 | 1 | 1 | 0 | 0 |
| Bacteroidota | 0 | 0 | 1 | 0 | 0 | 0 | 0 | 0 |
| Acidobacteriota | 0 | 0 | 1 | 0 | 0 | 0 | 0 | 0 |

This table lists bacterial taxa identified as part of the core microbiome across velvet worm gut samples. These core taxa represent consistently associated microbial members across sampling sites.Detection threshold = 0.001

### Table 9: Faith’s PD Summary Linear Mixed Model

Model Fit by REML, with Satterthwaite's Method

| | Fixed Effects | | | | |
|---|---|---|---|---|---|
| **Term** | **Estimate** | **Std. Error** | **df** | **t value** | **Pr(>\|t\|)** |
| (Intercept) | 2.0393 | 0.7744 | 56 | 2.6340 | 0.0109 |
| S2 | −0.7385 | 0.9939 | 56 | −0.7430 | 0.4606 |
| S3 | 0.2133 | 1.0737 | 56 | 0.1990 | 0.8433 |
| S4 | −0.3849 | 0.9830 | 56 | −0.3920 | 0.6969 |
| S5 | 0.2363 | 1.1964 | 56 | 0.1980 | 0.8441 |
| S6 | −0.3279 | 1.0191 | 56 | −0.3220 | 0.7488 |
| S7a | −0.4264 | 1.0166 | 56 | −0.4190 | 0.6765 |
| S8a | −1.0595 | 1.1299 | 56 | −0.9380 | 0.3524 |

Note: REML criterion at convergence: 169.6. Signif. codes: 0 ‘***’ 0.001 ‘**’ 0.01 ‘*’ 0.05 ‘.’ 0.1 ‘ ’ 1

### Table S10: Shannon Summary Linear Mixed Model

Model Fit by REML, with Satterthwaite's Method

| | Fixed Effects | | | | |
|---|---|---|---|---|---|
| **Term** | **Estimate** | **Std. Error** | **df** | **t value** | **Pr(>\|t\|)** |
| (Intercept) | 0.1651 | 0.6485 | 56 | 0.2550 | 0.8000 |
| S2 | −0.4759 | 0.7723 | 56 | −0.6160 | 0.5400 |
| S3 | −0.6585 | 0.8816 | 56 | −0.7470 | 0.4580 |
| S4 | −0.5324 | 0.7529 | 56 | −0.7070 | 0.4820 |
| S5 | 0.2882 | 1.0773 | 56 | 0.2680 | 0.7900 |
| S6 | −0.0227 | 0.7874 | 56 | −0.0290 | 0.9770 |
| S7a | −0.3071 | 0.7829 | 56 | −0.3920 | 0.6960 |
| S8a | −0.6816 | 0.9735 | 56 | −0.7000 | 0.4870 |

Note: REML criterion at convergence: 187.5. Signif. codes: 0 ‘***’ 0.001 ‘**’ 0.01 ‘*’ 0.05 ‘.’ 0.1 ‘ ’ 1

### Table S11: Species Richness Summary Linear Mixed Model

Model Fit by REML, with Satterthwaite's Method

| | Fixed Effects | | | | |
|---|---|---|---|---|---|
| **Term** | **Estimate** | **Std. Error** | **df** | **t value** | **Pr(>\|t\|)** |
| (Intercept) | 2.38267 | 0.70433 | 56 | 3.38300 | 0.00131 |
| SiteS2 | −0.63394 | 0.83394 | 56 | −0.76000 | 0.45034 |
| SiteS3 | 0.24675 | 0.96069 | 56 | 0.25700 | 0.79824 |
| SiteS4 | −0.41186 | 0.81456 | 56 | −0.50600 | 0.61510 |
| SiteS5 | 0.12265 | 1.15683 | 56 | 0.10600 | 0.91595 |
| SiteS6 | −0.22678 | 0.86749 | 56 | −0.26100 | 0.79473 |
| SiteS7a | 0.02017 | 0.86309 | 56 | 0.02300 | 0.98144 |
| SiteS8a | −0.86122 | 1.05239 | 56 | −0.81800 | 0.41663 |

Note: REML criterion at convergence: 192. Signif. codes: 0 '***' 0.001 '**' 0.01 '*' 0.05 '.' 0.1 ' ' 1

**Table 12.Significant Differential Abundance of Genera Across Site Comparisons**

*Filtered for Adjusted p-value < 0.05*

| Genus | Phylum | **Log2 Fold Change** | p-value | **Adjusted p-value** | Site Comparison |
|---|---|---|---|---|---|
| *Spiroplasma* | Firmicutes | 10.912 | 0.000 | 0.000 | S3 vs. S8a |
| *Spiroplasma* | Firmicutes | 10.912 | 0.000 | 0.000 | S3 vs. S6 |
| *Spiroplasma* | Firmicutes | 10.911 | 0.000 | 0.000 | S3 vs. S5 |
| *Ac37b* | Proteobacteria | 9.949 | 0.000 | 0.000 | S4 vs. S8a |
| *Spiroplasma* | Firmicutes | 9.862 | 0.000 | 0.000 | S3 vs. S7a |
| *Spiroplasma* | Firmicutes | 8.986 | 0.000 | 0.000 | S1 vs. S8a |
| *Spiroplasma* | Firmicutes | 8.986 | 0.000 | 0.000 | S1 vs. S6 |
| *Spiroplasma* | Firmicutes | 8.986 | 0.000 | 0.005 | S1 vs. S5 |
| *Ac37b* | Proteobacteria | 8.968 | 0.000 | 0.001 | S7a vs. S8a |
| *Spiroplasma* | Firmicutes | 7.936 | 0.000 | 0.000 | S1 vs. S7a |
| *Ac37b* | Proteobacteria | 7.764 | 0.001 | 0.011 | S3 vs. S8a |
| *Spiroplasma* | Firmicutes | 7.245 | 0.000 | 0.004 | S2 vs. S8a |
| *Spiroplasma* | Firmicutes | 7.245 | 0.000 | 0.000 | S2 vs. S6 |
| *Enterococcus* | Firmicutes | 6.559 | 0.000 | 0.000 | S3 vs. S6 |
| *Enterococcus* | Firmicutes | 6.559 | 0.000 | 0.002 | S3 vs. S5 |
| *Enterococcus* | Firmicutes | 6.558 | 0.000 | 0.000 | S3 vs. S8a |
| *Spiroplasma* | Firmicutes | 6.341 | 0.000 | 0.000 | S3 vs. S4 |
| *Borrelia* | Spirochaetota | 6.276 | 0.000 | 0.000 | S5 vs. S6 |
| *Borrelia* | Spirochaetota | 6.275 | 0.000 | 0.004 | S5 vs. S8a |
| *Spiroplasma* | Firmicutes | 6.196 | 0.000 | 0.000 | S2 vs. S7a |
| *Ac37b* | Proteobacteria | 6.013 | 0.000 | 0.000 | S1 vs. S7a |
| *Ac37b* | Proteobacteria | 6.012 | 0.000 | 0.000 | S1 vs. S3 |

**Note:** Data shows genera with significant differential abundance across sites.

**Table 12.Significant Differential Abundance of Genera Across Site Comparisons**

*Filtered for Adjusted p-value < 0.05*

| Genus | Phylum | **Log2 Fold Change** | p-value | **Adjusted p-value** | Site Comparison |
|---|---|---|---|---|---|
| *Ac37b* | Proteobacteria | 6.011 | 0.000 | 0.000 | S1 vs. S6 |
| *Ac37b* | Proteobacteria | 6.011 | 0.000 | 0.000 | S1 vs. S2 |
| *Ac37b* | Proteobacteria | 6.011 | 0.000 | 0.000 | S1 vs. S4 |
| *Ac37b* | Proteobacteria | 6.011 | 0.000 | 0.003 | S1 vs. S5 |
| *Ac37b* | Proteobacteria | 6.011 | 0.000 | 0.000 | S1 vs. S8a |
| *Borrelia* | Spirochaetota | 5.969 | 0.000 | 0.000 | S1 vs. S7a |
| *Borrelia* | Spirochaetota | 5.967 | 0.000 | 0.000 | S1 vs. S3 |
| *Borrelia* | Spirochaetota | 5.967 | 0.000 | 0.000 | S1 vs. S6 |
| *Borrelia* | Spirochaetota | 5.967 | 0.000 | 0.000 | S1 vs. S2 |
| *Borrelia* | Spirochaetota | 5.967 | 0.000 | 0.000 | S1 vs. S4 |
| *Borrelia* | Spirochaetota | 5.967 | 0.000 | 0.003 | S1 vs. S5 |
| *Borrelia* | Spirochaetota | 5.966 | 0.000 | 0.000 | S1 vs. S8a |
| *Bradyrhizobium* | Proteobacteria | 5.534 | 0.003 | 0.029 | S3 vs. S5 |
| *Bradyrhizobium* | Proteobacteria | 5.534 | 0.000 | 0.008 | S3 vs. S8a |
| *Blattabacterium* | Bacteroidota | 5.487 | 0.000 | 0.000 | S3 vs. S7a |
| *Blattabacterium* | Bacteroidota | 5.486 | 0.000 | 0.000 | S3 vs. S6 |
| *Blattabacterium* | Bacteroidota | 5.485 | 0.000 | 0.000 | S3 vs. S4 |
| *Blattabacterium* | Bacteroidota | 5.485 | 0.000 | 0.002 | S3 vs. S5 |
| *Blattabacterium* | Bacteroidota | 5.485 | 0.000 | 0.000 | S3 vs. S8a |
| *Rickettsiella* | Proteobacteria | 5.375 | 0.000 | 0.000 | S5 vs. S6 |
| *Rickettsiella* | Proteobacteria | 5.374 | 0.000 | 0.005 | S5 vs. S8a |
|  | Patescibacteria | 5.336 | 0.000 | 0.000 | S3 vs. S7a |

**Note:** Data shows genera with significant differential abundance across sites.

**Table 12.Significant Differential Abundance of Genera Across Site Comparisons**

*Filtered for Adjusted p-value < 0.05*

| Genus | Phylum | **Log2 Fold Change** | p-value | **Adjusted p-value** | Site Comparison |
|---|---|---|---|---|---|
| | Patescibacteria | 5.335 | 0.000 | 0.000 | S3 vs. S6 |
| | Patescibacteria | 5.334 | 0.000 | 0.000 | S3 vs. S4 |
| | Patescibacteria | 5.334 | 0.000 | 0.005 | S3 vs. S5 |
| | Patescibacteria | 5.334 | 0.000 | 0.000 | S3 vs. S8a |
| *Enterococcus* | Firmicutes | 5.172 | 0.000 | 0.000 | S3 vs. S4 |
| *Cutibacterium* | Actinobacteriota | 5.111 | 0.000 | 0.009 | S1 vs. S2 |
| *Enterococcus* | Firmicutes | 5.009 | 0.000 | 0.000 | S3 vs. S7a |
| *Bradyrhizobium* | Proteobacteria | 4.987 | 0.000 | 0.000 | S3 vs. S6 |
| *Mycobacterium* | Actinobacteriota | 4.878 | 0.000 | 0.000 | S3 vs. S7a |
| *Mycobacterium* | Actinobacteriota | 4.876 | 0.000 | 0.000 | S3 vs. S6 |
| *Mycobacterium* | Actinobacteriota | 4.876 | 0.001 | 0.013 | S3 vs. S5 |
| *Mycobacterium* | Actinobacteriota | 4.875 | 0.000 | 0.002 | S3 vs. S8a |
| *Borrelia* | Spirochaetota | 4.875 | 0.000 | 0.002 | S5 vs. S7a |
| *Spiroplasma* | Firmicutes | 4.757 | 0.000 | 0.000 | S5 vs. S7a |
| *Spiroplasma* | Firmicutes | 4.755 | 0.000 | 0.000 | S5 vs. S6 |
| *Spiroplasma* | Firmicutes | 4.754 | 0.000 | 0.001 | S5 vs. S8a |
| *Acinetobacter* | Proteobacteria | 4.689 | 0.000 | 0.000 | S3 vs. S7a |
| *Acinetobacter* | Proteobacteria | 4.688 | 0.000 | 0.000 | S3 vs. S6 |
| *Acinetobacter* | Proteobacteria | 4.687 | 0.000 | 0.000 | S3 vs. S4 |
| *Acinetobacter* | Proteobacteria | 4.687 | 0.001 | 0.014 | S3 vs. S5 |
| *Acinetobacter* | Proteobacteria | 4.687 | 0.000 | 0.002 | S3 vs. S8a |
| *Rickettsiella* | Proteobacteria | 4.666 | 0.000 | 0.000 | S5 vs. S7a |

**Note:** Data shows genera with significant differential abundance across sites.

### Table 12.Significant Differential Abundance of Genera Across Site Comparisons

*Filtered for Adjusted p-value < 0.05*

| Genus | Phylum | **Log2 Fold Change** | p-value | **Adjusted p-value** | Site Comparison |
|---|---|---|---|---|---|
| *Mycoplasma* | Firmicutes | 4.443 | 0.000 | 0.000 | S1 vs. S7a |
| *Mycoplasma* | Firmicutes | 4.442 | 0.000 | 0.000 | S1 vs. S3 |
| *Mycoplasma* | Firmicutes | 4.441 | 0.000 | 0.000 | S1 vs. S6 |
| *Mycoplasma* | Firmicutes | 4.441 | 0.000 | 0.000 | S1 vs. S2 |
| *Mycoplasma* | Firmicutes | 4.441 | 0.000 | 0.000 | S1 vs. S4 |
| *Mycoplasma* | Firmicutes | 4.441 | 0.001 | 0.035 | S1 vs. S5 |
| *Mycoplasma* | Firmicutes | 4.440 | 0.000 | 0.005 | S1 vs. S8a |
| *Spiroplasma* | Firmicutes | 4.415 | 0.002 | 0.028 | S1 vs. S4 |
| *Blattabacterium* | Bacteroidota | 4.406 | 0.000 | 0.000 | S3 vs. S7a |
| *Blattabacterium* | Bacteroidota | 4.404 | 0.000 | 0.000 | S3 vs. S6 |
| *Blattabacterium* | Bacteroidota | 4.404 | 0.004 | 0.032 | S3 vs. S5 |
| *Blattabacterium* | Bacteroidota | 4.403 | 0.001 | 0.011 | S3 vs. S8a |
| *Dysgonomonas* | Bacteroidota | 4.304 | 0.000 | 0.000 | S1 vs. S7a |
| *Dysgonomonas* | Bacteroidota | 4.302 | 0.000 | 0.001 | S1 vs. S3 |
| *Dysgonomonas* | Bacteroidota | 4.302 | 0.000 | 0.000 | S1 vs. S6 |
| *Dysgonomonas* | Bacteroidota | 4.302 | 0.000 | 0.000 | S1 vs. S2 |
| *Dysgonomonas* | Bacteroidota | 4.302 | 0.000 | 0.000 | S1 vs. S4 |
| *Dysgonomonas* | Bacteroidota | 4.302 | 0.002 | 0.040 | S1 vs. S5 |
| *Dysgonomonas* | Bacteroidota | 4.301 | 0.000 | 0.007 | S1 vs. S8a |
| | Patescibacteria | 4.263 | 0.000 | 0.000 | S3 vs. S7a |
| | Patescibacteria | 4.261 | 0.000 | 0.000 | S3 vs. S6 |
| | Patescibacteria | 4.261 | 0.000 | 0.000 | S3 vs. S4 |

**Note:** Data shows genera with significant differential abundance across sites.

### Table 12.Significant Differential Abundance of Genera Across Site Comparisons

*Filtered for Adjusted p-value < 0.05*

| Genus | Phylum | **Log2 Fold Change** | p-value | **Adjusted p-value** | Site Comparison |
|---|---|---|---|---|---|
| | Patescibacteria | 4.261 | 0.002 | 0.027 | S3 vs. S5 |
| | Patescibacteria | 4.261 | 0.000 | 0.006 | S3 vs. S8a |
| *Bacillus* | Firmicutes | 4.202 | 0.000 | 0.000 | S3 vs. S7a |
| *Bacillus* | Firmicutes | 4.200 | 0.000 | 0.000 | S3 vs. S6 |
| *Bacillus* | Firmicutes | 4.199 | 0.001 | 0.020 | S3 vs. S8a |
| *Mycobacterium* | Actinobacteriota | 4.184 | 0.000 | 0.000 | S3 vs. S4 |
| *Jatrophihabitans* | Actinobacteriota | 4.053 | 0.000 | 0.000 | S3 vs. S7a |
| *Jatrophihabitans* | Actinobacteriota | 4.051 | 0.000 | 0.000 | S3 vs. S6 |
| *Jatrophihabitans* | Actinobacteriota | 4.051 | 0.000 | 0.000 | S3 vs. S4 |
| *Jatrophihabitans* | Actinobacteriota | 4.051 | 0.003 | 0.030 | S3 vs. S5 |
| *Jatrophihabitans* | Actinobacteriota | 4.051 | 0.001 | 0.009 | S3 vs. S8a |
| | Patescibacteria | 4.036 | 0.000 | 0.000 | S3 vs. S7a |
| | Patescibacteria | 4.034 | 0.000 | 0.000 | S3 vs. S6 |
| | Patescibacteria | 4.034 | 0.000 | 0.000 | S3 vs. S4 |
| | Patescibacteria | 4.034 | 0.003 | 0.030 | S3 vs. S5 |
| | Patescibacteria | 4.033 | 0.001 | 0.009 | S3 vs. S8a |
| | Proteobacteria | 4.000 | 0.000 | 0.000 | S1 vs. S7a |
| | Proteobacteria | 3.999 | 0.000 | 0.002 | S1 vs. S3 |
| | Proteobacteria | 3.998 | 0.000 | 0.000 | S1 vs. S6 |
| | Proteobacteria | 3.998 | 0.000 | 0.000 | S1 vs. S4 |
| | Proteobacteria | 3.998 | 0.001 | 0.015 | S1 vs. S8a |
| *Blattabacterium* | Bacteroidota | 3.935 | 0.000 | 0.000 | S3 vs. S4 |

**Note:** Data shows genera with significant differential abundance across sites.

### Table 12.Significant Differential Abundance of Genera Across Site Comparisons

*Filtered for Adjusted p-value < 0.05*

| Genus | Phylum | **Log2 Fold Change** | p-value | **Adjusted p-value** | Site Comparison |
|---|---|---|---|---|---|
| | Firmicutes | 3.907 | 0.000 | 0.000 | S1 vs. S7a |
| | Firmicutes | 3.906 | 0.000 | 0.002 | S1 vs. S3 |
| | Firmicutes | 3.905 | 0.000 | 0.000 | S1 vs. S6 |
| | Firmicutes | 3.905 | 0.000 | 0.000 | S1 vs. S2 |
| | Firmicutes | 3.905 | 0.000 | 0.000 | S1 vs. S4 |
| | Firmicutes | 3.905 | 0.001 | 0.016 | S1 vs. S8a |
| | Proteobacteria | 3.883 | 0.000 | 0.000 | S1 vs. S7a |
| | Bacteroidota | 3.883 | 0.000 | 0.000 | S1 vs. S7a |
| | Proteobacteria | 3.881 | 0.000 | 0.002 | S1 vs. S3 |
| | Bacteroidota | 3.881 | 0.000 | 0.007 | S1 vs. S3 |
| | Proteobacteria | 3.881 | 0.000 | 0.000 | S1 vs. S6 |
| | Bacteroidota | 3.881 | 0.000 | 0.000 | S1 vs. S6 |
| | Proteobacteria | 3.881 | 0.000 | 0.000 | S1 vs. S2 |
| | Proteobacteria | 3.881 | 0.000 | 0.000 | S1 vs. S4 |
| | Bacteroidota | 3.881 | 0.000 | 0.001 | S1 vs. S2 |
| | Proteobacteria | 3.880 | 0.001 | 0.016 | S1 vs. S8a |
| | Bacteroidota | 3.880 | 0.003 | 0.041 | S1 vs. S8a |
| | WPS-2 | 3.868 | 0.000 | 0.000 | S3 vs. S7a |
| | WPS-2 | 3.866 | 0.000 | 0.000 | S3 vs. S4 |
| | WPS-2 | 3.865 | 0.002 | 0.025 | S3 vs. S8a |
| *uncultured* | Actinobacteriota | 3.858 | 0.000 | 0.000 | S1 vs. S7a |
| *uncultured* | Actinobacteriota | 3.857 | 0.000 | 0.002 | S1 vs. S3 |

**Note:** Data shows genera with significant differential abundance across sites.

### Table 12.Significant Differential Abundance of Genera Across Site Comparisons

*Filtered for Adjusted p-value < 0.05*

| Genus | Phylum | **Log2 Fold Change** | p-value | **Adjusted p-value** | Site Comparison |
|---|---|---|---|---|---|
| *uncultured* | Actinobacteriota | 3.856 | 0.000 | 0.000 | S1 vs. S6 |
| *uncultured* | Actinobacteriota | 3.856 | 0.000 | 0.000 | S1 vs. S2 |
| *uncultured* | Actinobacteriota | 3.856 | 0.000 | 0.000 | S1 vs. S4 |
| *uncultured* | Actinobacteriota | 3.856 | 0.001 | 0.016 | S1 vs. S8a |
| *Rickettsiella* | Proteobacteria | 3.789 | 0.000 | 0.000 | S3 vs. S7a |
| *Rickettsiella* | Proteobacteria | 3.787 | 0.000 | 0.000 | S3 vs. S6 |
| *Rickettsiella* | Proteobacteria | 3.786 | 0.003 | 0.034 | S3 vs. S8a |
| *Blattabacterium* | Bacteroidota | 3.782 | 0.000 | 0.001 | S1 vs. S7a |
| *Blattabacterium* | Bacteroidota | 3.780 | 0.000 | 0.001 | S1 vs. S6 |
| *Blattabacterium* | Bacteroidota | 3.780 | 0.000 | 0.002 | S1 vs. S2 |
| | Proteobacteria | 3.755 | 0.000 | 0.000 | S1 vs. S7a |
| | Proteobacteria | 3.754 | 0.000 | 0.002 | S1 vs. S3 |
| *Rickettsiella* | Proteobacteria | 3.754 | 0.000 | 0.007 | S1 vs. S3 |
| | Proteobacteria | 3.753 | 0.000 | 0.000 | S1 vs. S6 |
| *Rickettsiella* | Proteobacteria | 3.753 | 0.000 | 0.000 | S1 vs. S6 |
| | Proteobacteria | 3.753 | 0.000 | 0.000 | S1 vs. S2 |
| | Proteobacteria | 3.753 | 0.000 | 0.000 | S1 vs. S4 |
| *Rickettsiella* | Proteobacteria | 3.753 | 0.000 | 0.001 | S1 vs. S2 |
| *Rickettsiella* | Proteobacteria | 3.753 | 0.000 | 0.000 | S1 vs. S4 |
| | Proteobacteria | 3.753 | 0.001 | 0.019 | S1 vs. S8a |
| *Rickettsiella* | Proteobacteria | 3.752 | 0.003 | 0.041 | S1 vs. S8a |
| *Faecalibacterium* | Firmicutes | 3.728 | 0.000 | 0.000 | S1 vs. S7a |

**Note:** Data shows genera with significant differential abundance across sites.

### Table 12.Significant Differential Abundance of Genera Across Site Comparisons

*Filtered for Adjusted p-value < 0.05*

| Genus | Phylum | **Log2 Fold Change** | p-value | **Adjusted p-value** | Site Comparison |
|---|---|---|---|---|---|
| *Faecalibacterium* | Firmicutes | 3.727 | 0.000 | 0.002 | S1 vs. S3 |
| *Faecalibacterium* | Firmicutes | 3.726 | 0.000 | 0.000 | S1 vs. S6 |
| *Faecalibacterium* | Firmicutes | 3.726 | 0.000 | 0.000 | S1 vs. S2 |
| *Faecalibacterium* | Firmicutes | 3.726 | 0.000 | 0.000 | S1 vs. S4 |
| *Faecalibacterium* | Firmicutes | 3.726 | 0.001 | 0.020 | S1 vs. S8a |
| *Brevundimonas* | Proteobacteria | 3.702 | 0.000 | 0.002 | S5 vs. S7a |
| *Brevundimonas* | Proteobacteria | 3.701 | 0.000 | 0.002 | S5 vs. S6 |
| *Mycobacterium* | Actinobacteriota | 3.674 | 0.000 | 0.000 | S1 vs. S7a |
| *Mycobacterium* | Actinobacteriota | 3.673 | 0.000 | 0.003 | S1 vs. S3 |
| *Mycobacterium* | Actinobacteriota | 3.673 | 0.000 | 0.000 | S1 vs. S6 |
| *Mycobacterium* | Actinobacteriota | 3.672 | 0.000 | 0.000 | S1 vs. S2 |
| *Mycobacterium* | Actinobacteriota | 3.672 | 0.000 | 0.000 | S1 vs. S4 |
| *Mycobacterium* | Actinobacteriota | 3.672 | 0.001 | 0.022 | S1 vs. S8a |
| *Candidatus* | Crenarchaeota | 3.646 | 0.000 | 0.000 | S5 vs. S7a |
| *Candidatus* | Crenarchaeota | 3.644 | 0.000 | 0.000 | S5 vs. S6 |
| *Candidatus* | Crenarchaeota | 3.643 | 0.001 | 0.028 | S5 vs. S8a |
| | Proteobacteria | 3.557 | 0.000 | 0.000 | S1 vs. S7a |
| | Proteobacteria | 3.555 | 0.000 | 0.003 | S1 vs. S3 |
| | Proteobacteria | 3.555 | 0.000 | 0.000 | S1 vs. S6 |
| | Proteobacteria | 3.555 | 0.000 | 0.000 | S1 vs. S2 |
| | Proteobacteria | 3.555 | 0.000 | 0.000 | S1 vs. S4 |
| | Proteobacteria | 3.554 | 0.001 | 0.026 | S1 vs. S8a |

**Note:** Data shows genera with significant differential abundance across sites.

### Table 12.Significant Differential Abundance of Genera Across Site Comparisons

*Filtered for Adjusted p-value < 0.05*

| Genus | Phylum | **Log2 Fold Change** | p-value | **Adjusted p-value** | Site Comparison |
|---|---|---|---|---|---|
| | Proteobacteria | 3.513 | 0.000 | 0.000 | S1 vs. S2 |
| *Phenylobacterium* | Proteobacteria | 3.461 | 0.000 | 0.000 | S5 vs. S7a |
| *Tyzzerella* | Firmicutes | 3.460 | 0.000 | 0.000 | S1 vs. S7a |
| *Lysobacter* | Proteobacteria | 3.460 | 0.000 | 0.000 | S1 vs. S7a |
| *Roseiarcus* | Proteobacteria | 3.460 | 0.000 | 0.000 | S3 vs. S7a |
| *Phenylobacterium* | Proteobacteria | 3.460 | 0.000 | 0.000 | S5 vs. S6 |
| *Phenylobacterium* | Proteobacteria | 3.459 | 0.001 | 0.043 | S5 vs. S8a |
| *Tyzzerella* | Firmicutes | 3.458 | 0.000 | 0.004 | S1 vs. S3 |
| *Lysobacter* | Proteobacteria | 3.458 | 0.000 | 0.004 | S1 vs. S3 |
| *Tyzzerella* | Firmicutes | 3.458 | 0.000 | 0.000 | S1 vs. S6 |
| *Lysobacter* | Proteobacteria | 3.458 | 0.000 | 0.000 | S1 vs. S6 |
| *Roseiarcus* | Proteobacteria | 3.458 | 0.000 | 0.000 | S3 vs. S6 |
| *Tyzzerella* | Firmicutes | 3.458 | 0.000 | 0.000 | S1 vs. S2 |
| *Lysobacter* | Proteobacteria | 3.458 | 0.000 | 0.000 | S1 vs. S2 |
| *Tyzzerella* | Firmicutes | 3.458 | 0.000 | 0.000 | S1 vs. S4 |
| *Lysobacter* | Proteobacteria | 3.458 | 0.000 | 0.000 | S1 vs. S4 |
| *Roseiarcus* | Proteobacteria | 3.458 | 0.000 | 0.000 | S3 vs. S4 |
| *Tyzzerella* | Firmicutes | 3.457 | 0.002 | 0.031 | S1 vs. S8a |
| *Lysobacter* | Proteobacteria | 3.457 | 0.002 | 0.031 | S1 vs. S8a |
| *Roseiarcus* | Proteobacteria | 3.457 | 0.002 | 0.027 | S3 vs. S8a |
| *Enhydrobacter* | Proteobacteria | 3.407 | 0.000 | 0.000 | S3 vs. S7a |
| *Enhydrobacter* | Proteobacteria | 3.405 | 0.000 | 0.000 | S3 vs. S6 |

**Note:** Data shows genera with significant differential abundance across sites.

### Table 12. Significant Differential Abundance of Genera Across Site Comparisons

*Filtered for Adjusted p-value < 0.05*

| Genus | Phylum | **Log2 Fold Change** | p-value | **Adjusted p-value** | Site Comparison |
|---|---|---|---|---|---|
| *Enhydrobacter* | Proteobacteria | 3.405 | 0.000 | 0.000 | S3 vs. S4 |
| *Enhydrobacter* | Proteobacteria | 3.404 | 0.003 | 0.032 | S3 vs. S8a |
| *Escherichia-Shigella* | Proteobacteria | 3.394 | 0.000 | 0.000 | S5 vs. S7a |
| | Bacteroidota | 3.393 | 0.000 | 0.000 | S1 vs. S7a |
| *Escherichia-Shigella* | Proteobacteria | 3.393 | 0.000 | 0.000 | S5 vs. S6 |
| *Escherichia-Shigella* | Proteobacteria | 3.392 | 0.001 | 0.048 | S5 vs. S8a |
| | Bacteroidota | 3.391 | 0.000 | 0.005 | S1 vs. S3 |
| | Bacteroidota | 3.391 | 0.000 | 0.000 | S1 vs. S6 |
| | Bacteroidota | 3.391 | 0.000 | 0.000 | S1 vs. S2 |
| | Bacteroidota | 3.391 | 0.000 | 0.000 | S1 vs. S4 |
| | Bacteroidota | 3.390 | 0.002 | 0.035 | S1 vs. S8a |
| | Actinobacteriota | 3.322 | 0.000 | 0.000 | S1 vs. S7a |
| | Actinobacteriota | 3.321 | 0.000 | 0.006 | S1 vs. S3 |
| | Actinobacteriota | 3.321 | 0.000 | 0.000 | S1 vs. S6 |
| | Actinobacteriota | 3.320 | 0.000 | 0.000 | S1 vs. S2 |
| | Actinobacteriota | 3.320 | 0.000 | 0.000 | S1 vs. S4 |
| | Actinobacteriota | 3.320 | 0.002 | 0.039 | S1 vs. S8a |
| *Blattabacterium* | Bacteroidota | 3.310 | 0.000 | 0.003 | S1 vs. S4 |
| *Sphingomonas* | Proteobacteria | 3.265 | 0.000 | 0.005 | S3 vs. S7a |
| *Sphingomonas* | Proteobacteria | 3.263 | 0.000 | 0.004 | S3 vs. S4 |
| | Verrucomicrobiota | 3.248 | 0.000 | 0.000 | S1 vs. S7a |
| | Verrucomicrobiota | 3.247 | 0.000 | 0.007 | S1 vs. S3 |

**Note:** Data shows genera with significant differential abundance across sites.

### Table 12.Significant Differential Abundance of Genera Across Site Comparisons

*Filtered for Adjusted p-value < 0.05*

| Genus | Phylum | **Log2 Fold Change** | p-value | **Adjusted p-value** | Site Comparison |
|---|---|---|---|---|---|
| | Verrucomicrobiota | 3.247 | 0.000 | 0.000 | S1 vs. S6 |
| | Verrucomicrobiota | 3.246 | 0.000 | 0.001 | S1 vs. S2 |
| | Verrucomicrobiota | 3.246 | 0.000 | 0.000 | S1 vs. S4 |
| | Verrucomicrobiota | 3.246 | 0.003 | 0.042 | S1 vs. S8a |
| *Bradyrhizobium* | Proteobacteria | 3.212 | 0.001 | 0.017 | S3 vs. S4 |
| *Candidatus* | Crenarchaeota | 3.210 | 0.000 | 0.000 | S1 vs. S7a |
| *Candidatus* | Crenarchaeota | 3.208 | 0.000 | 0.007 | S1 vs. S3 |
| *Candidatus* | Crenarchaeota | 3.208 | 0.000 | 0.000 | S1 vs. S6 |
| *Candidatus* | Crenarchaeota | 3.208 | 0.000 | 0.001 | S1 vs. S2 |
| *Candidatus* | Crenarchaeota | 3.208 | 0.000 | 0.000 | S1 vs. S4 |
| *Candidatus* | Crenarchaeota | 3.207 | 0.003 | 0.044 | S1 vs. S8a |
| *Kitasatospora* | Actinobacteriota | 3.172 | 0.000 | 0.000 | S5 vs. S7a |
| | Firmicutes | 3.170 | 0.000 | 0.000 | S1 vs. S7a |
| *Kitasatospora* | Actinobacteriota | 3.170 | 0.000 | 0.000 | S5 vs. S6 |
| | Firmicutes | 3.169 | 0.000 | 0.008 | S1 vs. S3 |
| | Firmicutes | 3.169 | 0.000 | 0.000 | S1 vs. S6 |
| | Firmicutes | 3.168 | 0.000 | 0.001 | S1 vs. S2 |
| | Firmicutes | 3.168 | 0.000 | 0.000 | S1 vs. S4 |
| | Firmicutes | 3.168 | 0.003 | 0.047 | S1 vs. S8a |
| *Mycobacterium* | Actinobacteriota | 3.139 | 0.000 | 0.000 | S3 vs. S7a |
| *Mycobacterium* | Actinobacteriota | 3.137 | 0.000 | 0.000 | S3 vs. S6 |
| *Mycobacterium* | Actinobacteriota | 3.137 | 0.000 | 0.000 | S3 vs. S4 |

**Note:** Data shows genera with significant differential abundance across sites.

### Table 12.Significant Differential Abundance of Genera Across Site Comparisons

*Filtered for Adjusted p-value < 0.05*

| Genus | Phylum | **Log2 Fold Change** | p-value | **Adjusted p-value** | Site Comparison |
|---|---|---|---|---|---|
| *Mycobacterium* | Actinobacteriota | 3.136 | 0.003 | 0.034 | S3 vs. S8a |
| | | 3.105 | 0.000 | 0.000 | S3 vs. S7a |
| | Acidobacteriota | 3.105 | 0.000 | 0.000 | S3 vs. S7a |
| | | 3.103 | 0.000 | 0.000 | S3 vs. S6 |
| | Acidobacteriota | 3.103 | 0.000 | 0.000 | S3 vs. S6 |
| | | 3.103 | 0.000 | 0.000 | S3 vs. S4 |
| | Acidobacteriota | 3.103 | 0.000 | 0.000 | S3 vs. S4 |
| *Enterococcus* | Firmicutes | 3.089 | 0.000 | 0.000 | S5 vs. S7a |
| *Burkholderia-Caballeronia-Paraburkholderia* | Proteobacteria | 3.088 | 0.000 | 0.000 | S1 vs. S7a |
| *uncultured* | Bacteroidota | 3.088 | 0.000 | 0.000 | S1 vs. S7a |
| *Enterococcus* | Firmicutes | 3.088 | 0.000 | 0.000 | S5 vs. S6 |
| *Burkholderia-Caballeronia-Paraburkholderia* | Proteobacteria | 3.086 | 0.000 | 0.009 | S1 vs. S3 |
| *uncultured* | Bacteroidota | 3.086 | 0.000 | 0.009 | S1 vs. S3 |
| *Burkholderia-Caballeronia-Paraburkholderia* | Proteobacteria | 3.086 | 0.000 | 0.000 | S1 vs. S6 |
| *uncultured* | Bacteroidota | 3.086 | 0.000 | 0.000 | S1 vs. S6 |
| *Burkholderia-Caballeronia-Paraburkholderia* | Proteobacteria | 3.086 | 0.000 | 0.001 | S1 vs. S2 |
| *uncultured* | Bacteroidota | 3.086 | 0.000 | 0.001 | S1 vs. S2 |

**Note:** Data shows genera with significant differential abundance across sites.

**Table 12. Significant Differential Abundance of Genera Across Site Comparisons**

*Filtered for Adjusted p-value < 0.05*

| Genus | Phylum | **Log2 Fold Change** | p-value | **Adjusted p-value** | Site Comparison |
|---|---|---|---|---|---|
| *Burkholderia-Caballeronia-Paraburkholderia* | Proteobacteria | 3.086 | 0.000 | 0.000 | S1 vs. S4 |
| *uncultured* | Bacteroidota | 3.086 | 0.000 | 0.000 | S1 vs. S4 |
| *Chthoniobacter* | Verrucomicrobiota | 3.071 | 0.000 | 0.000 | S3 vs. S7a |
| *Chthoniobacter* | Verrucomicrobiota | 3.069 | 0.000 | 0.000 | S3 vs. S6 |
| *Chthoniobacter* | Verrucomicrobiota | 3.069 | 0.000 | 0.000 | S3 vs. S4 |
| *Anaeromyxobacter* | Myxococcota | 3.045 | 0.000 | 0.000 | S1 vs. S7a |
| *H1* | Bacteroidota | 3.045 | 0.000 | 0.000 | S1 vs. S7a |
| *Dendrosporobacter* | Firmicutes | 3.045 | 0.000 | 0.000 | S1 vs. S7a |
| *Atopobium* | Actinobacteriota | 3.045 | 0.000 | 0.000 | S1 vs. S7a |
| *Rickettsiella* | Proteobacteria | 3.044 | 0.000 | 0.002 | S1 vs. S7a |
| *Anaeromyxobacter* | Myxococcota | 3.043 | 0.001 | 0.009 | S1 vs. S3 |
| *H1* | Bacteroidota | 3.043 | 0.001 | 0.009 | S1 vs. S3 |
| *Dendrosporobacter* | Firmicutes | 3.043 | 0.001 | 0.009 | S1 vs. S3 |
| *Atopobium* | Actinobacteriota | 3.043 | 0.001 | 0.009 | S1 vs. S3 |
| *Anaeromyxobacter* | Myxococcota | 3.043 | 0.000 | 0.000 | S1 vs. S6 |
| *H1* | Bacteroidota | 3.043 | 0.000 | 0.000 | S1 vs. S6 |
| *Dendrosporobacter* | Firmicutes | 3.043 | 0.000 | 0.000 | S1 vs. S6 |
| *Atopobium* | Actinobacteriota | 3.043 | 0.000 | 0.000 | S1 vs. S6 |
| *Anaeromyxobacter* | Myxococcota | 3.043 | 0.000 | 0.001 | S1 vs. S2 |
| *H1* | Bacteroidota | 3.043 | 0.000 | 0.001 | S1 vs. S2 |
| *Dendrosporobacter* | Firmicutes | 3.043 | 0.000 | 0.001 | S1 vs. S2 |

**Note:** Data shows genera with significant differential abundance across sites.

### Table 12.Significant Differential Abundance of Genera Across Site Comparisons

*Filtered for Adjusted p-value < 0.05*

| Genus | Phylum | **Log2 Fold Change** | p-value | **Adjusted p-value** | Site Comparison |
|---|---|---|---|---|---|
| *Atopobium* | Actinobacteriota | 3.043 | 0.000 | 0.001 | S1 vs. S2 |
| *Anaeromyxobacter* | Myxococcota | 3.043 | 0.000 | 0.000 | S1 vs. S4 |
| *H1* | Bacteroidota | 3.043 | 0.000 | 0.000 | S1 vs. S4 |
| *Dendrosporobacter* | Firmicutes | 3.043 | 0.000 | 0.000 | S1 vs. S4 |
| *Atopobium* | Actinobacteriota | 3.043 | 0.000 | 0.000 | S1 vs. S4 |
| | Firmicutes | 3.001 | 0.000 | 0.000 | S1 vs. S7a |
| *Acidisphaera* | Proteobacteria | 3.001 | 0.000 | 0.000 | S3 vs. S7a |
| | Firmicutes | 2.999 | 0.001 | 0.010 | S1 vs. S3 |
| | Firmicutes | 2.999 | 0.000 | 0.000 | S1 vs. S6 |
| *Acidisphaera* | Proteobacteria | 2.999 | 0.000 | 0.000 | S3 vs. S6 |
| | Firmicutes | 2.999 | 0.000 | 0.001 | S1 vs. S2 |
| | Firmicutes | 2.999 | 0.000 | 0.000 | S1 vs. S4 |
| *Acidisphaera* | Proteobacteria | 2.999 | 0.000 | 0.000 | S3 vs. S4 |
| *NK4A214* | Firmicutes | 2.955 | 0.000 | 0.000 | S1 vs. S7a |
| *Candidatus* | Bacteroidota | 2.955 | 0.000 | 0.000 | S1 vs. S7a |
| *uncultured* | Verrucomicrobiota | 2.955 | 0.000 | 0.000 | S1 vs. S7a |
| *NK4A214* | Firmicutes | 2.953 | 0.001 | 0.011 | S1 vs. S3 |
| *Candidatus* | Bacteroidota | 2.953 | 0.001 | 0.011 | S1 vs. S3 |
| *uncultured* | Verrucomicrobiota | 2.953 | 0.001 | 0.011 | S1 vs. S3 |
| *NK4A214* | Firmicutes | 2.953 | 0.000 | 0.000 | S1 vs. S6 |
| *Candidatus* | Bacteroidota | 2.953 | 0.000 | 0.000 | S1 vs. S6 |
| *uncultured* | Verrucomicrobiota | 2.953 | 0.000 | 0.000 | S1 vs. S6 |

**Note:** Data shows genera with significant differential abundance across sites.

### Table 12.Significant Differential Abundance of Genera Across Site Comparisons

*Filtered for Adjusted p-value < 0.05*

| Genus | Phylum | **Log2 Fold Change** | p-value | **Adjusted p-value** | Site Comparison |
|---|---|---|---|---|---|
| *NK4A214* | Firmicutes | 2.953 | 0.000 | 0.001 | S1 vs. S2 |
| *Candidatus* | Bacteroidota | 2.953 | 0.000 | 0.001 | S1 vs. S2 |
| *uncultured* | Verrucomicrobiota | 2.953 | 0.000 | 0.001 | S1 vs. S2 |
| *NK4A214* | Firmicutes | 2.953 | 0.000 | 0.000 | S1 vs. S4 |
| *Candidatus* | Bacteroidota | 2.953 | 0.000 | 0.000 | S1 vs. S4 |
| *uncultured* | Verrucomicrobiota | 2.953 | 0.000 | 0.000 | S1 vs. S4 |
| | Proteobacteria | 2.953 | 0.001 | 0.018 | S1 vs. S2 |
| | Proteobacteria | 2.953 | 0.000 | 0.009 | S1 vs. S4 |
| *Rhodovastum* | Proteobacteria | 2.928 | 0.000 | 0.000 | S3 vs. S7a |
| *Micrococcus* | Actinobacteriota | 2.928 | 0.000 | 0.000 | S3 vs. S7a |
| *Micrococcus* | Actinobacteriota | 2.926 | 0.000 | 0.000 | S3 vs. S6 |
| *Rhodovastum* | Proteobacteria | 2.926 | 0.000 | 0.000 | S3 vs. S6 |
| *Rhodovastum* | Proteobacteria | 2.926 | 0.000 | 0.000 | S3 vs. S4 |
| *Micrococcus* | Actinobacteriota | 2.926 | 0.000 | 0.000 | S3 vs. S4 |
| *Spiroplasma* | Firmicutes | 2.909 | 0.000 | 0.001 | S5 vs. S7a |
| | Proteobacteria | 2.908 | 0.000 | 0.000 | S1 vs. S7a |
| *Rikenellaceae* | Bacteroidota | 2.908 | 0.000 | 0.000 | S1 vs. S7a |
| | Acidobacteriota | 2.908 | 0.000 | 0.000 | S1 vs. S7a |
| *Spiroplasma* | Firmicutes | 2.907 | 0.000 | 0.001 | S5 vs. S6 |
| | Proteobacteria | 2.906 | 0.001 | 0.011 | S1 vs. S3 |
| *Rikenellaceae* | Bacteroidota | 2.906 | 0.001 | 0.011 | S1 vs. S3 |
| | Acidobacteriota | 2.906 | 0.001 | 0.011 | S1 vs. S3 |

**Note:** Data shows genera with significant differential abundance across sites.

### Table 12.Significant Differential Abundance of Genera Across Site Comparisons

*Filtered for Adjusted p-value < 0.05*

| Genus | Phylum | **Log2 Fold Change** | p-value | **Adjusted p-value** | Site Comparison |
|---|---|---|---|---|---|
| | Proteobacteria | 2.906 | 0.000 | 0.000 | S1 vs. S6 |
| *Rikenellaceae* | Bacteroidota | 2.906 | 0.000 | 0.000 | S1 vs. S6 |
| | Acidobacteriota | 2.906 | 0.000 | 0.000 | S1 vs. S6 |
| | Proteobacteria | 2.906 | 0.000 | 0.001 | S1 vs. S2 |
| *Rikenellaceae* | Bacteroidota | 2.906 | 0.000 | 0.001 | S1 vs. S2 |
| | Acidobacteriota | 2.906 | 0.000 | 0.001 | S1 vs. S2 |
| | Proteobacteria | 2.906 | 0.000 | 0.000 | S1 vs. S4 |
| *Rikenellaceae* | Bacteroidota | 2.906 | 0.000 | 0.000 | S1 vs. S4 |
| | Acidobacteriota | 2.906 | 0.000 | 0.000 | S1 vs. S4 |
| | Bacteroidota | 2.859 | 0.000 | 0.000 | S1 vs. S7a |
| | Bacteroidota | 2.857 | 0.001 | 0.013 | S1 vs. S3 |
| | Bacteroidota | 2.857 | 0.000 | 0.000 | S1 vs. S6 |
| | Bacteroidota | 2.857 | 0.000 | 0.002 | S1 vs. S2 |
| | Bacteroidota | 2.857 | 0.000 | 0.000 | S1 vs. S4 |
| *uncultured* | Bacteroidota | 2.808 | 0.000 | 0.000 | S1 vs. S7a |
| *uncultured* | Bacteroidota | 2.806 | 0.001 | 0.015 | S1 vs. S3 |
| *uncultured* | Bacteroidota | 2.806 | 0.000 | 0.001 | S1 vs. S6 |
| *uncultured* | Bacteroidota | 2.806 | 0.000 | 0.002 | S1 vs. S2 |
| *uncultured* | Bacteroidota | 2.806 | 0.000 | 0.001 | S1 vs. S4 |
| *Conexibacter* | Actinobacteriota | 2.766 | 0.000 | 0.001 | S3 vs. S7a |
| | Dependentiae | 2.766 | 0.000 | 0.001 | S3 vs. S7a |
| | Planctomycetota | 2.766 | 0.000 | 0.001 | S3 vs. S7a |

**Note:** Data shows genera with significant differential abundance across sites.

### Table 12.Significant Differential Abundance of Genera Across Site Comparisons

*Filtered for Adjusted p-value < 0.05*

| Genus | Phylum | **Log2 Fold Change** | p-value | **Adjusted p-value** | Site Comparison |
|---|---|---|---|---|---|
| *Conexibacter* | Actinobacteriota | 2.764 | 0.000 | 0.001 | S3 vs. S6 |
| | Dependentiae | 2.764 | 0.000 | 0.001 | S3 vs. S6 |
| | Planctomycetota | 2.764 | 0.000 | 0.001 | S3 vs. S6 |
| *Conexibacter* | Actinobacteriota | 2.764 | 0.000 | 0.001 | S3 vs. S4 |
| | Dependentiae | 2.764 | 0.000 | 0.001 | S3 vs. S4 |
| | Planctomycetota | 2.764 | 0.000 | 0.001 | S3 vs. S4 |
| *Papillibacter* | Firmicutes | 2.756 | 0.000 | 0.000 | S1 vs. S7a |
| *Papillibacter* | Firmicutes | 2.754 | 0.001 | 0.017 | S1 vs. S3 |
| *Papillibacter* | Firmicutes | 2.754 | 0.000 | 0.001 | S1 vs. S6 |
| *Papillibacter* | Firmicutes | 2.754 | 0.000 | 0.002 | S1 vs. S2 |
| *Papillibacter* | Firmicutes | 2.754 | 0.000 | 0.001 | S1 vs. S4 |
| *Brevundimonas* | Proteobacteria | 2.724 | 0.000 | 0.002 | S3 vs. S7a |
| *uncultured* | Verrucomicrobiota | 2.723 | 0.000 | 0.001 | S3 vs. S7a |
| *Brevundimonas* | Proteobacteria | 2.722 | 0.000 | 0.002 | S3 vs. S6 |
| *Brevundimonas* | Proteobacteria | 2.722 | 0.000 | 0.002 | S3 vs. S4 |
| *uncultured* | Verrucomicrobiota | 2.721 | 0.000 | 0.001 | S3 vs. S6 |
| *uncultured* | Verrucomicrobiota | 2.721 | 0.000 | 0.001 | S3 vs. S4 |
| *Staphylococcus* | Firmicutes | 2.702 | 0.000 | 0.004 | S5 vs. S7a |
| *Pseudomonas* | Proteobacteria | 2.702 | 0.000 | 0.004 | S5 vs. S7a |
| *Staphylococcus* | Firmicutes | 2.701 | 0.000 | 0.004 | S5 vs. S6 |
| *Pseudomonas* | Proteobacteria | 2.701 | 0.000 | 0.004 | S5 vs. S6 |
| *uncultured* | Proteobacteria | 2.679 | 0.000 | 0.001 | S3 vs. S7a |

**Note:** Data shows genera with significant differential abundance across sites.

### Table 12.Significant Differential Abundance of Genera Across Site Comparisons

*Filtered for Adjusted p-value < 0.05*

| Genus | Phylum | **Log2 Fold Change** | p-value | **Adjusted p-value** | Site Comparison |
|---|---|---|---|---|---|
| *Pseudomonas* | Proteobacteria | 2.679 | 0.000 | 0.001 | S3 vs. S7a |
| *uncultured* | Proteobacteria | 2.677 | 0.000 | 0.001 | S3 vs. S6 |
| *Pseudomonas* | Proteobacteria | 2.677 | 0.000 | 0.001 | S3 vs. S6 |
| *uncultured* | Proteobacteria | 2.677 | 0.000 | 0.001 | S3 vs. S4 |
| *Pseudomonas* | Proteobacteria | 2.677 | 0.000 | 0.001 | S3 vs. S4 |
| *Sphingomonas* | Proteobacteria | 2.677 | 0.000 | 0.015 | S6 vs. S7a |
| *Corynebacterium* | Actinobacteriota | 2.645 | 0.000 | 0.001 | S1 vs. S7a |
| *Christensenellaceae* | Firmicutes | 2.645 | 0.000 | 0.001 | S1 vs. S7a |
| *Anaerovorax* | Firmicutes | 2.645 | 0.000 | 0.001 | S1 vs. S7a |
| | Proteobacteria | 2.645 | 0.000 | 0.001 | S1 vs. S7a |
| *Corynebacterium* | Actinobacteriota | 2.643 | 0.002 | 0.021 | S1 vs. S3 |
| *Christensenellaceae* | Firmicutes | 2.643 | 0.002 | 0.021 | S1 vs. S3 |
| *Anaerovorax* | Firmicutes | 2.643 | 0.002 | 0.021 | S1 vs. S3 |
| | Proteobacteria | 2.643 | 0.002 | 0.021 | S1 vs. S3 |
| *Corynebacterium* | Actinobacteriota | 2.643 | 0.000 | 0.001 | S1 vs. S6 |
| *Christensenellaceae* | Firmicutes | 2.643 | 0.000 | 0.001 | S1 vs. S6 |
| *Anaerovorax* | Firmicutes | 2.643 | 0.000 | 0.001 | S1 vs. S6 |
| | Proteobacteria | 2.643 | 0.000 | 0.001 | S1 vs. S6 |
| *Corynebacterium* | Actinobacteriota | 2.643 | 0.000 | 0.003 | S1 vs. S2 |
| *Christensenellaceae* | Firmicutes | 2.643 | 0.000 | 0.003 | S1 vs. S2 |
| *Anaerovorax* | Firmicutes | 2.643 | 0.000 | 0.003 | S1 vs. S2 |
| | Proteobacteria | 2.643 | 0.000 | 0.003 | S1 vs. S2 |

**Note:** Data shows genera with significant differential abundance across sites.

### Table 12.Significant Differential Abundance of Genera Across Site Comparisons

*Filtered for Adjusted p-value < 0.05*

| Genus | Phylum | **Log2 Fold Change** | p-value | **Adjusted p-value** | Site Comparison |
|---|---|---|---|---|---|
| *Corynebacterium* | Actinobacteriota | 2.643 | 0.000 | 0.001 | S1 vs. S4 |
| *Christensenellaceae* | Firmicutes | 2.643 | 0.000 | 0.001 | S1 vs. S4 |
| *Anaerovorax* | Firmicutes | 2.643 | 0.000 | 0.001 | S1 vs. S4 |
| | Proteobacteria | 2.643 | 0.000 | 0.001 | S1 vs. S4 |
| | Actinobacteriota | 2.633 | 0.000 | 0.001 | S3 vs. S7a |
| | Actinobacteriota | 2.631 | 0.000 | 0.001 | S3 vs. S6 |
| | Actinobacteriota | 2.631 | 0.000 | 0.001 | S3 vs. S4 |
| *uncultured* | Firmicutes | 2.586 | 0.000 | 0.001 | S1 vs. S7a |
| *Christensenellaceae* | Firmicutes | 2.586 | 0.000 | 0.001 | S1 vs. S7a |
| *Pseudomonas* | Proteobacteria | 2.586 | 0.000 | 0.001 | S1 vs. S7a |
| *uncultured* | Firmicutes | 2.584 | 0.002 | 0.023 | S1 vs. S3 |
| *Christensenellaceae* | Firmicutes | 2.584 | 0.002 | 0.023 | S1 vs. S3 |
| *Pseudomonas* | Proteobacteria | 2.584 | 0.002 | 0.023 | S1 vs. S3 |
| *uncultured* | Firmicutes | 2.584 | 0.000 | 0.001 | S1 vs. S6 |
| *Christensenellaceae* | Firmicutes | 2.584 | 0.000 | 0.001 | S1 vs. S6 |
| *Pseudomonas* | Proteobacteria | 2.584 | 0.000 | 0.001 | S1 vs. S6 |
| *uncultured* | Firmicutes | 2.584 | 0.000 | 0.004 | S1 vs. S2 |
| *Christensenellaceae* | Firmicutes | 2.584 | 0.000 | 0.004 | S1 vs. S2 |
| *Pseudomonas* | Proteobacteria | 2.584 | 0.000 | 0.004 | S1 vs. S2 |
| *uncultured* | Firmicutes | 2.584 | 0.000 | 0.001 | S1 vs. S4 |
| *Christensenellaceae* | Firmicutes | 2.584 | 0.000 | 0.001 | S1 vs. S4 |
| *Pseudomonas* | Proteobacteria | 2.584 | 0.000 | 0.001 | S1 vs. S4 |

**Note:** Data shows genera with significant differential abundance across sites.

**Table 12.Significant Differential Abundance of Genera Across Site Comparisons**

*Filtered for Adjusted p-value < 0.05*

| Genus | Phylum | **Log2 Fold Change** | p-value | **Adjusted p-value** | Site Comparison |
|---|---|---|---|---|---|
| | Actinobacteriota | 2.584 | 0.001 | 0.010 | S3 vs. S6 |
| | Actinobacteriota | 2.583 | 0.001 | 0.010 | S3 vs. S4 |
| | WPS-2 | 2.568 | 0.000 | 0.006 | S3 vs. S6 |
| *Spiroplasma* | Firmicutes | 2.538 | 0.000 | 0.002 | S3 vs. S7a |
| *Spiroplasma* | Firmicutes | 2.536 | 0.000 | 0.001 | S3 vs. S6 |
| *Spiroplasma* | Firmicutes | 2.536 | 0.000 | 0.001 | S3 vs. S4 |
| *Mycobacterium* | Actinobacteriota | 2.535 | 0.003 | 0.033 | S3 vs. S6 |
| *Uliginosibacterium* | Proteobacteria | 2.524 | 0.000 | 0.001 | S1 vs. S7a |
| *Nocardioides* | Actinobacteriota | 2.524 | 0.000 | 0.001 | S1 vs. S7a |
| | Proteobacteria | 2.524 | 0.000 | 0.001 | S1 vs. S7a |
| *Uliginosibacterium* | Proteobacteria | 2.523 | 0.003 | 0.026 | S1 vs. S3 |
| *Nocardioides* | Actinobacteriota | 2.523 | 0.003 | 0.026 | S1 vs. S3 |
| | Proteobacteria | 2.523 | 0.003 | 0.026 | S1 vs. S3 |
| *Uliginosibacterium* | Proteobacteria | 2.523 | 0.000 | 0.002 | S1 vs. S6 |
| *Nocardioides* | Actinobacteriota | 2.523 | 0.000 | 0.002 | S1 vs. S6 |
| | Proteobacteria | 2.523 | 0.000 | 0.002 | S1 vs. S6 |
| *Uliginosibacterium* | Proteobacteria | 2.522 | 0.000 | 0.005 | S1 vs. S2 |
| *Nocardioides* | Actinobacteriota | 2.522 | 0.000 | 0.005 | S1 vs. S2 |
| | Proteobacteria | 2.522 | 0.000 | 0.005 | S1 vs. S2 |
| *Uliginosibacterium* | Proteobacteria | 2.522 | 0.000 | 0.002 | S1 vs. S4 |
| *Nocardioides* | Actinobacteriota | 2.522 | 0.000 | 0.002 | S1 vs. S4 |
| | Proteobacteria | 2.522 | 0.000 | 0.002 | S1 vs. S4 |

**Note:** Data shows genera with significant differential abundance across sites.

### Table 12.Significant Differential Abundance of Genera Across Site Comparisons

*Filtered for Adjusted p-value < 0.05*

| Genus | Phylum | **Log2 Fold Change** | p-value | **Adjusted p-value** | Site Comparison |
|---|---|---|---|---|---|
| *Cutibacterium* | Actinobacteriota | 2.461 | 0.000 | 0.002 | S1 vs. S7a |
| | Desulfobacterota | 2.460 | 0.000 | 0.002 | S1 vs. S7a |
| *Cutibacterium* | Actinobacteriota | 2.460 | 0.003 | 0.031 | S1 vs. S3 |
| *Cutibacterium* | Actinobacteriota | 2.459 | 0.000 | 0.002 | S1 vs. S6 |
| *Cutibacterium* | Actinobacteriota | 2.459 | 0.000 | 0.007 | S1 vs. S2 |
| *Cutibacterium* | Actinobacteriota | 2.459 | 0.000 | 0.003 | S1 vs. S4 |
| | Desulfobacterota | 2.459 | 0.003 | 0.031 | S1 vs. S3 |
| | Desulfobacterota | 2.459 | 0.000 | 0.002 | S1 vs. S6 |
| | Desulfobacterota | 2.458 | 0.000 | 0.007 | S1 vs. S2 |
| | Desulfobacterota | 2.458 | 0.000 | 0.003 | S1 vs. S4 |
| | Acidobacteriota | 2.393 | 0.000 | 0.002 | S1 vs. S7a |
| | Firmicutes | 2.393 | 0.000 | 0.002 | S1 vs. S7a |
| *Tyzzerella* | Firmicutes | 2.393 | 0.000 | 0.002 | S1 vs. S7a |
| | Acidobacteriota | 2.392 | 0.004 | 0.035 | S1 vs. S3 |
| | Firmicutes | 2.392 | 0.004 | 0.035 | S1 vs. S3 |
| *Tyzzerella* | Firmicutes | 2.392 | 0.004 | 0.035 | S1 vs. S3 |
| | Acidobacteriota | 2.391 | 0.000 | 0.003 | S1 vs. S6 |
| | Firmicutes | 2.391 | 0.000 | 0.003 | S1 vs. S6 |
| *Tyzzerella* | Firmicutes | 2.391 | 0.000 | 0.003 | S1 vs. S6 |
| | Acidobacteriota | 2.391 | 0.000 | 0.009 | S1 vs. S2 |
| | Firmicutes | 2.391 | 0.000 | 0.009 | S1 vs. S2 |
| *Tyzzerella* | Firmicutes | 2.391 | 0.000 | 0.009 | S1 vs. S2 |

**Note:** Data shows genera with significant differential abundance across sites.

### Table 12.Significant Differential Abundance of Genera Across Site Comparisons

*Filtered for Adjusted p-value < 0.05*

| Genus | Phylum | **Log2 Fold Change** | p-value | **Adjusted p-value** | Site Comparison |
|---|---|---|---|---|---|
| | Acidobacteriota | 2.391 | 0.000 | 0.003 | S1 vs. S4 |
| | Firmicutes | 2.391 | 0.000 | 0.003 | S1 vs. S4 |
| *Tyzzerella* | Firmicutes | 2.391 | 0.000 | 0.003 | S1 vs. S4 |
| | Actinobacteriota | 2.324 | 0.001 | 0.045 | S5 vs. S7a |
| | Acidobacteriota | 2.324 | 0.000 | 0.005 | S3 vs. S7a |
| *Monoglobus* | Firmicutes | 2.323 | 0.000 | 0.004 | S1 vs. S7a |
| *Burkholderia-Caballeronia-Paraburkholderia* | Proteobacteria | 2.323 | 0.000 | 0.005 | S3 vs. S7a |
| *uncultured* | Proteobacteria | 2.323 | 0.000 | 0.005 | S3 vs. S7a |
| | Actinobacteriota | 2.322 | 0.001 | 0.047 | S5 vs. S6 |
| | Acidobacteriota | 2.322 | 0.000 | 0.004 | S3 vs. S6 |
| | Acidobacteriota | 2.322 | 0.000 | 0.004 | S3 vs. S4 |
| *Monoglobus* | Firmicutes | 2.321 | 0.005 | 0.042 | S1 vs. S3 |
| *Monoglobus* | Firmicutes | 2.321 | 0.000 | 0.005 | S1 vs. S6 |
| *Monoglobus* | Firmicutes | 2.321 | 0.001 | 0.012 | S1 vs. S2 |
| *Burkholderia-Caballeronia-Paraburkholderia* | Proteobacteria | 2.321 | 0.000 | 0.004 | S3 vs. S6 |
| *uncultured* | Proteobacteria | 2.321 | 0.000 | 0.004 | S3 vs. S6 |
| *Monoglobus* | Firmicutes | 2.321 | 0.000 | 0.005 | S1 vs. S4 |
| *Burkholderia-Caballeronia-Paraburkholderia* | Proteobacteria | 2.321 | 0.000 | 0.004 | S3 vs. S4 |
| *uncultured* | Proteobacteria | 2.321 | 0.000 | 0.004 | S3 vs. S4 |

**Note:** Data shows genera with significant differential abundance across sites.

### Table 12.Significant Differential Abundance of Genera Across Site Comparisons

*Filtered for Adjusted p-value < 0.05*

| Genus | Phylum | **Log2 Fold Change** | p-value | **Adjusted p-value** | Site Comparison |
|---|---|---|---|---|---|
| *Lactobacillus* | Firmicutes | 2.265 | 0.000 | 0.006 | S3 vs. S7a |
| *28-YEA-48* | Proteobacteria | 2.264 | 0.000 | 0.006 | S3 vs. S7a |
| *Lactobacillus* | Firmicutes | 2.263 | 0.000 | 0.005 | S3 vs. S6 |
| *Lactobacillus* | Firmicutes | 2.263 | 0.000 | 0.005 | S3 vs. S4 |
| *28-YEA-48* | Proteobacteria | 2.262 | 0.000 | 0.005 | S3 vs. S6 |
| *28-YEA-48* | Proteobacteria | 2.262 | 0.000 | 0.005 | S3 vs. S4 |
| *Collimonas* | Proteobacteria | 2.262 | 0.003 | 0.040 | S3 vs. S6 |
| *Collimonas* | Proteobacteria | 2.262 | 0.003 | 0.039 | S3 vs. S4 |
| *Pseudomonas* | Proteobacteria | 2.250 | 0.000 | 0.005 | S1 vs. S7a |
| | Proteobacteria | 2.250 | 0.000 | 0.005 | S1 vs. S7a |
| *Paracoccus* | Proteobacteria | 2.249 | 0.000 | 0.005 | S1 vs. S7a |
| *Candidatus* | Bacteroidota | 2.249 | 0.000 | 0.005 | S1 vs. S7a |
| | Bacteroidota | 2.249 | 0.000 | 0.005 | S1 vs. S7a |
| | Firmicutes | 2.249 | 0.000 | 0.005 | S1 vs. S7a |
| *Candidatus* | Firmicutes | 2.249 | 0.000 | 0.005 | S1 vs. S7a |
| *Deinococcus* | Deinococcota | 2.249 | 0.000 | 0.005 | S1 vs. S7a |
| | Bacteroidota | 2.249 | 0.000 | 0.005 | S1 vs. S7a |
| *Pseudomonas* | Proteobacteria | 2.249 | 0.000 | 0.005 | S1 vs. S7a |
| *Pseudomonas* | Proteobacteria | 2.249 | 0.007 | 0.046 | S1 vs. S3 |
| *Pseudomonas* | Proteobacteria | 2.248 | 0.000 | 0.006 | S1 vs. S6 |
| | Proteobacteria | 2.248 | 0.007 | 0.046 | S1 vs. S3 |
| *Pseudomonas* | Proteobacteria | 2.248 | 0.001 | 0.015 | S1 vs. S2 |

**Note:** Data shows genera with significant differential abundance across sites.

### Table 12.Significant Differential Abundance of Genera Across Site Comparisons

*Filtered for Adjusted p-value < 0.05*

| Genus | Phylum | **Log2 Fold Change** | p-value | **Adjusted p-value** | Site Comparison |
|---|---|---|---|---|---|
| *Pseudomonas* | Proteobacteria | 2.248 | 0.000 | 0.006 | S1 vs. S4 |
| | Proteobacteria | 2.248 | 0.000 | 0.006 | S1 vs. S6 |
| | Proteobacteria | 2.248 | 0.001 | 0.015 | S1 vs. S2 |
| | Proteobacteria | 2.248 | 0.000 | 0.006 | S1 vs. S4 |
| *Paracoccus* | Proteobacteria | 2.247 | 0.007 | 0.046 | S1 vs. S3 |
| *Candidatus* | Bacteroidota | 2.247 | 0.007 | 0.046 | S1 vs. S3 |
| | Bacteroidota | 2.247 | 0.007 | 0.046 | S1 vs. S3 |
| | Firmicutes | 2.247 | 0.007 | 0.046 | S1 vs. S3 |
| *Candidatus* | Firmicutes | 2.247 | 0.007 | 0.046 | S1 vs. S3 |
| *Deinococcus* | Deinococcota | 2.247 | 0.007 | 0.046 | S1 vs. S3 |
| | Bacteroidota | 2.247 | 0.007 | 0.046 | S1 vs. S3 |
| *Pseudomonas* | Proteobacteria | 2.247 | 0.007 | 0.046 | S1 vs. S3 |
| *Paracoccus* | Proteobacteria | 2.247 | 0.000 | 0.006 | S1 vs. S6 |
| *Candidatus* | Bacteroidota | 2.247 | 0.000 | 0.006 | S1 vs. S6 |
| | Bacteroidota | 2.247 | 0.000 | 0.006 | S1 vs. S6 |
| | Firmicutes | 2.247 | 0.000 | 0.006 | S1 vs. S6 |
| *Candidatus* | Firmicutes | 2.247 | 0.000 | 0.006 | S1 vs. S6 |
| *Deinococcus* | Deinococcota | 2.247 | 0.000 | 0.006 | S1 vs. S6 |
| | Bacteroidota | 2.247 | 0.000 | 0.006 | S1 vs. S6 |
| *Pseudomonas* | Proteobacteria | 2.247 | 0.000 | 0.006 | S1 vs. S6 |
| *Paracoccus* | Proteobacteria | 2.247 | 0.001 | 0.015 | S1 vs. S2 |
| *Candidatus* | Bacteroidota | 2.247 | 0.001 | 0.015 | S1 vs. S2 |

**Note:** Data shows genera with significant differential abundance across sites.

### Table 12. Significant Differential Abundance of Genera Across Site Comparisons

*Filtered for Adjusted p-value < 0.05*

| Genus | Phylum | **Log2 Fold Change** | p-value | **Adjusted p-value** | Site Comparison |
|---|---|---|---|---|---|
| | Bacteroidota | 2.247 | 0.001 | 0.015 | S1 vs. S2 |
| | Firmicutes | 2.247 | 0.001 | 0.015 | S1 vs. S2 |
| *Candidatus* | Firmicutes | 2.247 | 0.001 | 0.015 | S1 vs. S2 |
| *Deinococcus* | Deinococcota | 2.247 | 0.001 | 0.015 | S1 vs. S2 |
| | Bacteroidota | 2.247 | 0.001 | 0.015 | S1 vs. S2 |
| *Pseudomonas* | Proteobacteria | 2.247 | 0.001 | 0.015 | S1 vs. S2 |
| *Paracoccus* | Proteobacteria | 2.247 | 0.000 | 0.006 | S1 vs. S4 |
| *Candidatus* | Bacteroidota | 2.247 | 0.000 | 0.006 | S1 vs. S4 |
| | Bacteroidota | 2.247 | 0.000 | 0.006 | S1 vs. S4 |
| | Firmicutes | 2.247 | 0.000 | 0.006 | S1 vs. S4 |
| *Candidatus* | Firmicutes | 2.247 | 0.000 | 0.006 | S1 vs. S4 |
| *Deinococcus* | Deinococcota | 2.247 | 0.000 | 0.006 | S1 vs. S4 |
| | Bacteroidota | 2.247 | 0.000 | 0.006 | S1 vs. S4 |
| *Pseudomonas* | Proteobacteria | 2.247 | 0.000 | 0.006 | S1 vs. S4 |
| *Polaromonas* | Proteobacteria | 2.203 | 0.000 | 0.008 | S3 vs. S7a |
| *Acidisoma* | Proteobacteria | 2.203 | 0.000 | 0.008 | S3 vs. S7a |
| *Afipia* | Proteobacteria | 2.203 | 0.000 | 0.008 | S3 vs. S7a |
| *Acidothermus* | Actinobacteriota | 2.203 | 0.000 | 0.008 | S3 vs. S7a |
| *Methylovirgula* | Proteobacteria | 2.203 | 0.000 | 0.009 | S3 vs. S7a |
| *Rickettsiella* | Proteobacteria | 2.202 | 0.003 | 0.037 | S3 vs. S4 |
| *Polaromonas* | Proteobacteria | 2.202 | 0.000 | 0.006 | S3 vs. S6 |
| *Polaromonas* | Proteobacteria | 2.201 | 0.000 | 0.006 | S3 vs. S4 |

**Note:** Data shows genera with significant differential abundance across sites.

**Table 12.Significant Differential Abundance of Genera Across Site Comparisons**

*Filtered for Adjusted p-value < 0.05*

| Genus | Phylum | **Log2 Fold Change** | p-value | **Adjusted p-value** | Site Comparison |
|---|---|---|---|---|---|
| *Acidisoma* | Proteobacteria | 2.201 | 0.000 | 0.006 | S3 vs. S6 |
| *Afipia* | Proteobacteria | 2.201 | 0.000 | 0.006 | S3 vs. S6 |
| *Acidothermus* | Actinobacteriota | 2.201 | 0.000 | 0.006 | S3 vs. S6 |
| *Methylovirgula* | Proteobacteria | 2.201 | 0.000 | 0.007 | S3 vs. S6 |
| *Acidisoma* | Proteobacteria | 2.201 | 0.000 | 0.006 | S3 vs. S4 |
| *Afipia* | Proteobacteria | 2.201 | 0.000 | 0.006 | S3 vs. S4 |
| *Acidothermus* | Actinobacteriota | 2.201 | 0.000 | 0.006 | S3 vs. S4 |
| *Bacillus* | Firmicutes | 2.200 | 0.004 | 0.047 | S3 vs. S4 |
| *Spiroplasma* | Firmicutes | 2.172 | 0.000 | 0.007 | S1 vs. S7a |
| | Proteobacteria | 2.171 | 0.000 | 0.007 | S1 vs. S7a |
| *Spiroplasma* | Firmicutes | 2.171 | 0.000 | 0.007 | S1 vs. S7a |
| *Pseudomonas* | Proteobacteria | 2.171 | 0.002 | 0.031 | S1 vs. S7a |
| *Spiroplasma* | Firmicutes | 2.170 | 0.000 | 0.009 | S1 vs. S6 |
| *Spiroplasma* | Firmicutes | 2.170 | 0.001 | 0.020 | S1 vs. S2 |
| *Spiroplasma* | Firmicutes | 2.170 | 0.000 | 0.009 | S1 vs. S4 |
| | Proteobacteria | 2.169 | 0.000 | 0.009 | S1 vs. S6 |
| *Spiroplasma* | Firmicutes | 2.169 | 0.000 | 0.009 | S1 vs. S6 |
| | Proteobacteria | 2.169 | 0.001 | 0.020 | S1 vs. S2 |
| *Spiroplasma* | Firmicutes | 2.169 | 0.001 | 0.020 | S1 vs. S2 |
| | Proteobacteria | 2.169 | 0.000 | 0.009 | S1 vs. S4 |
| *Spiroplasma* | Firmicutes | 2.169 | 0.000 | 0.009 | S1 vs. S4 |
| *Pseudomonas* | Proteobacteria | 2.169 | 0.002 | 0.036 | S1 vs. S6 |

**Note:** Data shows genera with significant differential abundance across sites.

### Table 12.Significant Differential Abundance of Genera Across Site Comparisons

*Filtered for Adjusted p-value < 0.05*

| Genus | Phylum | **Log2 Fold Change** | p-value | **Adjusted p-value** | Site Comparison |
|---|---|---|---|---|---|
| *Pseudomonas* | Proteobacteria | 2.169 | 0.002 | 0.036 | S1 vs. S4 |
| *Pedobacter* | Bacteroidota | 2.138 | 0.000 | 0.010 | S3 vs. S7a |
| *Acinetobacter* | Proteobacteria | 2.138 | 0.000 | 0.010 | S3 vs. S7a |
| *Ac37b* | Proteobacteria | 2.138 | 0.000 | 0.032 | S2 vs. S7a |
| *Pedobacter* | Bacteroidota | 2.137 | 0.000 | 0.008 | S3 vs. S6 |
| *Acinetobacter* | Proteobacteria | 2.137 | 0.000 | 0.008 | S3 vs. S6 |
| *Ac37b* | Proteobacteria | 2.137 | 0.001 | 0.043 | S2 vs. S6 |
| *Pedobacter* | Bacteroidota | 2.136 | 0.000 | 0.008 | S3 vs. S4 |
| *Acinetobacter* | Proteobacteria | 2.136 | 0.000 | 0.008 | S3 vs. S4 |
| *Ac37b* | Proteobacteria | 2.136 | 0.001 | 0.044 | S2 vs. S4 |
| *Streptococcus* | Firmicutes | 2.106 | 0.000 | 0.032 | S2 vs. S7a |
| *Streptococcus* | Firmicutes | 2.105 | 0.001 | 0.043 | S2 vs. S6 |
| *Streptococcus* | Firmicutes | 2.104 | 0.001 | 0.044 | S2 vs. S4 |
| *Papillibacter* | Firmicutes | 2.089 | 0.001 | 0.011 | S1 vs. S7a |
| *Hydrogenophilus* | Proteobacteria | 2.089 | 0.001 | 0.011 | S1 vs. S7a |
| *Papillibacter* | Firmicutes | 2.087 | 0.001 | 0.014 | S1 vs. S6 |
| *Hydrogenophilus* | Proteobacteria | 2.087 | 0.001 | 0.014 | S1 vs. S6 |
| *Papillibacter* | Firmicutes | 2.087 | 0.002 | 0.030 | S1 vs. S2 |
| *Hydrogenophilus* | Proteobacteria | 2.087 | 0.002 | 0.030 | S1 vs. S2 |
| *Papillibacter* | Firmicutes | 2.087 | 0.001 | 0.014 | S1 vs. S4 |
| *Hydrogenophilus* | Proteobacteria | 2.087 | 0.001 | 0.014 | S1 vs. S4 |
| *Sphingopyxis* | Proteobacteria | 2.072 | 0.001 | 0.032 | S2 vs. S7a |

**Note:** Data shows genera with significant differential abundance across sites.

### Table 12.Significant Differential Abundance of Genera Across Site Comparisons

*Filtered for Adjusted p-value < 0.05*

| Genus | Phylum | **Log2 Fold Change** | p-value | **Adjusted p-value** | Site Comparison |
|---|---|---|---|---|---|
| | Proteobacteria | 2.072 | 0.001 | 0.032 | S2 vs. S7a |
| *Acidisoma* | Proteobacteria | 2.072 | 0.001 | 0.013 | S3 vs. S7a |
| *uncultured* | Proteobacteria | 2.072 | 0.001 | 0.013 | S3 vs. S7a |
| *Corynebacterium* | Actinobacteriota | 2.072 | 0.001 | 0.013 | S3 vs. S7a |
| *uncultured* | Proteobacteria | 2.071 | 0.001 | 0.013 | S3 vs. S7a |
| *Acinetobacter* | Proteobacteria | 2.071 | 0.001 | 0.013 | S3 vs. S7a |
| | Proteobacteria | 2.071 | 0.001 | 0.043 | S2 vs. S6 |
| *Sphingopyxis* | Proteobacteria | 2.071 | 0.001 | 0.043 | S2 vs. S6 |
| | Proteobacteria | 2.070 | 0.001 | 0.044 | S2 vs. S4 |
| *Sphingopyxis* | Proteobacteria | 2.070 | 0.001 | 0.044 | S2 vs. S4 |
| *uncultured* | Proteobacteria | 2.070 | 0.001 | 0.010 | S3 vs. S6 |
| *Acidisoma* | Proteobacteria | 2.070 | 0.001 | 0.010 | S3 vs. S6 |
| *Corynebacterium* | Actinobacteriota | 2.070 | 0.001 | 0.010 | S3 vs. S6 |
| *uncultured* | Proteobacteria | 2.070 | 0.001 | 0.010 | S3 vs. S4 |
| *Acidisoma* | Proteobacteria | 2.070 | 0.001 | 0.010 | S3 vs. S4 |
| *Corynebacterium* | Actinobacteriota | 2.070 | 0.001 | 0.010 | S3 vs. S4 |
| *uncultured* | Proteobacteria | 2.070 | 0.001 | 0.010 | S3 vs. S6 |
| *Acinetobacter* | Proteobacteria | 2.070 | 0.001 | 0.010 | S3 vs. S6 |
| *uncultured* | Proteobacteria | 2.069 | 0.001 | 0.010 | S3 vs. S4 |
| *Acinetobacter* | Proteobacteria | 2.069 | 0.001 | 0.010 | S3 vs. S4 |
| *Bacillus* | Firmicutes | 2.038 | 0.002 | 0.047 | S2 vs. S7a |
| | Actinobacteriota | 2.001 | 0.001 | 0.018 | S1 vs. S7a |

**Note:** Data shows genera with significant differential abundance across sites.

**Table 12.Significant Differential Abundance of Genera Across Site Comparisons**

*Filtered for Adjusted p-value < 0.05*

| Genus | Phylum | **Log2 Fold Change** | p-value | **Adjusted p-value** | Site Comparison |
|---|---|---|---|---|---|
| | Firmicutes | 2.001 | 0.001 | 0.018 | S1 vs. S7a |
| *Ereboglobus* | Verrucomicrobiota | 2.001 | 0.001 | 0.018 | S1 vs. S7a |
| *Ac37b* | Proteobacteria | 2.001 | 0.001 | 0.034 | S2 vs. S7a |
| *Ac37b* | Proteobacteria | 2.001 | 0.001 | 0.034 | S2 vs. S7a |
| *Ac37b* | Proteobacteria | 2.001 | 0.001 | 0.034 | S2 vs. S7a |
| | Actinobacteriota | 1.999 | 0.001 | 0.022 | S1 vs. S6 |
| | Firmicutes | 1.999 | 0.001 | 0.022 | S1 vs. S6 |
| *Ereboglobus* | Verrucomicrobiota | 1.999 | 0.001 | 0.022 | S1 vs. S6 |
| | Actinobacteriota | 1.999 | 0.003 | 0.044 | S1 vs. S2 |
| | Firmicutes | 1.999 | 0.003 | 0.044 | S1 vs. S2 |
| *Ereboglobus* | Verrucomicrobiota | 1.999 | 0.003 | 0.044 | S1 vs. S2 |
| | Actinobacteriota | 1.999 | 0.001 | 0.022 | S1 vs. S4 |
| | Firmicutes | 1.999 | 0.001 | 0.022 | S1 vs. S4 |
| *Ereboglobus* | Verrucomicrobiota | 1.999 | 0.001 | 0.022 | S1 vs. S4 |
| *Ac37b* | Proteobacteria | 1.999 | 0.001 | 0.043 | S2 vs. S6 |
| *Ac37b* | Proteobacteria | 1.999 | 0.001 | 0.043 | S2 vs. S6 |
| *Ac37b* | Proteobacteria | 1.999 | 0.001 | 0.043 | S2 vs. S6 |
| *Pseudomonas* | Proteobacteria | 1.999 | 0.003 | 0.039 | S3 vs. S6 |
| *Ac37b* | Proteobacteria | 1.999 | 0.001 | 0.044 | S2 vs. S4 |
| *Ac37b* | Proteobacteria | 1.999 | 0.001 | 0.044 | S2 vs. S4 |
| *Ac37b* | Proteobacteria | 1.999 | 0.001 | 0.044 | S2 vs. S4 |
| *Pseudomonas* | Proteobacteria | 1.999 | 0.003 | 0.038 | S3 vs. S4 |

**Note:** Data shows genera with significant differential abundance across sites.

### Table 12.Significant Differential Abundance of Genera Across Site Comparisons

*Filtered for Adjusted p-value < 0.05*

| Genus | Phylum | **Log2 Fold Change** | p-value | **Adjusted p-value** | Site Comparison |
|---|---|---|---|---|---|
| | Actinobacteriota | 1.928 | 0.001 | 0.043 | S2 vs. S7a |
| | Proteobacteria | 1.927 | 0.001 | 0.028 | S3 vs. S7a |
| | Bacteroidota | 1.927 | 0.001 | 0.028 | S3 vs. S7a |
| | Proteobacteria | 1.925 | 0.001 | 0.020 | S3 vs. S6 |
| | Bacteroidota | 1.925 | 0.001 | 0.020 | S3 vs. S6 |
| | Proteobacteria | 1.925 | 0.001 | 0.020 | S3 vs. S4 |
| | Bacteroidota | 1.925 | 0.001 | 0.020 | S3 vs. S4 |
| | Actinobacteriota | 1.908 | 0.002 | 0.030 | S1 vs. S7a |
| *uncultured* | Bacteroidota | 1.908 | 0.002 | 0.030 | S1 vs. S7a |
| | Actinobacteriota | 1.906 | 0.002 | 0.036 | S1 vs. S6 |
| *uncultured* | Bacteroidota | 1.906 | 0.002 | 0.036 | S1 vs. S6 |
| | Actinobacteriota | 1.906 | 0.002 | 0.036 | S1 vs. S4 |
| *uncultured* | Bacteroidota | 1.906 | 0.002 | 0.036 | S1 vs. S4 |
| *Methylovirgula* | Proteobacteria | 1.901 | 0.002 | 0.021 | S3 vs. S4 |
| *Burkholderia-Caballeronia-Paraburkholderia* | Proteobacteria | 1.890 | 0.001 | 0.045 | S2 vs. S7a |
| *Rhodopila* | Proteobacteria | 1.890 | 0.001 | 0.045 | S2 vs. S7a |
| | WPS-2 | 1.884 | 0.002 | 0.047 | S6 vs. S7a |
| *Spiroplasma* | Firmicutes | 1.856 | 0.002 | 0.047 | S6 vs. S7a |
| *Escherichia-Shigella* | Proteobacteria | 1.856 | 0.002 | 0.047 | S6 vs. S7a |
| *Acidocella* | Proteobacteria | 1.850 | 0.002 | 0.047 | S2 vs. S7a |
| *Acidothermus* | Actinobacteriota | 1.849 | 0.002 | 0.042 | S3 vs. S7a |

**Note:** Data shows genera with significant differential abundance across sites.

### Table 12.Significant Differential Abundance of Genera Across Site Comparisons

*Filtered for Adjusted p-value < 0.05*

| Genus | Phylum | **Log2 Fold Change** | p-value | **Adjusted p-value** | Site Comparison |
|---|---|---|---|---|---|
| *Acidothermus* | Actinobacteriota | 1.847 | 0.002 | 0.030 | S3 vs. S6 |
| *Acidothermus* | Actinobacteriota | 1.847 | 0.002 | 0.030 | S3 vs. S4 |
| | Proteobacteria | 1.823 | 0.002 | 0.047 | S6 vs. S7a |
| *Burkholderia-Caballeronia-Paraburkholderia* | Proteobacteria | 1.823 | 0.007 | 0.047 | S6 vs. S7a |
| *Sphingomonas* | Proteobacteria | 1.765 | 0.004 | 0.042 | S3 vs. S6 |
| | Proteobacteria | 1.765 | 0.004 | 0.042 | S3 vs. S6 |
| *Mycobacterium* | Actinobacteriota | 1.765 | 0.004 | 0.042 | S3 vs. S6 |
| *Dyella* | Proteobacteria | 1.765 | 0.004 | 0.042 | S3 vs. S6 |
| *Sphingomonas* | Proteobacteria | 1.765 | 0.004 | 0.041 | S3 vs. S4 |
| | Proteobacteria | 1.765 | 0.004 | 0.041 | S3 vs. S4 |
| *Mycobacterium* | Actinobacteriota | 1.765 | 0.004 | 0.041 | S3 vs. S4 |
| *Dyella* | Proteobacteria | 1.765 | 0.004 | 0.041 | S3 vs. S4 |
| *Aquisphaera* | Planctomycetota | 1.726 | 0.003 | 0.047 | S6 vs. S7a |
| | Acidobacteriota | 1.726 | 0.003 | 0.047 | S6 vs. S7a |
| | Proteobacteria | 1.692 | 0.004 | 0.047 | S6 vs. S7a |
| *Methyloferula* | Proteobacteria | 1.657 | 0.004 | 0.047 | S6 vs. S7a |
| *Acidothermus* | Actinobacteriota | 1.657 | 0.004 | 0.047 | S6 vs. S7a |
| *uncultured* | Planctomycetota | 1.657 | 0.004 | 0.047 | S6 vs. S7a |
| | Proteobacteria | 1.657 | 0.004 | 0.047 | S6 vs. S7a |
| | Acidobacteriota | 1.657 | 0.006 | 0.047 | S6 vs. S7a |
| *uncultured* | Planctomycetota | 1.622 | 0.005 | 0.047 | S6 vs. S7a |

**Note:** Data shows genera with significant differential abundance across sites.

### Table 12.Significant Differential Abundance of Genera Across Site Comparisons

*Filtered for Adjusted p-value < 0.05*

| Genus | Phylum | **Log2 Fold Change** | p-value | **Adjusted p-value** | Site Comparison |
|---|---|---|---|---|---|
| *uncultured* | Planctomycetota | 1.622 | 0.005 | 0.047 | S6 vs. S7a |
| *Acidothermus* | Actinobacteriota | 1.585 | 0.006 | 0.047 | S6 vs. S7a |
| *uncultured* | Planctomycetota | 1.585 | 0.006 | 0.047 | S6 vs. S7a |
| | Proteobacteria | 1.585 | 0.006 | 0.047 | S6 vs. S7a |
| *Aquisphaera* | Planctomycetota | 1.548 | 0.006 | 0.047 | S6 vs. S7a |
| | Proteobacteria | 1.548 | 0.006 | 0.047 | S6 vs. S7a |
| *Leuconostoc* | Firmicutes | 1.511 | 0.008 | 0.047 | S6 vs. S7a |
| *uncultured* | Actinobacteriota | 1.509 | 0.008 | 0.047 | S6 vs. S7a |
| | Acidobacteriota | 1.509 | 0.008 | 0.047 | S6 vs. S7a |
| *Aquisphaera* | Planctomycetota | 1.470 | 0.009 | 0.048 | S6 vs. S7a |
| *GOUTA6* | Proteobacteria | −1.504 | 0.010 | 0.048 | S6 vs. S7a |
| *uncultured* | Proteobacteria | −1.504 | 0.010 | 0.048 | S6 vs. S7a |
| | Actinobacteriota | −1.504 | 0.010 | 0.048 | S6 vs. S7a |
| | Actinobacteriota | −1.504 | 0.010 | 0.048 | S6 vs. S7a |
| *Mycobacterium* | Actinobacteriota | −1.504 | 0.010 | 0.048 | S6 vs. S7a |
| *Candidatus* | Acidobacteriota | −1.504 | 0.010 | 0.048 | S6 vs. S7a |
| | Acidobacteriota | −1.504 | 0.010 | 0.048 | S6 vs. S7a |
| *uncultured* | Gemmatimonadota | −1.504 | 0.010 | 0.048 | S6 vs. S7a |
| *Dyella* | Proteobacteria | −1.504 | 0.010 | 0.048 | S6 vs. S7a |
| *Ac37b* | Proteobacteria | −1.513 | 0.009 | 0.048 | S6 vs. S7a |
| *Ac37b* | Proteobacteria | −1.513 | 0.009 | 0.048 | S6 vs. S7a |
| *Ac37b* | Proteobacteria | −1.513 | 0.009 | 0.048 | S6 vs. S7a |

**Note:** Data shows genera with significant differential abundance across sites.

**Table 12.Significant Differential Abundance of Genera Across Site Comparisons**

*Filtered for Adjusted p-value < 0.05*

| Genus | Phylum | **Log2 Fold Change** | p-value | **Adjusted p-value** | Site Comparison |
|---|---|---|---|---|---|
| | Proteobacteria | −1.539 | 0.008 | 0.047 | S6 vs. S7a |
| *uncultured* | Proteobacteria | −1.539 | 0.008 | 0.047 | S6 vs. S7a |
| *uncultured* | Proteobacteria | −1.539 | 0.008 | 0.047 | S6 vs. S7a |
| | Proteobacteria | −1.539 | 0.008 | 0.047 | S6 vs. S7a |
| *uncultured* | Proteobacteria | −1.539 | 0.008 | 0.047 | S6 vs. S7a |
| *uncultured* | Verrucomicrobiota | −1.539 | 0.008 | 0.047 | S6 vs. S7a |
| *Pedosphaera* | Verrucomicrobiota | −1.539 | 0.008 | 0.047 | S6 vs. S7a |
| *Opitutus* | Verrucomicrobiota | −1.539 | 0.008 | 0.047 | S6 vs. S7a |
| *Bryobacter* | Acidobacteriota | −1.539 | 0.008 | 0.047 | S6 vs. S7a |
| *Ac37b* | Proteobacteria | −1.548 | 0.008 | 0.047 | S6 vs. S7a |
| *Weissella* | Firmicutes | −1.549 | 0.008 | 0.047 | S6 vs. S7a |
| | Acidobacteriota | −1.549 | 0.008 | 0.047 | S6 vs. S7a |
| *uncultured* | Proteobacteria | −1.574 | 0.007 | 0.047 | S6 vs. S7a |
| | Actinobacteriota | −1.574 | 0.007 | 0.047 | S6 vs. S7a |
| *Acidothermus* | Actinobacteriota | −1.574 | 0.007 | 0.047 | S6 vs. S7a |
| | Acidobacteriota | −1.574 | 0.007 | 0.047 | S6 vs. S7a |
| | Acidobacteriota | −1.574 | 0.007 | 0.047 | S6 vs. S7a |
| *Flavobacterium* | Bacteroidota | −1.574 | 0.007 | 0.047 | S6 vs. S7a |
| *Puia* | Bacteroidota | −1.574 | 0.007 | 0.047 | S6 vs. S7a |
| *Pseudomonas* | Proteobacteria | −1.574 | 0.007 | 0.047 | S6 vs. S7a |
| *Ac37b* | Proteobacteria | −1.583 | 0.007 | 0.047 | S6 vs. S7a |
| *Ac37b* | Proteobacteria | −1.583 | 0.007 | 0.047 | S6 vs. S7a |

**Note:** Data shows genera with significant differential abundance across sites.

### Table 12. Significant Differential Abundance of Genera Across Site Comparisons

*Filtered for Adjusted p-value < 0.05*

| Genus | Phylum | **Log2 Fold Change** | p-value | **Adjusted p-value** | Site Comparison |
|---|---|---|---|---|---|
| *Rothia* | Actinobacteriota | −1.584 | 0.007 | 0.047 | S6 vs. S7a |
| *Burkholderia-Caballeronia-Paraburkholderia* | Proteobacteria | −1.607 | 0.006 | 0.047 | S6 vs. S7a |
| *uncultured* | Proteobacteria | −1.607 | 0.006 | 0.047 | S6 vs. S7a |
| *Phenylobacterium* | Proteobacteria | −1.607 | 0.006 | 0.047 | S6 vs. S7a |
| *Mucilaginibacter* | Bacteroidota | −1.607 | 0.006 | 0.047 | S6 vs. S7a |
| *Ac37b* | Proteobacteria | −1.617 | 0.006 | 0.047 | S6 vs. S7a |
| | WPS-2 | −1.640 | 0.007 | 0.047 | S6 vs. S7a |
| *Pandoraea* | Proteobacteria | −1.640 | 0.006 | 0.047 | S6 vs. S7a |
| | Proteobacteria | −1.640 | 0.006 | 0.047 | S6 vs. S7a |
| *Candidatus* | Acidobacteriota | −1.640 | 0.006 | 0.047 | S6 vs. S7a |
| | Acidobacteriota | −1.640 | 0.006 | 0.047 | S6 vs. S7a |
| *Puia* | Bacteroidota | −1.640 | 0.006 | 0.047 | S6 vs. S7a |
| *Inquilinus* | Proteobacteria | −1.672 | 0.005 | 0.047 | S6 vs. S7a |
| *Acidocella* | Proteobacteria | −1.672 | 0.005 | 0.047 | S6 vs. S7a |
| | Proteobacteria | −1.672 | 0.005 | 0.047 | S6 vs. S7a |
| *uncultured* | Proteobacteria | −1.672 | 0.005 | 0.047 | S6 vs. S7a |
| *uncultured* | Verrucomicrobiota | −1.672 | 0.005 | 0.047 | S6 vs. S7a |
| *uncultured* | Verrucomicrobiota | −1.672 | 0.005 | 0.047 | S6 vs. S7a |
| *Bryobacter* | Acidobacteriota | −1.672 | 0.005 | 0.047 | S6 vs. S7a |
| *Rhodanobacter* | Proteobacteria | −1.672 | 0.005 | 0.047 | S6 vs. S7a |
| *Ac37b* | Proteobacteria | −1.683 | 0.005 | 0.047 | S6 vs. S7a |

**Note:** Data shows genera with significant differential abundance across sites.

### Table 12.Significant Differential Abundance of Genera Across Site Comparisons

*Filtered for Adjusted p-value < 0.05*

| Genus | Phylum | **Log2 Fold Change** | p-value | **Adjusted p-value** | Site Comparison |
|---|---|---|---|---|---|
| *Bacillus* | Firmicutes | −1.684 | 0.005 | 0.047 | S6 vs. S7a |
| *Acidothermus* | Actinobacteriota | −1.704 | 0.004 | 0.047 | S6 vs. S7a |
| *Schlesneria* | Planctomycetota | −1.704 | 0.004 | 0.047 | S6 vs. S7a |
|  | Acidobacteriota | −1.704 | 0.004 | 0.047 | S6 vs. S7a |
| *Rudaea* | Proteobacteria | −1.704 | 0.004 | 0.047 | S6 vs. S7a |
|  | Proteobacteria | −1.734 | 0.004 | 0.047 | S6 vs. S7a |
| *uncultured* | Verrucomicrobiota | −1.734 | 0.004 | 0.047 | S6 vs. S7a |
| *Opitutus* | Verrucomicrobiota | −1.734 | 0.004 | 0.047 | S6 vs. S7a |
|  | Acidobacteriota | −1.734 | 0.004 | 0.047 | S6 vs. S7a |
|  | Acidobacteriota | −1.734 | 0.004 | 0.047 | S6 vs. S7a |
| *Puia* | Bacteroidota | −1.734 | 0.004 | 0.047 | S6 vs. S7a |
|  | Proteobacteria | −1.734 | 0.004 | 0.047 | S6 vs. S7a |
| *Ac37b* | Proteobacteria | −1.745 | 0.004 | 0.047 | S6 vs. S7a |
| *Ac37b* | Proteobacteria | −1.745 | 0.004 | 0.047 | S6 vs. S7a |
|  | Proteobacteria | −1.746 | 0.004 | 0.047 | S6 vs. S7a |
| *Mycobacterium* | Actinobacteriota | −1.763 | 0.009 | 0.048 | S6 vs. S7a |
| *GOUTA6* | Proteobacteria | −1.765 | 0.003 | 0.047 | S6 vs. S7a |
| *Novosphingobium* | Proteobacteria | −1.765 | 0.003 | 0.047 | S6 vs. S7a |
|  | Proteobacteria | −1.765 | 0.003 | 0.047 | S6 vs. S7a |
| *Mucilaginibacter* | Bacteroidota | −1.765 | 0.003 | 0.047 | S6 vs. S7a |
| *uncultured* | Bacteroidota | −1.765 | 0.003 | 0.047 | S6 vs. S7a |
| *Ferruginibacter* | Bacteroidota | −1.765 | 0.003 | 0.047 | S6 vs. S7a |

**Note:** Data shows genera with significant differential abundance across sites.

### Table 12.Significant Differential Abundance of Genera Across Site Comparisons

*Filtered for Adjusted p-value < 0.05*

| Genus | Phylum | **Log2 Fold Change** | p-value | **Adjusted p-value** | Site Comparison |
|---|---|---|---|---|---|
| *Rhodanobacter* | Proteobacteria | −1.765 | 0.003 | 0.047 | S6 vs. S7a |
| *Ac37b* | Proteobacteria | −1.776 | 0.003 | 0.047 | S6 vs. S7a |
| *Ac37b* | Proteobacteria | −1.776 | 0.003 | 0.047 | S6 vs. S7a |
| *Ac37b* | Proteobacteria | −1.776 | 0.003 | 0.047 | S6 vs. S7a |
| *Ac37b* | Proteobacteria | −1.776 | 0.003 | 0.047 | S6 vs. S7a |
| | Proteobacteria | −1.777 | 0.003 | 0.047 | S6 vs. S7a |
| *uncultured* | Proteobacteria | −1.794 | 0.003 | 0.047 | S6 vs. S7a |
| *Labrys* | Proteobacteria | −1.794 | 0.003 | 0.047 | S6 vs. S7a |
| *Bryobacter* | Acidobacteriota | −1.794 | 0.003 | 0.047 | S6 vs. S7a |
| | Acidobacteriota | −1.794 | 0.003 | 0.047 | S6 vs. S7a |
| | Acidobacteriota | −1.794 | 0.003 | 0.047 | S6 vs. S7a |
| | Proteobacteria | −1.806 | 0.003 | 0.047 | S6 vs. S7a |
| *Ktedonobacter* | Chloroflexi | −1.823 | 0.003 | 0.047 | S6 vs. S7a |
| | Verrucomicrobiota | −1.823 | 0.003 | 0.047 | S6 vs. S7a |
| *Staphylococcus* | Firmicutes | −1.836 | 0.010 | 0.049 | S6 vs. S7a |
| | Proteobacteria | −1.925 | 0.003 | 0.048 | S2 vs. S3 |
| | Bacteroidota | −1.925 | 0.003 | 0.048 | S2 vs. S3 |
| *Burkholderia-Caballeronia-Paraburkholderia* | Proteobacteria | −1.989 | 0.001 | 0.047 | S6 vs. S7a |
| *uncultured* | Proteobacteria | −2.069 | 0.002 | 0.025 | S2 vs. S3 |
| *Acinetobacter* | Proteobacteria | −2.069 | 0.002 | 0.025 | S2 vs. S3 |
| *uncultured* | Proteobacteria | −2.070 | 0.002 | 0.025 | S2 vs. S3 |

**Note:** Data shows genera with significant differential abundance across sites.

### Table 12. Significant Differential Abundance of Genera Across Site Comparisons

*Filtered for Adjusted p-value < 0.05*

| Genus | Phylum | **Log2 Fold Change** | p-value | **Adjusted p-value** | Site Comparison |
|---|---|---|---|---|---|
| *Acidisoma* | Proteobacteria | −2.070 | 0.002 | 0.025 | S2 vs. S3 |
| *Corynebacterium* | Actinobacteriota | −2.070 | 0.002 | 0.025 | S2 vs. S3 |
| *Enhydrobacter* | Proteobacteria | −2.083 | 0.001 | 0.025 | S2 vs. S3 |
| *Pedobacter* | Bacteroidota | −2.136 | 0.001 | 0.021 | S2 vs. S3 |
| *Acinetobacter* | Proteobacteria | −2.136 | 0.001 | 0.021 | S2 vs. S3 |
| *Methylovirgula* | Proteobacteria | −2.201 | 0.001 | 0.020 | S2 vs. S3 |
| *Acidisoma* | Proteobacteria | −2.201 | 0.001 | 0.017 | S2 vs. S3 |
| *Afipia* | Proteobacteria | −2.201 | 0.001 | 0.017 | S2 vs. S3 |
| *Acidothermus* | Actinobacteriota | −2.201 | 0.001 | 0.017 | S2 vs. S3 |
| *Polaromonas* | Proteobacteria | −2.201 | 0.001 | 0.017 | S2 vs. S3 |
| *Cloacibacterium* | Bacteroidota | −2.254 | 0.001 | 0.044 | S2 vs. S4 |
| *28-YEA-48* | Proteobacteria | −2.262 | 0.001 | 0.014 | S2 vs. S3 |
| *Lactobacillus* | Firmicutes | −2.263 | 0.001 | 0.014 | S2 vs. S3 |
| *Burkholderia-Caballeronia-Paraburkholderia* | Proteobacteria | −2.321 | 0.001 | 0.012 | S2 vs. S3 |
| *uncultured* | Proteobacteria | −2.321 | 0.001 | 0.012 | S2 vs. S3 |
| | Acidobacteriota | −2.322 | 0.001 | 0.012 | S2 vs. S3 |
| *Spiroplasma* | Firmicutes | −2.415 | 0.000 | 0.016 | S4 vs. S8a |
| *Spiroplasma* | Firmicutes | −2.415 | 0.000 | 0.020 | S2 vs. S8a |
| *Spiroplasma* | Firmicutes | −2.416 | 0.000 | 0.013 | S6 vs. S8a |
| *Spiroplasma* | Firmicutes | −2.417 | 0.000 | 0.008 | S7a vs. S8a |
| *Cutibacterium* | Actinobacteriota | −2.491 | 0.008 | 0.047 | S6 vs. S7a |

**Note:** Data shows genera with significant differential abundance across sites.

### Table 12.Significant Differential Abundance of Genera Across Site Comparisons

*Filtered for Adjusted p-value < 0.05*

| Genus | Phylum | **Log2 Fold Change** | p-value | **Adjusted p-value** | Site Comparison |
|---|---|---|---|---|---|
| *Ac37b* | Proteobacteria | −2.514 | 0.000 | 0.032 | S2 vs. S7a |
| *Ac37b* | Proteobacteria | −2.514 | 0.000 | 0.015 | S6 vs. S7a |
| *Spiroplasma* | Firmicutes | −2.536 | 0.000 | 0.005 | S2 vs. S3 |
| *Spiroplasma* | Firmicutes | −2.536 | 0.006 | 0.046 | S1 vs. S3 |
| | Actinobacteriota | −2.583 | 0.001 | 0.025 | S2 vs. S3 |
| | Actinobacteriota | −2.631 | 0.000 | 0.003 | S2 vs. S3 |
| | Actinobacteriota | −2.631 | 0.005 | 0.040 | S1 vs. S3 |
| *Sphingomonas* | Proteobacteria | −2.675 | 0.000 | 0.043 | S2 vs. S6 |
| *uncultured* | Proteobacteria | −2.677 | 0.000 | 0.003 | S2 vs. S3 |
| *Pseudomonas* | Proteobacteria | −2.677 | 0.000 | 0.003 | S2 vs. S3 |
| *uncultured* | Proteobacteria | −2.677 | 0.005 | 0.036 | S1 vs. S3 |
| *Pseudomonas* | Proteobacteria | −2.677 | 0.005 | 0.036 | S1 vs. S3 |
| | | −2.684 | 0.002 | 0.047 | S6 vs. S7a |
| *Staphylococcus* | Firmicutes | −2.700 | 0.000 | 0.006 | S4 vs. S5 |
| *Pseudomonas* | Proteobacteria | −2.700 | 0.000 | 0.006 | S4 vs. S5 |
| *Staphylococcus* | Firmicutes | −2.700 | 0.000 | 0.011 | S2 vs. S5 |
| *Pseudomonas* | Proteobacteria | −2.700 | 0.000 | 0.011 | S2 vs. S5 |
| | | −2.721 | 0.000 | 0.003 | S2 vs. S3 |
| *uncultured* | Verrucomicrobiota | −2.721 | 0.000 | 0.003 | S2 vs. S3 |
| | | −2.721 | 0.004 | 0.034 | S1 vs. S3 |
| *uncultured* | Verrucomicrobiota | −2.721 | 0.004 | 0.034 | S1 vs. S3 |
| *Brevundimonas* | Proteobacteria | −2.722 | 0.000 | 0.006 | S2 vs. S3 |

**Note:** Data shows genera with significant differential abundance across sites.

### Table 12.Significant Differential Abundance of Genera Across Site Comparisons

*Filtered for Adjusted p-value < 0.05*

| Genus | Phylum | **Log2 Fold Change** | p-value | **Adjusted p-value** | Site Comparison |
|---|---|---|---|---|---|
| *Brevundimonas* | Proteobacteria | −2.723 | 0.007 | 0.046 | S1 vs. S3 |
| *Conexibacter* | Actinobacteriota | −2.764 | 0.000 | 0.002 | S2 vs. S3 |
| | Dependentiae | −2.764 | 0.000 | 0.002 | S2 vs. S3 |
| | Planctomycetota | −2.764 | 0.000 | 0.002 | S2 vs. S3 |
| *Conexibacter* | Actinobacteriota | −2.764 | 0.004 | 0.032 | S1 vs. S3 |
| | Dependentiae | −2.764 | 0.004 | 0.032 | S1 vs. S3 |
| | Planctomycetota | −2.764 | 0.004 | 0.032 | S1 vs. S3 |
| *Spiroplasma* | Firmicutes | −2.907 | 0.000 | 0.002 | S4 vs. S5 |
| *Spiroplasma* | Firmicutes | −2.907 | 0.000 | 0.005 | S2 vs. S5 |
| *Rhodovastum* | Proteobacteria | −2.926 | 0.000 | 0.001 | S2 vs. S3 |
| *Micrococcus* | Actinobacteriota | −2.926 | 0.000 | 0.001 | S2 vs. S3 |
| *Micrococcus* | Actinobacteriota | −2.926 | 0.002 | 0.023 | S1 vs. S3 |
| *Rhodovastum* | Proteobacteria | −2.926 | 0.002 | 0.023 | S1 vs. S3 |
| *Acidisphaera* | Proteobacteria | −2.999 | 0.000 | 0.001 | S2 vs. S3 |
| *Acidisphaera* | Proteobacteria | −2.999 | 0.002 | 0.021 | S1 vs. S3 |
| *Diplorickettsia* | Proteobacteria | −3.059 | 0.000 | 0.000 | S4 vs. S8a |
| *Diplorickettsia* | Proteobacteria | −3.059 | 0.000 | 0.001 | S2 vs. S8a |
| *Diplorickettsia* | Proteobacteria | −3.059 | 0.000 | 0.000 | S6 vs. S8a |
| *Diplorickettsia* | Proteobacteria | −3.059 | 0.001 | 0.023 | S1 vs. S8a |
| *Diplorickettsia* | Proteobacteria | −3.060 | 0.000 | 0.008 | S3 vs. S8a |
| *Diplorickettsia* | Proteobacteria | −3.061 | 0.000 | 0.000 | S7a vs. S8a |
| *Chthoniobacter* | Verrucomicrobiota | −3.069 | 0.000 | 0.001 | S2 vs. S3 |

**Note:** Data shows genera with significant differential abundance across sites.

### Table 12.Significant Differential Abundance of Genera Across Site Comparisons

*Filtered for Adjusted p-value < 0.05*

| Genus | Phylum | **Log2 Fold Change** | p-value | **Adjusted p-value** | Site Comparison |
|---|---|---|---|---|---|
| *Chthoniobacter* | Verrucomicrobiota | −3.069 | 0.002 | 0.019 | S1 vs. S3 |
| *Enterococcus* | Firmicutes | −3.087 | 0.000 | 0.001 | S4 vs. S5 |
| *Enterococcus* | Firmicutes | −3.087 | 0.000 | 0.002 | S2 vs. S5 |
| | Acidobacteriota | −3.103 | 0.000 | 0.001 | S2 vs. S3 |
| | Acidobacteriota | −3.103 | 0.002 | 0.018 | S1 vs. S3 |
| *Mycobacterium* | Actinobacteriota | −3.137 | 0.000 | 0.000 | S2 vs. S3 |
| *Mycobacterium* | Actinobacteriota | −3.137 | 0.001 | 0.011 | S1 vs. S3 |
| *Kitasatospora* | Actinobacteriota | −3.170 | 0.000 | 0.000 | S4 vs. S5 |
| *Kitasatospora* | Actinobacteriota | −3.170 | 0.000 | 0.001 | S2 vs. S5 |
| *Bradyrhizobium* | Proteobacteria | −3.178 | 0.000 | 0.015 | S6 vs. S7a |
| *Bradyrhizobium* | Proteobacteria | −3.240 | 0.000 | 0.028 | S2 vs. S7a |
| *Sphingomonas* | Proteobacteria | −3.263 | 0.000 | 0.010 | S2 vs. S3 |
| *Escherichia-Shigella* | Proteobacteria | −3.392 | 0.000 | 0.000 | S4 vs. S5 |
| *Escherichia-Shigella* | Proteobacteria | −3.392 | 0.000 | 0.000 | S2 vs. S5 |
| *Enhydrobacter* | Proteobacteria | −3.405 | 0.001 | 0.011 | S1 vs. S3 |
| *Roseiarcus* | Proteobacteria | −3.458 | 0.000 | 0.000 | S2 vs. S3 |
| *Roseiarcus* | Proteobacteria | −3.458 | 0.001 | 0.009 | S1 vs. S3 |
| *Phenylobacterium* | Proteobacteria | −3.459 | 0.000 | 0.000 | S4 vs. S5 |
| *Phenylobacterium* | Proteobacteria | −3.459 | 0.000 | 0.000 | S2 vs. S5 |
| *Cutibacterium* | Actinobacteriota | −3.621 | 0.001 | 0.032 | S2 vs. S7a |
| *Candidatus* | Crenarchaeota | −3.644 | 0.000 | 0.000 | S4 vs. S5 |
| *Candidatus* | Crenarchaeota | −3.644 | 0.000 | 0.000 | S2 vs. S5 |

**Note:** Data shows genera with significant differential abundance across sites.

### Table 12.Significant Differential Abundance of Genera Across Site Comparisons

*Filtered for Adjusted p-value < 0.05*

| Genus | Phylum | **Log2 Fold Change** | p-value | **Adjusted p-value** | Site Comparison |
|---|---|---|---|---|---|
| *Cutibacterium* | Actinobacteriota | −3.648 | 0.001 | 0.044 | S2 vs. S4 |
| *Brevundimonas* | Proteobacteria | −3.700 | 0.000 | 0.003 | S4 vs. S5 |
| *Brevundimonas* | Proteobacteria | −3.700 | 0.000 | 0.005 | S2 vs. S5 |
| *Brevundimonas* | Proteobacteria | −3.701 | 0.001 | 0.035 | S1 vs. S5 |
| *Rickettsiella* | Proteobacteria | −3.787 | 0.000 | 0.001 | S2 vs. S3 |
| *Rickettsiella* | Proteobacteria | −3.787 | 0.001 | 0.012 | S1 vs. S3 |
| | WPS-2 | −3.866 | 0.000 | 0.000 | S2 vs. S3 |
| | WPS-2 | −3.866 | 0.001 | 0.009 | S1 vs. S3 |
| | Patescibacteria | −4.034 | 0.000 | 0.000 | S2 vs. S3 |
| | Patescibacteria | −4.034 | 0.000 | 0.003 | S1 vs. S3 |
| *Jatrophihabitans* | Actinobacteriota | −4.051 | 0.000 | 0.000 | S2 vs. S3 |
| *Jatrophihabitans* | Actinobacteriota | −4.052 | 0.000 | 0.003 | S1 vs. S3 |
| *Bacillus* | Firmicutes | −4.200 | 0.000 | 0.000 | S2 vs. S3 |
| *Bacillus* | Firmicutes | −4.200 | 0.000 | 0.008 | S1 vs. S3 |
| | Patescibacteria | −4.261 | 0.000 | 0.000 | S2 vs. S3 |
| | Patescibacteria | −4.261 | 0.000 | 0.002 | S1 vs. S3 |
| *Blattabacterium* | Bacteroidota | −4.404 | 0.000 | 0.000 | S2 vs. S3 |
| *Borrelia* | Spirochaetota | −4.598 | 0.000 | 0.008 | S2 vs. S5 |
| *Acinetobacter* | Proteobacteria | −4.687 | 0.000 | 0.000 | S2 vs. S3 |
| *Acinetobacter* | Proteobacteria | −4.688 | 0.000 | 0.001 | S1 vs. S3 |
| *Spiroplasma* | Firmicutes | −4.755 | 0.000 | 0.000 | S4 vs. S5 |
| *Spiroplasma* | Firmicutes | −4.755 | 0.000 | 0.000 | S2 vs. S5 |

**Note:** Data shows genera with significant differential abundance across sites.

### Table 12.Significant Differential Abundance of Genera Across Site Comparisons

*Filtered for Adjusted p-value < 0.05*

| Genus | Phylum | **Log2 Fold Change** | p-value | **Adjusted p-value** | Site Comparison |
|---|---|---|---|---|---|
| *Spiroplasma* | Firmicutes | −4.755 | 0.000 | 0.000 | S1 vs. S5 |
| *Spiroplasma* | Firmicutes | −4.755 | 0.000 | 0.000 | S3 vs. S5 |
| *Mycobacterium* | Actinobacteriota | −4.876 | 0.000 | 0.000 | S2 vs. S3 |
| *Mycobacterium* | Actinobacteriota | −4.876 | 0.000 | 0.001 | S1 vs. S3 |
| *Ac37b* | Proteobacteria | −5.039 | 0.000 | 0.005 | S6 vs. S7a |
| *Bradyrhizobium* | Proteobacteria | −5.049 | 0.000 | 0.000 | S2 vs. S3 |
| | Patescibacteria | −5.334 | 0.000 | 0.000 | S2 vs. S3 |
| | Patescibacteria | −5.335 | 0.000 | 0.000 | S1 vs. S3 |
| *Rickettsiella* | Proteobacteria | −5.375 | 0.000 | 0.000 | S4 vs. S5 |
| *Rickettsiella* | Proteobacteria | −5.375 | 0.000 | 0.000 | S2 vs. S5 |
| *Rickettsiella* | Proteobacteria | −5.375 | 0.000 | 0.000 | S3 vs. S5 |
| *Blattabacterium* | Bacteroidota | −5.485 | 0.000 | 0.000 | S2 vs. S3 |
| *Blattabacterium* | Bacteroidota | −5.486 | 0.000 | 0.000 | S1 vs. S3 |
| *Bradyrhizobium* | Proteobacteria | −5.535 | 0.000 | 0.003 | S1 vs. S3 |
| *Ac37b* | Proteobacteria | −5.577 | 0.001 | 0.017 | S1 vs. S4 |
| *Borrelia* | Spirochaetota | −5.655 | 0.000 | 0.000 | S4 vs. S5 |
| *Borrelia* | Spirochaetota | −6.277 | 0.000 | 0.000 | S3 vs. S5 |
| *Ac37b* | Proteobacteria | −6.386 | 0.000 | 0.003 | S2 vs. S3 |
| *Enterococcus* | Firmicutes | −6.559 | 0.000 | 0.000 | S2 vs. S3 |
| *Enterococcus* | Firmicutes | −6.559 | 0.000 | 0.000 | S1 vs. S3 |
| *Spiroplasma* | Firmicutes | −6.931 | 0.000 | 0.000 | S5 vs. S8a |
| *Spiroplasma* | Firmicutes | −6.931 | 0.000 | 0.000 | S4 vs. S8a |

**Note:** Data shows genera with significant differential abundance across sites.

### Table 12.Significant Differential Abundance of Genera Across Site Comparisons

*Filtered for Adjusted p-value < 0.05*

| Genus | Phylum | **Log2 Fold Change** | p-value | **Adjusted p-value** | Site Comparison |
|---|---|---|---|---|---|
| *Spiroplasma* | Firmicutes | −6.931 | 0.000 | 0.000 | S2 vs. S8a |
| *Spiroplasma* | Firmicutes | −6.931 | 0.000 | 0.000 | S6 vs. S8a |
| *Spiroplasma* | Firmicutes | −6.931 | 0.000 | 0.000 | S1 vs. S8a |
| *Spiroplasma* | Firmicutes | −6.931 | 0.000 | 0.000 | S3 vs. S8a |
| *Spiroplasma* | Firmicutes | −6.933 | 0.000 | 0.000 | S7a vs. S8a |
| *Ac37b* | Proteobacteria | −7.590 | 0.000 | 0.000 | S2 vs. S7a |
| *Spiroplasma* | Firmicutes | −7.972 | 0.000 | 0.000 | S5 vs. S8a |
| *Spiroplasma* | Firmicutes | −7.972 | 0.000 | 0.000 | S4 vs. S8a |
| *Spiroplasma* | Firmicutes | −7.973 | 0.000 | 0.000 | S2 vs. S8a |
| *Spiroplasma* | Firmicutes | −7.973 | 0.000 | 0.000 | S6 vs. S8a |
| *Spiroplasma* | Firmicutes | −7.973 | 0.000 | 0.000 | S1 vs. S8a |
| *Spiroplasma* | Firmicutes | −7.973 | 0.000 | 0.000 | S3 vs. S8a |
| *Spiroplasma* | Firmicutes | −7.974 | 0.000 | 0.000 | S7a vs. S8a |
| *Spiroplasma* | Firmicutes | −8.087 | 0.000 | 0.000 | S5 vs. S8a |
| *Spiroplasma* | Firmicutes | −8.087 | 0.000 | 0.000 | S4 vs. S8a |
| *Spiroplasma* | Firmicutes | −8.087 | 0.000 | 0.000 | S2 vs. S8a |
| *Spiroplasma* | Firmicutes | −8.088 | 0.000 | 0.000 | S6 vs. S8a |
| *Spiroplasma* | Firmicutes | −8.088 | 0.000 | 0.000 | S1 vs. S8a |
| *Spiroplasma* | Firmicutes | −8.088 | 0.000 | 0.000 | S3 vs. S8a |
| *Spiroplasma* | Firmicutes | −8.089 | 0.000 | 0.000 | S7a vs. S8a |
| *Spiroplasma* | Firmicutes | −8.522 | 0.000 | 0.000 | S2 vs. S8a |
| *Spiroplasma* | Firmicutes | −8.522 | 0.000 | 0.000 | S4 vs. S8a |

**Note:** Data shows genera with significant differential abundance across sites.

**Table 12.Significant Differential Abundance of Genera Across Site Comparisons**

*Filtered for Adjusted p-value < 0.05*

| Genus | Phylum | **Log2 Fold Change** | p-value | **Adjusted p-value** | Site Comparison |
|---|---|---|---|---|---|
| *Spiroplasma* | Firmicutes | −8.522 | 0.000 | 0.000 | S1 vs. S8a |
| *Spiroplasma* | Firmicutes | −8.522 | 0.000 | 0.000 | S5 vs. S8a |
| *Spiroplasma* | Firmicutes | −8.522 | 0.000 | 0.000 | S6 vs. S8a |
| *Spiroplasma* | Firmicutes | −8.523 | 0.000 | 0.000 | S3 vs. S8a |
| *Spiroplasma* | Firmicutes | −8.524 | 0.000 | 0.000 | S7a vs. S8a |
| *Ac37b* | Proteobacteria | −8.571 | 0.000 | 0.000 | S2 vs. S4 |
| *Spiroplasma* | Firmicutes | −8.921 | 0.000 | 0.000 | S2 vs. S8a |
| *Spiroplasma* | Firmicutes | −8.921 | 0.000 | 0.000 | S5 vs. S8a |
| *Spiroplasma* | Firmicutes | −8.921 | 0.000 | 0.000 | S4 vs. S8a |
| *Spiroplasma* | Firmicutes | −8.921 | 0.000 | 0.000 | S6 vs. S8a |
| *Spiroplasma* | Firmicutes | −8.921 | 0.000 | 0.000 | S3 vs. S8a |
| *Spiroplasma* | Firmicutes | −8.921 | 0.000 | 0.000 | S1 vs. S8a |
| *Spiroplasma* | Firmicutes | −8.923 | 0.000 | 0.000 | S7a vs. S8a |
| *Spiroplasma* | Firmicutes | −8.935 | 0.000 | 0.000 | S4 vs. S8a |
| *Spiroplasma* | Firmicutes | −8.936 | 0.000 | 0.000 | S5 vs. S8a |
| *Spiroplasma* | Firmicutes | −8.936 | 0.000 | 0.000 | S3 vs. S8a |
| *Spiroplasma* | Firmicutes | −8.936 | 0.000 | 0.000 | S2 vs. S8a |
| *Spiroplasma* | Firmicutes | −8.936 | 0.000 | 0.000 | S6 vs. S8a |
| *Spiroplasma* | Firmicutes | −8.936 | 0.000 | 0.000 | S1 vs. S8a |
| *Spiroplasma* | Firmicutes | −8.938 | 0.000 | 0.000 | S7a vs. S8a |
| *Spiroplasma* | Firmicutes | −9.494 | 0.000 | 0.000 | S2 vs. S8a |
| *Spiroplasma* | Firmicutes | −9.494 | 0.000 | 0.000 | S5 vs. S8a |

**Note:** Data shows genera with significant differential abundance across sites.

### Table 12. Significant Differential Abundance of Genera Across Site Comparisons

*Filtered for Adjusted p-value < 0.05*

| Genus | Phylum | **Log2 Fold Change** | p-value | **Adjusted p-value** | Site Comparison |
|---|---|---|---|---|---|
| *Spiroplasma* | Firmicutes | −9.494 | 0.000 | 0.000 | S1 vs. S8a |
| *Spiroplasma* | Firmicutes | −9.494 | 0.000 | 0.000 | S4 vs. S8a |
| *Spiroplasma* | Firmicutes | −9.494 | 0.000 | 0.000 | S6 vs. S8a |
| *Spiroplasma* | Firmicutes | −9.494 | 0.000 | 0.000 | S3 vs. S8a |
| *Spiroplasma* | Firmicutes | −9.495 | 0.000 | 0.000 | S7a vs. S8a |
| *Spiroplasma* | Firmicutes | −9.712 | 0.000 | 0.000 | S5 vs. S8a |
| *Spiroplasma* | Firmicutes | −9.712 | 0.000 | 0.000 | S4 vs. S8a |
| *Spiroplasma* | Firmicutes | −9.712 | 0.000 | 0.000 | S2 vs. S8a |
| *Spiroplasma* | Firmicutes | −9.712 | 0.000 | 0.000 | S6 vs. S8a |
| *Spiroplasma* | Firmicutes | −9.713 | 0.000 | 0.000 | S3 vs. S8a |
| *Spiroplasma* | Firmicutes | −9.713 | 0.000 | 0.000 | S1 vs. S8a |
| *Spiroplasma* | Firmicutes | −9.714 | 0.000 | 0.000 | S7a vs. S8a |
| *Spiroplasma* | Firmicutes | −10.058 | 0.000 | 0.000 | S2 vs. S8a |
| *Spiroplasma* | Firmicutes | −10.058 | 0.000 | 0.000 | S4 vs. S8a |
| *Spiroplasma* | Firmicutes | −10.058 | 0.000 | 0.000 | S5 vs. S8a |
| *Spiroplasma* | Firmicutes | −10.058 | 0.000 | 0.000 | S6 vs. S8a |
| *Spiroplasma* | Firmicutes | −10.058 | 0.000 | 0.000 | S3 vs. S8a |
| *Spiroplasma* | Firmicutes | −10.058 | 0.000 | 0.000 | S1 vs. S8a |
| *Spiroplasma* | Firmicutes | −10.060 | 0.000 | 0.000 | S7a vs. S8a |
| *Spiroplasma* | Firmicutes | −10.252 | 0.000 | 0.000 | S5 vs. S8a |
| *Spiroplasma* | Firmicutes | −10.252 | 0.000 | 0.000 | S2 vs. S8a |
| *Spiroplasma* | Firmicutes | −10.252 | 0.000 | 0.000 | S4 vs. S8a |

**Note:** Data shows genera with significant differential abundance across sites.

### Table 12.Significant Differential Abundance of Genera Across Site Comparisons

*Filtered for Adjusted p-value < 0.05*

| Genus | Phylum | **Log2 Fold Change** | p-value | **Adjusted p-value** | Site Comparison |
|---|---|---|---|---|---|
| *Spiroplasma* | Firmicutes | −10.252 | 0.000 | 0.000 | S6 vs. S8a |
| *Spiroplasma* | Firmicutes | −10.252 | 0.000 | 0.000 | S1 vs. S8a |
| *Spiroplasma* | Firmicutes | −10.252 | 0.000 | 0.000 | S3 vs. S8a |
| *Spiroplasma* | Firmicutes | −10.254 | 0.000 | 0.000 | S7a vs. S8a |

**Note:** Data shows genera with significant differential abundance across sites.
